# Why woodiness repeatedly evolves—and disappears

**DOI:** 10.64898/2026.09.04.749323

**Authors:** Kasper P. Hendriks, Ihsan A. Al-Shehbaz, Dmitry A. German, Marcus A. Koch, Lydia van Slooten, Carlijn Kusse, Lisa J.M.A. Dominicus, Marit Kuijt, Lila E. Trepp, Sander van Zon, Elena Castillo-Lorenzo, Barış Özüdoğru, Andreas Berger, Michael D. Windham, Leopoldo Medina Domingo, Nora Walden, Christiane Kiefer, David S. Aukes, Darío J. Schiavinato, Edie Burns, C. Donovan Bailey, Raquel Negrão, Stanislav Španiel, Alexey P. Seregin, Hamid Moazzeni, Óscar Toro-Núñez, Vanessa R. Invernón, Atena Eslami-Farouji, Peter Heenan, Terezie Mandáková, Hongliang Chen, Martin A. Lysak, Natalia M. Shiyan, Nikolai M. Hay, Mats Thulin, Robert Vogt, Mathieu Chambouleyron, Steven B. Janssens, Marie Briggs, Michaela Schmull, Alan Forrest, Hang Sun, Alessia Guggisberg, Fred W. Stauffer, Pieter J. Winter, M. Eric Schranz, Lachezar A. Nikolov, Alexandre R. Zuntini, William J. Baker, Félix Forest, Olivier Maurin, Klaus Mummenhoff, Frederic Lens

## Abstract

Derived woodiness—the evolution of woody growth from herbaceous ancestors—has arisen hundreds of times across flowering plants, yet the environmental conditions associated with its repeated evolution remain poorly understood. Here, we analyse woodiness evolution in the mustard family (Brassicaceae; ∼4,150 species) using a time-calibrated phylogeny of 2,927 species, including 374 of the 385 known woody species, together with global growth-form and climatic niche data. We infer 231 independent origins of woodiness alongside 176 reversals to herbaceousness, indicating that woodiness evolves repeatedly but remains evolutionarily unstable. Although woody species are enriched on islands, most occur on the mainland, where woodiness is consistently associated with drought and reduced frost. Correlated-evolution analyses reveal that drought is associated primarily with the persistence of woodiness, whereas reduced frost is associated with gains of woodiness and increased frost with its loss. These findings identify distinct climatic associations with the gain, persistence, and loss of woody growth forms.

## INTRODUCTION

Convergent evolution is widespread in plants, with similar traits arising repeatedly in response to environmental conditions. In angiosperms, growth form exemplifies one of the most striking cases of such repeated evolution. Woody species, characterised by pronounced secondary growth produced by a wood-forming vascular cambium in stems and roots, represent the ancestral condition in angiosperms, with multiple evolutionary reductions of secondary growth giving rise to herbaceous lineages (*1*, *2*). Yet woodiness has evolved repeatedly from herbaceous lineages, giving rise to hundreds of independently derived woody lineages across angiosperms (*3*, *4*). Why transitions between woody and herbaceous growth forms have occurred so frequently remains debated.

Derived woodiness is especially conspicuous on oceanic islands—a phenomenon first recognised by Darwin (*5*) and later formalised by Carlquist (*6*)—where woody species can constitute 10–25% of angiosperm diversity (*7*). Much later, however, it was found that most derived woody species actually occur on the mainland, particularly in open, harsh continental environments with recurrent drought cycles, as well as alpine systems (*4*). This broader distribution suggests that the drivers of woodiness extend beyond those of classical island syndromes. Proposed explanations include competition for light and variables that increase plant longevity, such as climatic stability and release from herbivory. These ideas were later complemented by the drought hypothesis, which proposes that drought could have been an important driver of woodiness transitions over evolutionary time and across broad geographic scales (*7*, *8*).

Progress in understanding woodiness has been limited by the lack of densely sampled lineage-specific analyses integrating phylogeny, woody versus herbaceous growth form, and climate at the global scale. Most studies have focused either on small clades (*9–11*) or on broad comparative datasets with limited representation of derived woody species and their closest non-woody relatives (*12*), reducing both statistical power and the ability to evaluate whether proposed drivers of woodiness gains and losses are consistently associated with growth-form transitions. The mustard family (Brassicaceae), comprising approximately 4,150 accepted species distributed across nearly all major island and continental environments and characterised by an unusually high number of independent transitions between herbaceous and woody growth forms, provides a uniquely powerful system to address this gap (*13*). Notably, the family captures approximately one out of every seven inferred angiosperm transitions to derived woodiness based on a global literature review (*4*), making it one of the most powerful systems for studying repeated growth-form evolution. Moreover, Brassicaceae span extreme environmental gradients, from Mediterranean shrublands to arid deserts and frost-prone mountains, allowing direct evaluation of ecological drivers of repeated woodiness gains and losses at a global scale.

Here, we combine a time-calibrated phylogeny comprising 2,927 Brassicaceae species, including 374 of the 385 known woody species and more than 2,500 herbaceous relatives, with species-level growth-form and climatic niche data to ask how environmental conditions are associated with the repeated gain, persistence, and loss of woodiness over evolutionary time. We show that woodiness evolved repeatedly but remains evolutionarily unstable, exhibiting substantially shorter persistence times than herbaceousness. Incorporating climatic context reveals that drought and frost are associated with distinct aspects of woodiness evolution: drought is associated primarily with the persistence of woodiness, whereas reduced frost is associated primarily with gains of woodiness. Together, these results provide a family-wide test of major hypotheses for woodiness evolution and demonstrate how environmental conditions are associated with the origin and persistence of plant growth forms through evolutionary time.

## RESULTS

### Woodiness is repeatedly gained and lost

Across ∼4,150 Brassicaceae species, the 385 woody species are unevenly distributed, being concentrated in arid and seasonally dry regions, mountains and oceanic islands (Fig. 1).

**Fig. 1.**
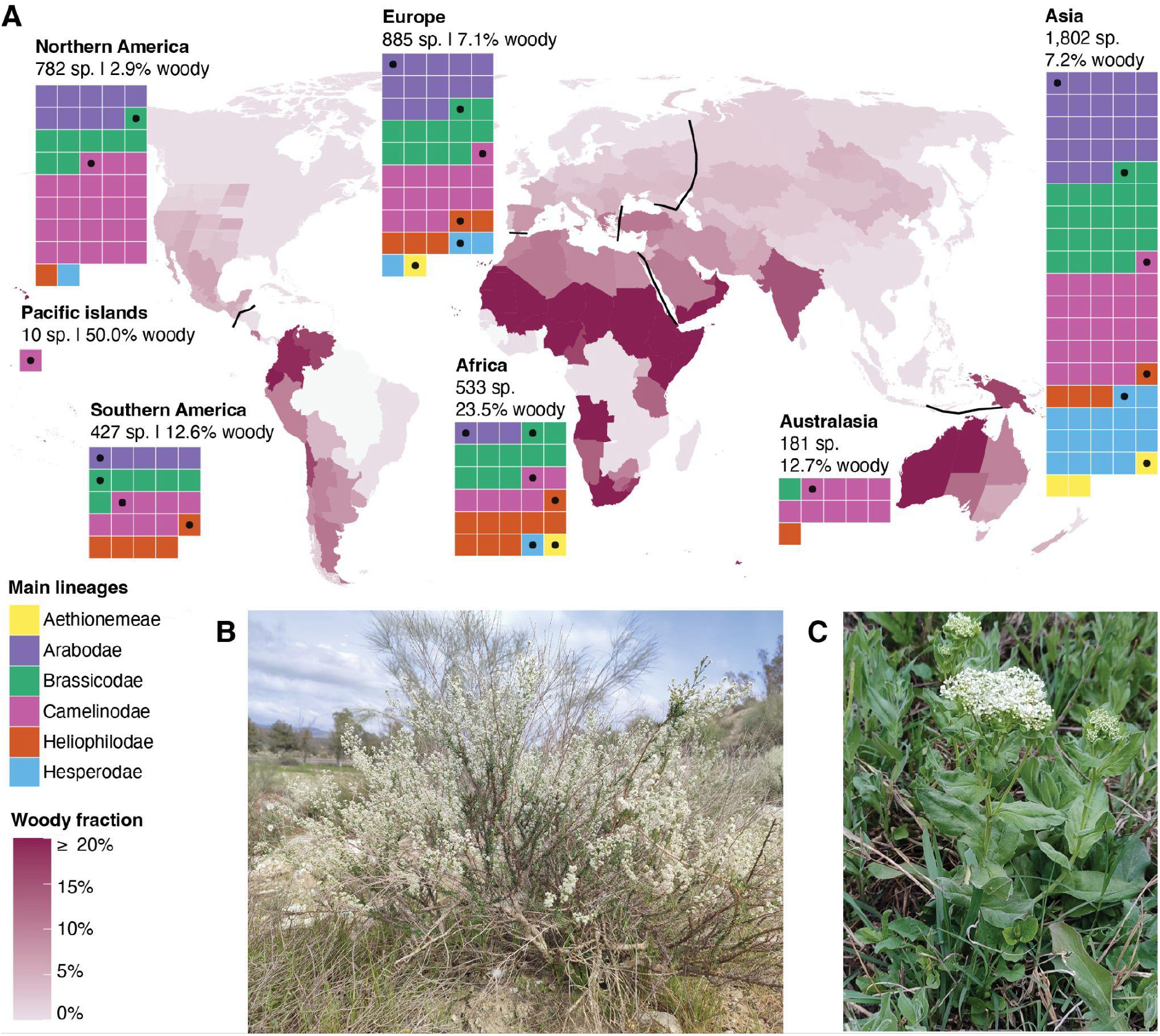
Woodiness across the Brassicaceae (A) Global variation in the proportion of woody Brassicaceae species. Purple shading indicates the proportion of woody species within each botanical country. Waffle plots summarise continental species richness and lineage composition, with each tile representing 20 species (rounded up) coloured according to the six main Brassicaceae lineages. Black dots indicate bins containing at least one woody species. (B–C) Contrasting growth forms within closely-related Brassicaceae: the shrub *Lepidium subulatum* (B) and herbaceous *Lepidium draba* (C). Photographs © the first author.

Although woody species occur across all main evolutionary lineages and continents, they are taxonomically clustered: 80% occur in just eight of the family’s 58 tribes, while 25 tribes contain at least one woody species (hereafter referred to as “tribes with woody species”; Fig. S1). Thus, woodiness is geographically and taxonomically widespread but unevenly distributed across the family. Null-model analyses revealed significant enrichment of woody species in arid regions, including the Horn of Africa, Arabian Peninsula, western Australia and parts of the Andes, and significant depletion in high-latitude, frost-prone regions (Fig. S2 and Data file S1).

To place woodiness transitions in an evolutionary context, we reconstructed a new time-calibrated Brassicaceae phylogeny (Brassicaceae Tree of Life, or BrassiToL; Fig. 2, Figs. S3–S4 and the Supplementary Results; https://tol.naturalis.nl/collection/brassicaceae), comprising 2,927 Brassicaceae species covering all 58 tribes and 362 of ∼368 genera, including 374 of the 385 woody Brassicaceae species (Data file S2). The family originated during the Eocene, between 54.7 and 36.9 Ma (Table S1). Growth form exhibits a strong phylogenetic signal (Pagel’s λ = 0.945; Fritz & Purvis’ D = 0.202; Table S2). Ancestral state reconstruction using an all-rates-different (ARD) model showed that the family was ancestrally herbaceous, with 231 independent origins of woodiness and 176 reversals to herbaceousness (Tables S3–S4).

**Fig. 2.**
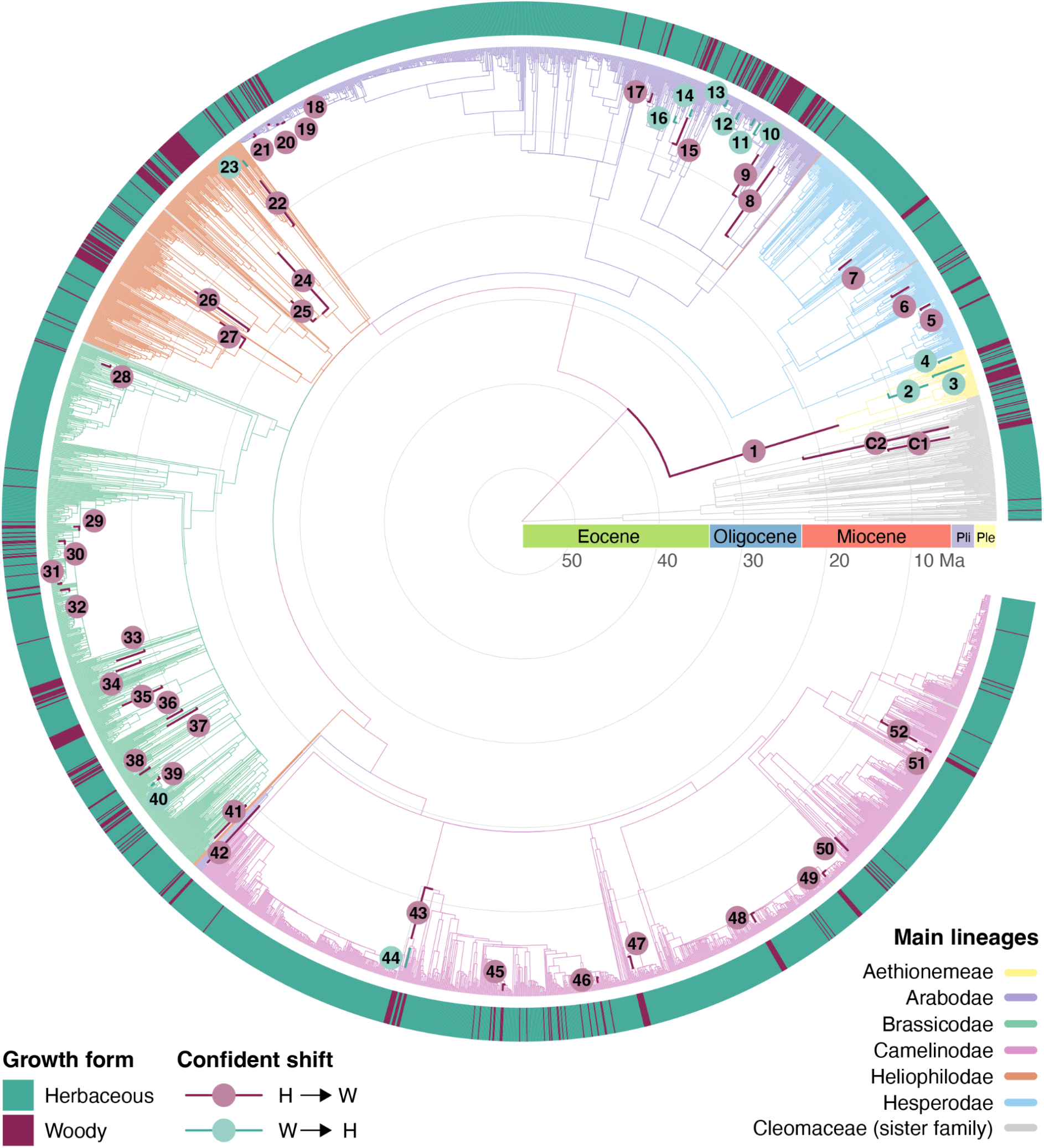
Repeated gains and losses of woodiness across the Brassicaceae phylogeny Time-calibrated Brassicaceae phylogeny inferred from 1,052 single-copy nuclear genes and comprising 2,927 species, including 374 of the 385 currently recognised woody species. Growth form is indicated in the outer ring, with woody species shown in purple and herbaceous species in turquoise. Confidently inferred growth-form transitions supported by both ancestral-state reconstruction and stochastic character mapping are indicated on branches. Across the phylogeny, we identified 42 internal and 101 terminal gains of woodiness (H→W), together with 12 internal and 63 terminal reversals to herbaceousness (W→H). Growth-form transitions occur throughout the evolutionary history of the Brassicaceae and across all six main lineages, highlighting the repeated and widespread evolution of both woody and herbaceous growth forms within the family. Numbered internal transitions correspond to the clades shown in Fig. 3A.

Restricting analyses to confident transitions between woody and herbaceous states, defined as shifts supported by both ancestral-state reconstruction and stochastic character mapping, identified 42 internal and 101 terminal transitions to woodiness, alongside 12 internal and 63 terminal reversals (Fig. 3A, Data file S3 and Tables S5–S6). These shifts occur across all main lineages and throughout the family’s evolutionary history, with no evidence for concentration in any particular time interval (Fig. 3B and Table S6). In all but one tribe with woody species, woodiness originated after tribe origin, with a mean lag time of 9.6 Myr (Fig. 3A and Table S7), and multiple independent origins within tribes were common (Table S6).

**Fig. 3.**
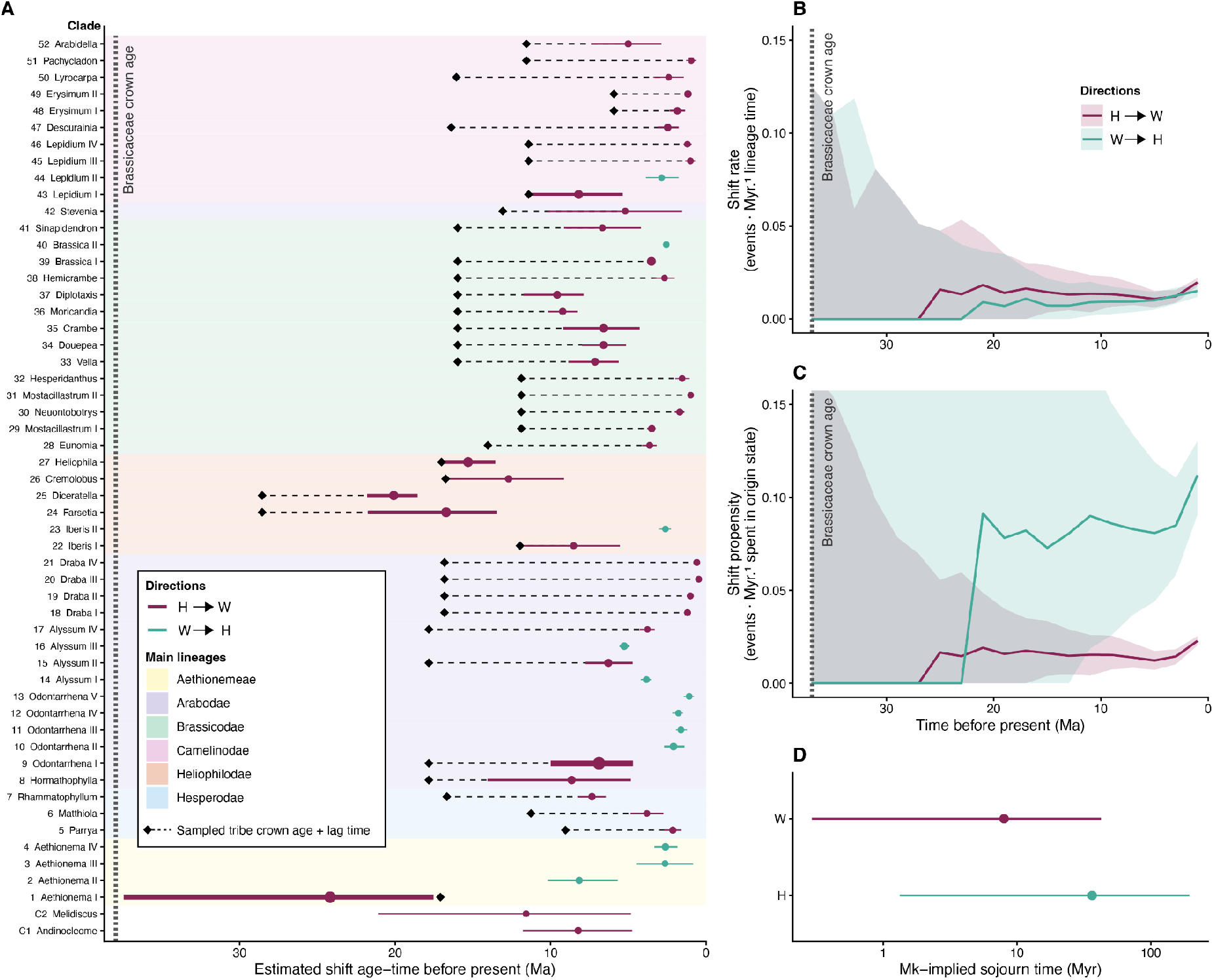
Timing and asymmetric dynamics reveal evolutionary instability of woodiness (A) Posterior age estimates for confident internal growth form shifts (herbaceous → woody; woody → herbaceous) for numbered clades highlighted in Fig. 2. Points show medians and error bars indicate 95% intervals. For H→W shifts, black diamonds indicate tribe crown node ages and hyphenated lines the lag time between tribe crown node origin and shift; background colour denotes the family’s main lineages. Line thickness is proportional to the number of descendant species represented by the corresponding clade in the phylogeny. (B) Similar transition rates through time for H→W and W→H (events per million years per unit lineage time). (C) Much higher shift propensity through time for W→H than for H→W (events per million years spent in the origin state). (D) Mk-implied sojourn time (expected state duration in million years) of herbaceous and woody states under the fitted continuous-time Markov model. In (A–C), the vertical dashed line marks the Brassicaceae crown node age. In (B–C), solid lines show medians and ribbons 95% intervals. Together, these results show that although woodiness evolves repeatedly, it is evolutionarily unstable and susceptible to reversion.

Despite frequent transitions, woodiness evolution is strongly asymmetric (Table S8). While transition rates inferred from stochastic character mapping are similar in both directions (herbaceous→woody and woody→herbaceous; Fig. 3B), shift propensity—accounting for time spent in each state—is four to six times higher towards herbaceousness (Fig. 3C). Consistently, Mk-implied persistence times (expected state durations under the fitted model) are much shorter for woody lineages (median 8.0 Myr) than for herbaceous lineages (36.4 Myr; Fig. 3D). Together, these results indicate that woodiness evolved repeatedly but remains evolutionarily unstable and prone to reversion.

### Woodiness is associated with drought and reduced frost

We tested environmental (drought, frost, heat) and geographical variables (island endemicity, elevation) associated with derived woodiness across three niche-data thresholds (ge1, ge5 and ge10, i.e. species occurrence in ≥1, ≥5 and ≥10 grid cells, respectively; Data files S4– S7 and Figs. S5–S6). Most Brassicaceae species are not island endemics: ∼85% of woody species and >98% of herbaceous species occur on the mainland or have broader distributions. Nevertheless, among island endemics (including both oceanic and continental islands), woodiness is disproportionately common, representing approximately half of all species (Table S9), and models including island endemicity recovered a strong positive association with woodiness (Fig. 4, Figs. S7–S8 and Tables S10–S16). We therefore repeated the analyses on a mainland-only subset to remove insular effects.

**Fig. 4.**
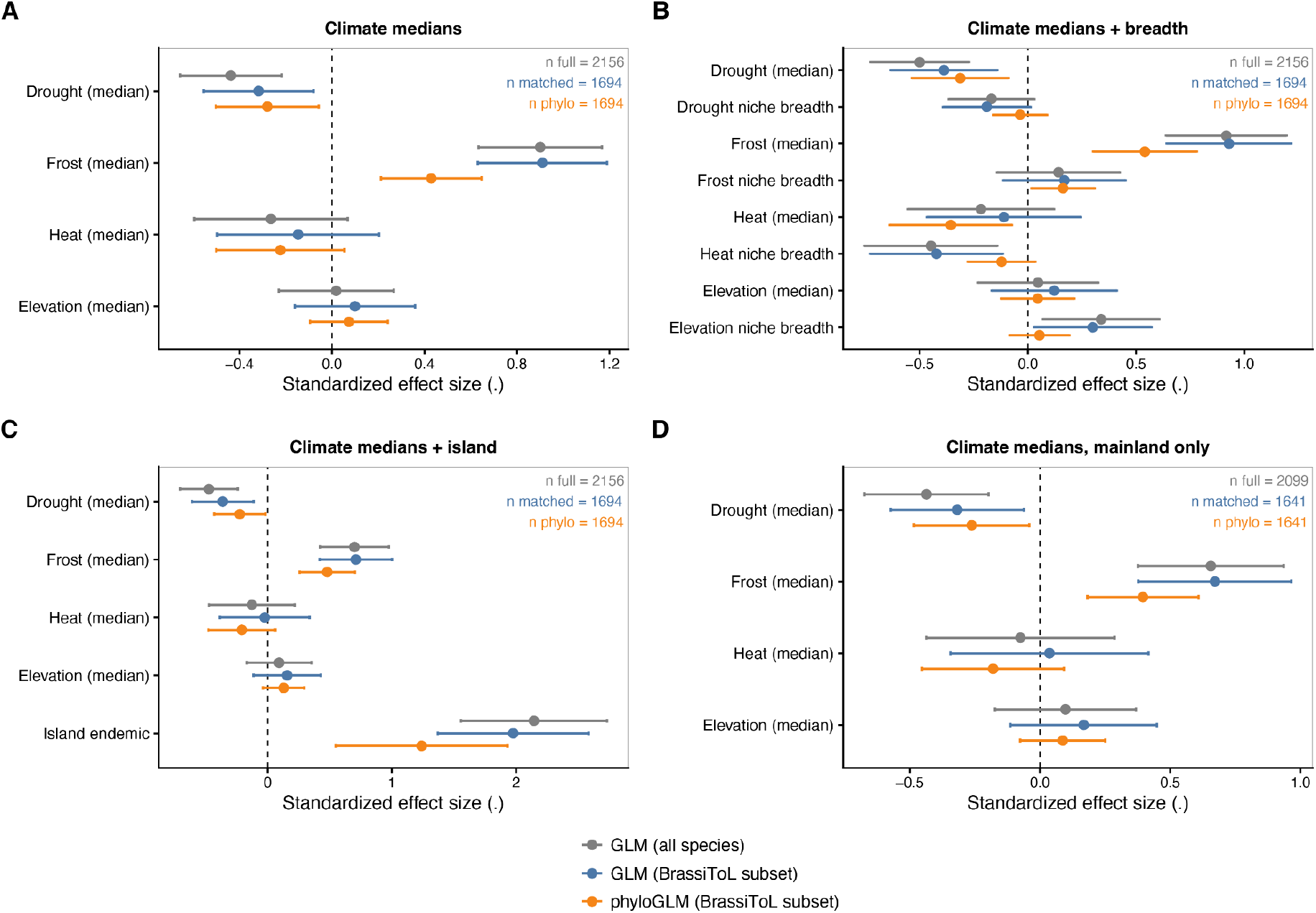
Drought and reduced frost are consistently associated with woodiness across analytical frameworks (A) Standardised effect sizes (β ± 95% Wald CI) for climatic niche median values in the ge10 dataset. (B) Equivalent analysis including both climatic niche median values and niche breadth estimates. (C) Equivalent analysis including climatic niche median values and island endemicity. (D) Equivalent analysis restricted to mainland species. Results are shown for generalised linear models (GLMs) fitted using all species, GLMs restricted to species present in the Brassicaceae Tree of Life (BrassiToL), and phylogenetic GLMs (phyloGLMs) fitted to the same BrassiToL subset. Species numbers for each model are indicated within panels. Predictors include drought (MCWD), frost (BIO6), heat (BIO10), elevation, their corresponding niche-breadth terms where applicable, and island endemicity. Across model formulations, drought and reduced frost show the strongest and most consistent associations with woodiness. Restricting analyses to species represented in the phylogeny has little effect on estimated coefficients, whereas phylogenetic correction generally reduces effect sizes, indicating that part of the observed climatic signal is explained by shared evolutionary history. Equivalent analyses for the ge1 and ge5 datasets are provided in Fig. S7.

Across all model formulations, drought (represented by Maximum Cumulative Water Deficit; MCWD) and frost (represented by the minimum temperature of the coldest month; BIO6) emerged as the primary correlates of woodiness, whereas elevation and heat effects (represented by mean temperature of the warmest quarter; BIO10) were weak and inconsistent (Fig. 4). In the ge10 dataset, model selection consistently favoured the mainland-only climate model (Fig. S10 and Table S13), indicating that climatic associations on the mainland explain woodiness patterns better than additional predictor complexity.

In all model frameworks, woodiness was associated with more severe drought and reduced frost: woody species occurred in regions with lower MCWD values (indicating stronger drought stress) and higher minimum temperatures of the coldest month (indicating reduced frost stress) (Fig. 5). Although drought and frost were moderately correlated (Spearman ρ ≈ −0.5), both retained independent effects when included in the same models (Tables S15–S16).

**Fig. 5.**
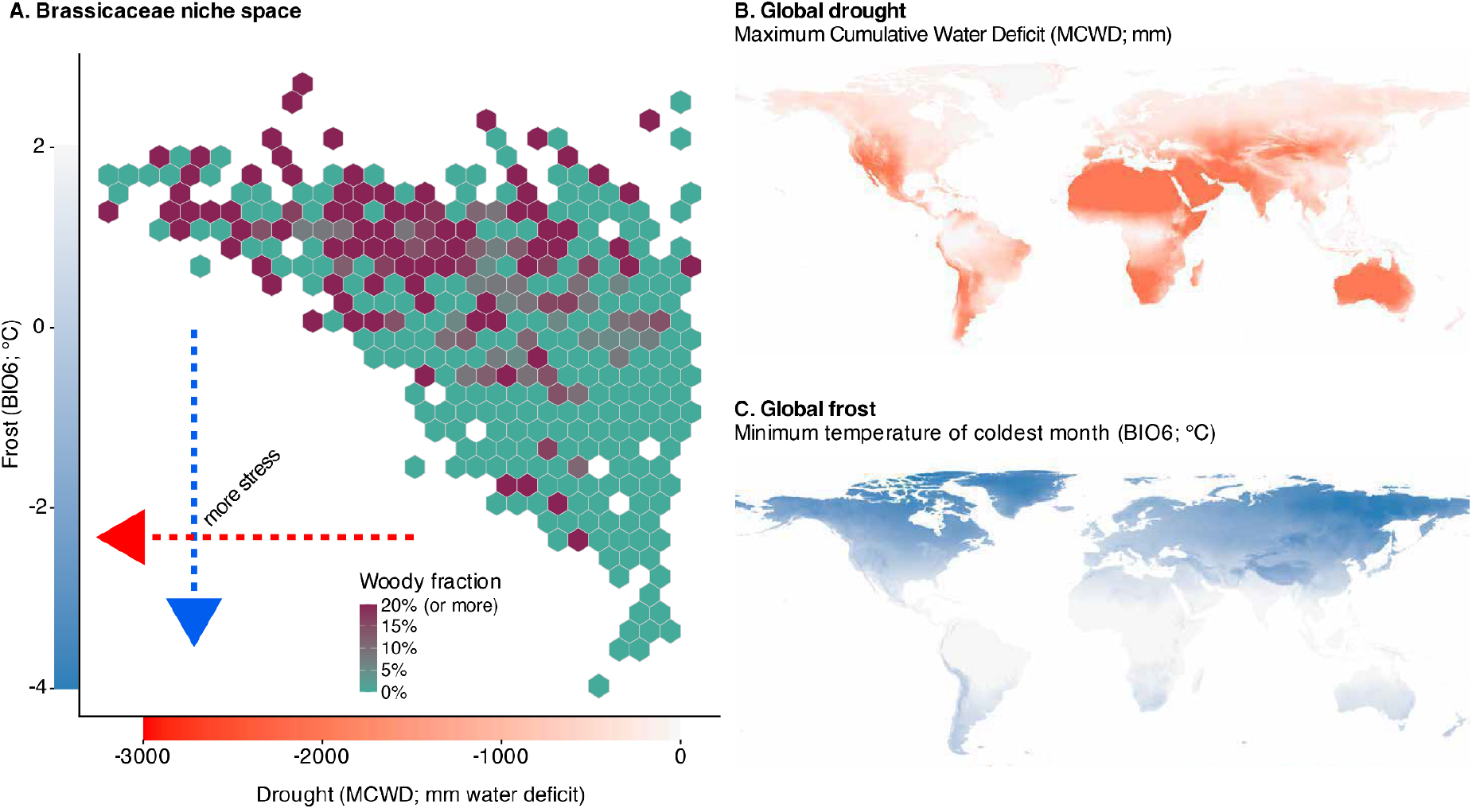
Woody Brassicaceae are concentrated in drought-prone, frost-free environments (A) Hexagon-binned climatic niche space of Brassicaceae species in the ge10 dataset, defined by species-level median drought (Maximum Cumulative Water Deficit; MCWD) and frost (minimum temperature of the coldest month; BIO6). Fill colour indicates the fraction of woody species per bin; the colour scale is capped at 20% woody species to enhance resolution across the observed range. Arrows indicate increasing drought and frost stress. Drought and frost were moderately correlated (Spearman ρ ≈ −0.5), but retained independent effects. (B) Global distribution of drought (MCWD), with more negative values indicating drier conditions (colour scale as in panel A). (C) Global distribution of frost (BIO6), with lower values indicating increased frost (colour scale as in panel A).

Accounting for phylogeny reduced effect sizes relative to non-phylogenetic models, indicating substantial phylogenetic structuring of the signal (phyloGLM α = 0.05–0.06; Fig. 4 and Table S10). Allowing supertribe-specific climate effects substantially improved model fit (ΔAIC = 69.4; likelihood-ratio test P < 10⁻¹³; Table S17), indicating ecological heterogeneity among main Brassicaceae lineages. Despite this heterogeneity, most supertribes retained the global association between woodiness, drought and reduced frost (Figs. S11–S12 and Tables S18–S19). The main exception was tribe Aethionemeae, a small and phylogenetically distinct lineage in which woodiness was associated with hotter summers and colder winters rather than the family-wide drought signal. Supertribe Arabodae showed a weaker association with drought and frost than the overall family trend, although these deviations were comparatively modest.

### Drought and frost influence woodiness through distinct evolutionary routes

To test whether growth form and environmental conditions evolved in a correlated manner, we fitted discrete-state models in BayesTraits using the ge10 dataset (Fig. 6A, Tables S20–S23 and Data file S8). Across all environmental thresholds (drought and frost percentile thresholds p40, p50 and p60, as well as the freezing threshold of 0 °C for frost), the dependent model was strongly favoured over the independent model, indicating correlated evolution between woodiness and both drought and frost (median logBF = 8–33 for drought and 12–33 for frost; Table S21).

**Fig. 6.**
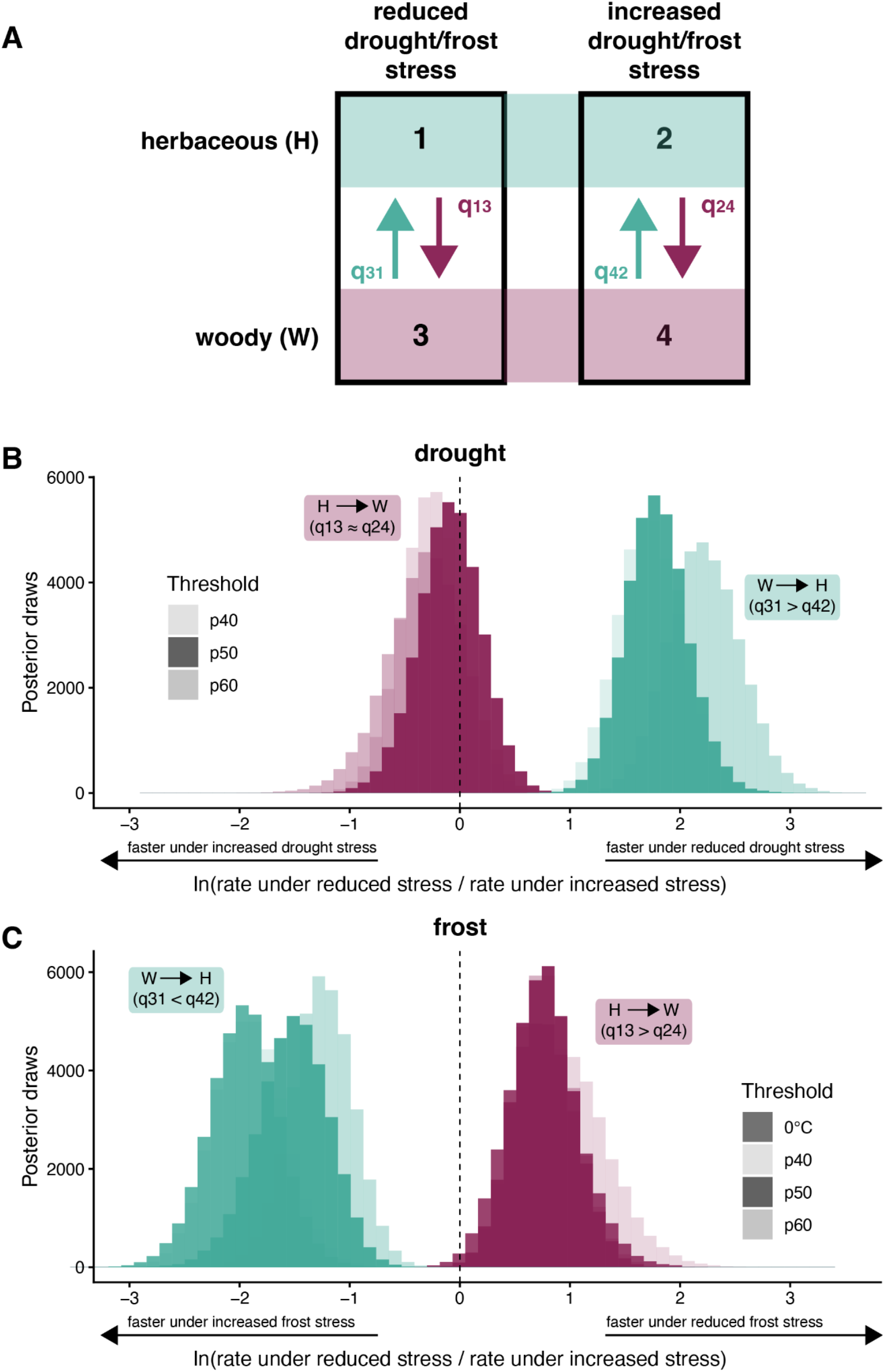
Drought and frost influence woodiness transitions through different evolutionary routes (A) Four-state BayesTraits model used to test correlated evolution between growth form and environmental stress. States combine herbaceous (H) and woody (W) growth forms with reduced versus increased drought/frost stress. Transition rates between growth forms are represented by q13, q24, q31 and q42. (B, C) Posterior distributions of natural-log-transformed transition-rate ratios estimated with BayesTraits for drought (B) and frost (C). Values are plotted as ln(rate under reduced stress/rate under increased stress), with negative values indicating faster transitions under increased stress and positive values indicating faster transitions under reduced stress. For drought, reversals to herbaceousness are substantially faster under reduced drought stress, whereas gains of woodiness show little dependence on drought stress. Under frost, woodiness evolves more rapidly under reduced stress, whereas reversals to herbaceousness are accelerated under increased stress. Darker histograms indicate median (p50) thresholds; lighter histograms indicate p40 and p60 thresholds. Frost analyses additionally include a mechanistic 0 °C threshold.

The direction of this correlation differed between variables. Under drought, woodiness was affected primarily through reduced reversal rates: woody-to-herbaceous transitions were substantially slower under strong drought, whereas gains of woodiness showed little consistent dependence on drought (Fig. 6B). This pattern was consistent across thresholds.

Frost showed a contrasting pattern (Fig. 6C). Transitions to woodiness were faster under reduced or absent frost, whereas reversals to herbaceousness were faster under frost. This pattern was recovered across percentile thresholds and the mechanistic freezing threshold of 0 °C (Table S23), indicating that reduced frost is associated with gains of woodiness, whereas increased frost is associated with its loss.

Thus, drought and frost are associated with distinct components of woodiness evolution: drought is associated primarily with the persistence of woodiness, whereas reduced frost is associated primarily with gains of woodiness.

## DISCUSSION

By integrating a time-calibrated phylogeny of 2,927 Brassicaceae species, including 374 of the 385 known woody species and more than 2,500 herbaceous relatives, with global growth-form and climatic niche data, we provide a comprehensive macroevolutionary analysis of growth-form transitions within a single plant family. Our results reveal an extraordinary degree of evolutionary lability, with woodiness originating independently approximately 231 times, more than doubling previous estimates within the family. The large number of inferred reversals (176), together with the substantially higher propensity for transitions back to herbaceousness, shows that derived woodiness remains evolutionarily unstable despite a strong phylogenetic signal in growth form. Although woody species are disproportionately enriched on islands, most woody Brassicaceae occur on the mainland, where woodiness is consistently associated with increased drought and reduced frost. Most importantly, drought and frost are associated with different aspects of woodiness evolution: drought is associated primarily with the persistence of woodiness, whereas reduced frost is associated primarily with gains of woodiness.

The >200 inferred origins of woodiness occur across all major Brassicaceae lineages and in environmental settings that harbour many derived woody species from other angiosperm families, including the Old-World Dry Belt, southern Africa, arid Australia and parts of the Andes (*4*) (Fig. 3 and Fig. S4). Together with the large number of reversals, this evolutionary lability suggests that the developmental capacity for secondary growth remains broadly accessible throughout much of the family (*14*). Experimental studies in *Arabidopsis thaliana* show that relatively small changes in the activity of developmental regulators can profoundly alter vascular cambial activity and longevity. For example, knocking out the flowering-time regulators SOC1 and FUL, or overexpressing the downstream rejuvenator gene AHL15, can shift annual herbaceous *Arabidopsis* towards a more perennial and woody growth form (*15*, *16*). These findings suggest that secondary growth remains an intrinsic developmental capacity throughout much of the family, and that repeated gains and losses of woodiness may often reflect changes in the regulation, timing or duration of its expression rather than the repeated evolution of entirely novel genetic machinery.

Our phylogenetic framework enables a more nuanced view of how drought and frost are associated with growth-form transitions. Rather than primarily promoting the origin of woodiness, drought is associated mainly with the persistence of woodiness through reduced rates of reversal, whereas reduced frost is associated primarily with gains of woodiness. This distinction helps reconcile two long-standing explanations for woodiness evolution. Reduced frost aligns closely with the favourable-climate hypothesis, as longer growing seasons and reduced environmental constraints may increase longevity and the opportunity for secondary growth to accumulate (*6*). Drought, by contrast, appears to influence the long-term retention of woody growth forms. Species with increased levels of secondary growth are often more resistant to drought-induced hydraulic failure than their herbaceous relatives because their conduits are embedded within thicker-walled, highly lignified tissues that help maintain hydraulic function under water limitation (*8*, *17*). Similar relationships have also been documented across herbaceous angiosperms, where increased stem lignification is associated with greater hydraulic safety and enhanced drought tolerance (*18–20*). Notably, many of the continental hotspots of derived woodiness identified across angiosperms coincide with regions that have experienced prolonged aridification through geological time (*4*). This concordance suggests that the environmental conditions associated with the persistence of woody lineages may themselves remain stable over evolutionary timescales. Together, these observations provide a plausible explanation for why drought is associated primarily with the persistence rather than the origin of woodiness.

The opposite trend towards herbaceousness, reflected by both accelerated reversals to herbaceousness and reduced gains of woodiness, is associated with increased frost. This pattern is consistent with long-standing hypotheses linking woodiness to climatically favourable, frost-free environments, particularly on oceanic islands (*4*, *6*, *7*). Freezing environments impose additional functional requirements on woody plants, and successful occupation of frost-prone regions often requires specialised hydraulic strategies, including xylem architectures with narrower conduits and, in many groups, seasonal leaf shedding to avoid freeze–thaw damage (*21*). Herbaceous species can instead avoid many of these constraints through short life spans or seasonal senescence of comparatively inexpensive above-ground tissues (*21*). Our results are therefore consistent with the idea that frost shifts the balance towards herbaceous life histories, whereas frost-free environments facilitate the repeated evolution of woody lineages.

These findings place the classical phenomenon of island woodiness within a broader evolutionary context. Consistent with recent angiosperm-wide analyses (*4*), woody Brassicaceae are enriched on islands but occur predominantly in continental open habitats with seasonal drought, including steppes, deserts, rocky slopes and wastelands. Rather than representing a distinct evolutionary phenomenon, island woodiness may reflect one end of a broader environmental continuum in which woody species are more abundant under conditions that favour either the evolutionary origin (through reduced frost) or the persistence (through drought) of the woody growth form. By showing that these associations recur within a single densely sampled family and by distinguishing gains, persistence and loss, our study extends previous work from geographic patterns of derived woodiness to the evolutionary processes shaping them.

More broadly, the Brassicaceae Tree of Life established here provides a foundation for investigating the genomic basis of secondary growth beyond *Arabidopsis*, the family’s classic developmental model. By combining phylogenomics, macroecology and comparative developmental biology, our study demonstrates how environmental conditions are associated with both the origin and persistence of complex traits across deep evolutionary time, while establishing a powerful framework for testing the mechanisms underlying repeated evolutionary transitions.

## METHODS

### Brassicaceae Tree of Life reconstruction

To reconstruct the Brassicaceae Tree of Life (BrassiToL), we largely followed the methods of (*22*) and the latest taxonomic standards in the family (*23*, *24*). We assembled target-capture sequencing data for 2,927 Brassicaceae species, 135 Cleomaceae species, and 23 species representing the remaining 16 families of Brassicales. The final dataset comprised 3,093 samples, including 2,140 newly generated for this study, and 953 obtained from previous studies (*22*, *25–29*) and additional datasets available through the NCBI Sequence Read Archive (SRA) and European Nucleotide Archive (ENA) (Data file S2).

For the 2,140 newly sequenced samples, genomic DNA was extracted from approximately 25 mg of dry herbarium tissue using the DNeasy Plant Pro Kit (Qiagen) and archived in the Naturalis DNA Bank (Data file S2). Samples with fragment peaks >400 bp were sheared to ∼300 bp using a Covaris M220 ultrasonicator. Libraries were prepared at quarter-volume using the NEBNext Ultra II FS Kit (New England Biolabs) and pooled (20–40 libraries per pool) for target capture of 1,081 single-copy nuclear genes using a combination of the Angiosperms353 bait set (*30*) and Brassicaceae764 bait set (*26*) (36 loci overlapping (*31*)). Sequencing was performed on an Illumina NovaSeq 6000 platform (BaseClear), generating 150-bp paired-end reads at approximately 100× target coverage.

Sequence processing followed a custom bioinformatic pipeline. Reads were mapped to target sequences using HybPiper v2.1.2 (*32*), and consensus sequences were generated using custom scripts adapted from HybPhaser v2.1 (*33*). Genes flagged as potential paralogs—either by HybPiper or as SNP outliers following HybPhaser—were excluded, yielding 1,052 nuclear loci (Figs. S13–S14 and Data file S9).

Codon-aware alignments were generated with OMM-MACSE (*34*) and trimmed using trimAl v1.2 (*35*). Maximum likelihood gene trees were inferred with IQ-TREE v3.0.1 (*36*) using ultrafast bootstrap (*37*) and SH-aLRT support (*38*). Nodes with low support (UFBoot <80, SH-aLRT <70) or extremely short branches (<1 × 10⁻⁵) were collapsed prior to coalescent-based species tree inference in ASTRAL-IV (*39*).

Divergence times were estimated on the fixed ASTRAL species-tree topology using Least-Squares Dating v.2 (*40*) (LSD2). To select loci suitable for time calibration, candidate genes were first ranked using SortaDate (*41*) based on taxon coverage and clock-likeness, followed by custom coverage-aware filtering to maximise representation across the phylogeny. A subset of the 200 top-ranked loci was then used to estimate partitioned branch lengths in IQ-TREE v.3 under a fixed topology, after which divergence times were inferred in LSD2 using a relaxed-clock framework. The Brassicales root was constrained to 90 Ma using *Macrohasseltia macroterantha* (Malvaceae) as outgroup taxon, corresponding to the estimated stem age of Brassicales based on the fossil *Dressiantha bicarpellata* (93.6–89.3 Ma (*42*)). Confidence intervals were estimated using 100 parametric replicates.

### Woodiness evolution

Growth-form scores were compiled from expert knowledge of Brassicaceae specialists, informed by floristic and taxonomic literature. Species were classified as woody only when they normally produce a continuous wood cylinder extending into the upper stem. Herbs, suffrutescent taxa and species with secondary growth restricted to the stem base were classified as herbaceous, following criteria applied in previous studies of derived woodiness (*7*) (Data file S7).

Species’ native distributions were summarised by botanical country following the World Geographical Scheme for Recording Plant Distributions (WGSRPD) level-3 regions distribution data from WCVP: World Checklist of Vascular Plants (*43*). For each botanical country, we calculated species richness and the proportion of woody species. Countries with significantly higher or lower numbers of woody species than expected given their Brassicaceae richness were identified using a null-model randomisation in which woody growth form was randomly assigned across the global species pool while preserving the observed number of woody species (1,000 replicates). Standardised effect sizes (SES) and empirical P-values were calculated by comparing observed woody-species richness to the resulting null distributions (Data file S1).

Phylogenetic signal was quantified using Pagel’s λ (*44*) and Fritz and Purvis’ D (*45*) (Table S2). Ancestral states were reconstructed under Mk models with equal rates (ER) and all rates different (ARD), selecting the best-fitting model using the Akaike information criterion (AIC; Table S3). Because the ancestor of Brassicales is inferred to be woody (*13*), analyses were conducted with a woody root prior; results were robust to alternative priors.

To estimate the number, timing and direction of transitions, we performed stochastic character mapping (SIMMAP (*46*)) under the best-fitting model (ARD), summarising 500 posterior maps (Table S4). Transitions were classified as confident when ancestral-state reconstruction and stochastic mapping supported the same directional change along a branch (Tables S5–S7 and Data file S3). For inferred shifts to woodiness, we additionally calculated lag times relative to the origin of the corresponding tribe (Table S7).

Temporal dynamics were characterised by estimating (i) transition rates through time, (ii) state-specific shift propensity (transition rates conditioned on time spent in each state; Table S8), and (iii) Mk-implied sojourn times (expected time spent in a state before transition). All analyses were conducted in R.

### Niche estimation

Climatic niches were characterised using 10,298,458 occurrence records across 4,102 Brassicaceae species obtained from GBIF (GBIF.org 2025; https://doi.org/10.15468/dd.x9aqbz), supplemented with 290 expert records for 46 rare species (Data file S4). Analyses were restricted to accepted species and excluded infraspecific taxa.

Records were filtered using a custom automated pipeline, retaining only occurrences within native ranges, defined using WCVP botanical countries with a 100-km buffer, and applying standard coordinate-cleaning procedures (including CoordinateCleaner (*47*)), including the removal of records located in cities, biodiversity institutions and oceans. For species with insufficient coordinates (target ≥20 records), locality descriptions contained in GBIF metadata were automatically parsed and standardised using the OpenAI language model API within the pipeline, after which locations were georeferenced through the Google Maps API. Geographic precision estimates returned by Google Maps were translated into coordinate-uncertainty values and incorporated into the same filtering pipeline. Georeferencing was attempted for 2,007 species, with final contributions to 237 species after filtering.

Additional manual validation was performed for 29,017 records across 699 taxonomically challenging species, such as species complexes, frequently misidentified taxa and species with geographically unexpected occurrence records, using a custom Shiny application involving 17 taxonomic experts (Data file S5). For each species, up to 100 records were selected to maximise geographic coverage while prioritising directly georeferenced records and low coordinate uncertainty.

Records were projected onto a 30 arc-second grid and thinned to one record per cell. Environmental and elevation data were extracted from WorldClim v2 (*48*) and the Zomer v3 evapotranspiration dataset (*49*). Species-level summaries were calculated for Maximum Cumulative Water Deficit (MCWD; representing drought), minimum temperature of the coldest month (BIO6; frost), mean temperature of the warmest quarter (BIO10; heat), and elevation (Data file S6).

Species-level niche summaries were calculated as the median and median absolute deviation (MAD) across retained cells, representing central tendency and niche breadth. Because sampling depth varied, datasets were constructed for species represented by ≥1, ≥5 and ≥10 grid cells (ge1, ge5 and ge10; where “ge” denotes the number of retained grid cells per species; Data file S7). Across 4,123 species, this yielded 3,415 (ge1), 2,602 (ge5) and 2,159 species (ge10).

### Woodiness–niche associations

To test how climatic niche variation was associated with woodiness, we fitted logistic regression models with woodiness as a binary response variable. Four predictor sets were compared: a climate-only model including median values for MCWD, BIO6, BIO10 and elevation (m_clim); an extended model including niche breadth (median absolute deviation; MAD) for these variables (m_clim_breadth); a model including climatic predictors and island endemicity (m_clim_island); and a mainland-only model excluding island endemics (m_clim_mainland) (Table S10).

Each model set was fitted in three frameworks: (i) generalised linear models (GLMs) using all species, (ii) GLMs restricted to species present in the BrassiToL, and (iii) phylogenetic GLMs (phyloGLMs) fitted to this matched subset (Table S10). Dataset restriction had minor effects on coefficients (|Δβ| ≈ 0–0.2; Table S11), whereas phylogenetic correction had moderate effects (|Δβ| ≈ 0–0.5; Table S12). Model fit was assessed using AIC (Table S13). McFadden’s pseudo-R² values were modest (R² ≈ 0.07–0.14) and consistently lower in phyloGLMs, reflecting that part of the explanatory variation is captured by phylogenetic structure rather than environmental predictors alone (Table S14).

Pairwise correlations among predictors were evaluated for each dataset, and variance inflation factors (VIFs) were calculated to assess collinearity (Tables S15–S16). Correlations were generally moderate and VIF values low, indicating limited collinearity. In particular, MCWD and BIO6 were moderately correlated but retained independent contributions.

Analyses were repeated for ge1, ge5 and ge10 datasets, with emphasis on ge10 as the most conservative threshold. For ge10, lineage-specific effects were tested using models allowing supertribe-specific effects of climatic predictors. Deviations from overall coefficients were used to quantify heterogeneity among lineages (Tables S17–S19).

### Correlated evolution of woodiness and drought/frost

To test whether woodiness and environmental conditions evolved independently or in a correlated manner, we used BayesTraits v5 (*50*) to fit discrete-state continuous-time Markov models on the BrassiToL. Analyses were conducted on the ge10 dataset, using species present in both the phylogenetic tree and niche dataset (*n* = 1,687; Table S20).

For the two environmental variables with significant correlations in our phyloGLM tests (MCWD for drought, BIO6 for frost), species were assigned to one of four combined states representing growth form and environmental condition (Fig. 6A): herbaceous under reduced stress (state 1), herbaceous under increased stress (state 2), woody under reduced stress (state 3), and woody under increased stress (state 4). The model therefore estimates separate transition rates between herbaceous and woody states under contrasting environmental conditions (q13, q24, q31, q42), as well as transitions between environmental states within herbaceous and woody lineages (q12, q21, q34, q43). Because our primary objective was to test whether environmental context altered gains and losses of woodiness, subsequent analyses focused on transition rates involving changes in growth form.

Drought (MCWD) and frost (BIO6) were discretised using percentile-based thresholds (p40, p50, p60). For frost, an additional mechanistic threshold at 0 °C was included to represent freezing conditions (Table S20). For each environmental variable and threshold, we fitted three model classes: an independent model, a dependent four-state model, and a reversible-jump (RJ) dependent model allowing transition-rate parameters to be set to zero to identify model simplifications. Each model was run in 20 independent MCMC chains.

Model performance was evaluated using acceptance rates (Fig. S15), marginal likelihoods estimated by stepping-stone sampling, and effective sample sizes (ESS) (Data file S8). Support for correlated evolution was assessed using log Bayes factors comparing dependent and independent models (Fig. S16 and Table S21). RJ analyses were used to assess model complexity by summarising posterior inclusion frequencies of transition-rate parameters (Table S22).

To evaluate how environmental context influenced the evolution of woodiness, we compared posterior distributions of transition-rate ratios between equivalent transitions under contrasting environmental conditions (Fig. S17 and Table S23). Specifically, we compared rates of gains of woodiness under reduced versus increased stress (q13/q24) and reversals to herbaceousness under reduced versus increased stress (q31/q42).

### Use of generative AI tools

Generative artificial intelligence (AI) tools, including ChatGPT (OpenAI), Gemini (Google) and Ecosia Chat, were used to assist with debugging, optimising and documenting custom bioinformatic scripts used in this study. All generated code was critically evaluated, tested and modified by the authors before use in downstream analyses.

## Supporting information

Data_file_1_Geographic variation in woodiness across Brassicaceae botanical countries

Data_file_2_Provenance and voucher information for samples included in the Brassicaceae Tree of Life

Data_file_3_Catalogue of confident growth-form shifts identified on the Brassicaceae Tree of Life

Data_file_4_Expert-curated occurrence records for 46 rare Brassicaceae species

Data_file_5_Manually validated occurrence records for taxonomically challenging Brassicaceae species

Data_file_6_Species-level climatic niche summary table for Brassicaceae

Data_file_7_Species-level dataset used for non-phylogenetic woodiness analyses

Data_file_8_BayesTraits model diagnostics and MCMC performance metrics

Data_file_9_Summary statistics for data recovery, filtering, and paralog identification

## Acknowledgements

This work was supported by the Dutch Research Council (NWO; grant VI.Veni.222.201 to K.P.H.) and the German Research Foundation (DFG; grant MU1137/17-1 to K.M.). T.M. acknowledges support from the Czech Science Foundation (project no. 24-11371S). S.Š. acknowledges support from the Grant Agency VEGA, Bratislava, Slovakia (grant no. 2/0010/25), awarded to J. Zozomová-Lihová. Sequencing was performed by BaseClear.

This work was partly supported by grants from the Calleva Foundation to the Plant and Fungal Trees of Life (PAFTOL) project at the Royal Botanic Gardens, Kew. We thank the many herbaria, institutions, curators, collectors, taxonomists and collaborators who contributed specimens, expertise and associated metadata to the Brassicaceae Tree of Life (BrassiToL). An overview of contributing collections, specimens and associated metadata is available through the BrassiToL portal. We are particularly grateful to Eric Roalson for sharing growth-form data for Cleomaceae, Carlos Salazar for facilitating access to material of *Moricandia rytidocarpoides*, and The Emirates Center for Wildlife Propagation (Morocco) for its support. We thank Laura van Hoek, Rosario Franco Berriel and Ryan Brewer (Naturalis Biodiversity Center) for sharing raw data and for laboratory and technical support.

We also thank the herbaria and botanical institutions that provided material for genomic analyses or housed voucher specimens associated with analysed samples. Herbarium acronyms follow Index Herbariorum. A complete list of contributing institutions, specimen providers, collectors and associated acknowledgements is provided through the BrassiToL portal and in Data file S2. We are grateful to all contributors who helped build the Brassicaceae Tree of Life over the course of the project.

We acknowledge all national, regional and local authorities that authorised, facilitated or supported the collection, transfer and study of plant material used in this research.

## Author contributions

K.P.H., K.M. and F.L. conceived and coordinated the study. K.P.H. designed the analytical framework, curated the data, developed computational workflows, performed all analyses, generated the figures and wrote the first draft of the manuscript. F.L. and K.M. contributed to study design, supervision, interpretation of the results and manuscript development. D.A.G. and I.A.A.-S. contributed substantially to study design, taxonomic expertise, specimen acquisition, data curation, interpretation of the results and manuscript revision. W.J.B., M.E.S., C.Kie. and M.A.K. contributed to project development, study design, interpretation of the results and manuscript revision.

Laboratory work, data generation and associated investigations were performed by K.P.H., L.v.S., C.Ku., L.J.M.A.D., M.K., L.E.T., E.C.-L. and S.v.Z. D.S.A. contributed to software development and digital research infrastructure. C.Kie. contributed to methodology development, validation, formal analyses, project administration and data curation. N.W. contributed to data curation, interpretation of the results and manuscript development.

Specimens, occurrence records, taxonomic expertise and other essential resources were contributed by I.A.A.-S., D.A.G., M.A.K., A.R.Z., A.F., A.G., A.P.S., A.B., A.E.-F., B.Ö., C.D.B., D.J.S., E.B., F.F., F.W.S., H.M., H.S., H.C., L.M.D., M.A.L., M.B., M.C., M.D.W., M.T., M.S., N.M.H., N.M.S., N.W., O.M., O.T.-N., P.H., P.J.W., R.N., R.V., S.Š., S.B.J., T.M. and V.R.I. L.A.N. contributed to study development, interpretation of the results and manuscript revision.

Funding acquisition was led by K.P.H., F.L. and K.M., with additional support from W.J.B., F.F. and S.Š. All authors contributed to the interpretation of the results, critically revised the manuscript and approved the final version.

## Data availability

The raw sequencing reads generated in this study will be deposited in the NCBI Sequence Read Archive (SRA) under BioProject PRJNA1482360 (accession numbers listed in Data file S2). Previously published sequencing data were obtained from public repositories as detailed in Data file S2. Species occurrence records, growth-form classifications and environmental niche data underlying the analyses are provided in Data files S1–S9.

## Code availability

All custom scripts and software used for sequence processing, phylogenetic analyses, occurrence filtering, georeferencing, record validation and statistical analyses are available from Zenodo (doi.org/10.5281/zenodo.20931862).

