## Supplementary materials for "Why woodiness repeatedly evolves—and disappears"

Kasper P. Hendriks *et al*.

---

#### Contents

Supplementary Results
Supplementary Figures 1–18
Supplementary Tables 1–23
Supplementary References

#### *Supplementary Results*

### Overview of the Brassicaceae Tree of Life

##### Overview

We reconstructed a new Brassicaceae Tree of Life (BrassiToL) comprising 2,927 Brassicaceae species, 135 Cleomaceae species and 23 representatives of the remaining Brassicales families (Fig. 2, Supplementary Fig. 1 and Supplementary Data 2). Because the full phylogeny is too large to visualise effectively in print, an interactive version is available online (https://tol.naturalis.nl/collection/brassicaceae). The family crown age was estimated at 36.9 Ma (Q1 = 0.934, LPP = 1, sCF = 61.9, gCF = 64), with a stem age of 54.8 Ma (Supplementary Table 1).

Consistent with previous phylogenomic studies (*1–3*), most tribes were recovered as well-supported monophyletic groups (Extended Data Fig. 3 and Supplementary Table 1). Tribe crown nodes generally received higher support than contemporary nodes elsewhere in the phylogeny, supporting the utility of tribal classification as a taxonomic framework within Brassicaceae. The mean tribe crown age was 12.3 Ma (range 1.7–27.8 Ma).

Of the 58 currently recognised tribes, tribes Biscutelleae, Iberideae and Subularieae were recovered as non-monophyletic, with constituent genera occurring in separate parts of the phylogeny (Supplementary Fig. 1), largely corroborating earlier findings[2].

##### Backbone relationships

The Brassicaceae comprise two subfamilies: the early-diverging Aethionemoideae, represented by tribe Aethionemeae, and the much larger Brassicoideae containing the remaining 57 tribes (*4*). Within Brassicoideae, five supertribes are currently recognised: Arabodae, Brassicodae, Camelinodae, Heliophilodae and Hesperodae.

Consistent with previous nuclear phylogenomic studies (*1–3*), Aethionemoideae was recovered as sister to all remaining Brassicaceae, and Hesperodae as the earliest-diverging supertribe within core Brassicaceae (Fig. S18). We further recovered a clade comprising Brassicodae, Camelinodae and Heliophilodae, although the branching order among these supertribes differed from previous analyses, with Brassicodae recovered as sister to Camelinodae plus Heliophilodae.

As noted previously (*3*), backbone relationships among the supertribes remain characterised by short internal branches and conflicting phylogenetic signal. Support values for these deep divergences were consistently lower than for most tribal relationships (Extended Data Fig. 3), indicating that the earliest diversification of core Brassicaceae remains incompletely resolved.

**Rogue tribes**

Several Brassicaceae tribes have previously been identified as phylogenetically unstable (rogue tribes), with placements that vary among datasets and analytical approaches, in some cases reflecting ancient hybridisation events (*5*–*8*).

Our BrassiToL largely corroborates previous nuclear phylogenomic analyses (*3*). Cochlearieae was recovered as sister to supertribe Brassicodae. Anastaticeae, Megacarpaeeae and *Iberis* (Iberideae) formed a clade within Heliophilodae, together with part of Biscutelleae (*Lunaria* and *Ricotia*). Biscutelleae, Iberideae and Subularieae were each recovered as non-monophyletic, with their constituent genera occupying separate positions in the phylogeny. In particular, *Teesdalia* grouped with *Subularia* and allied tribes, whereas *Idahoa* grouped with Asteae, supporting the interpretation of Subularieae as another rogue tribe. Overall, the placements recovered here are highly consistent with previous nuclear phylogenomic analyses (*3*).

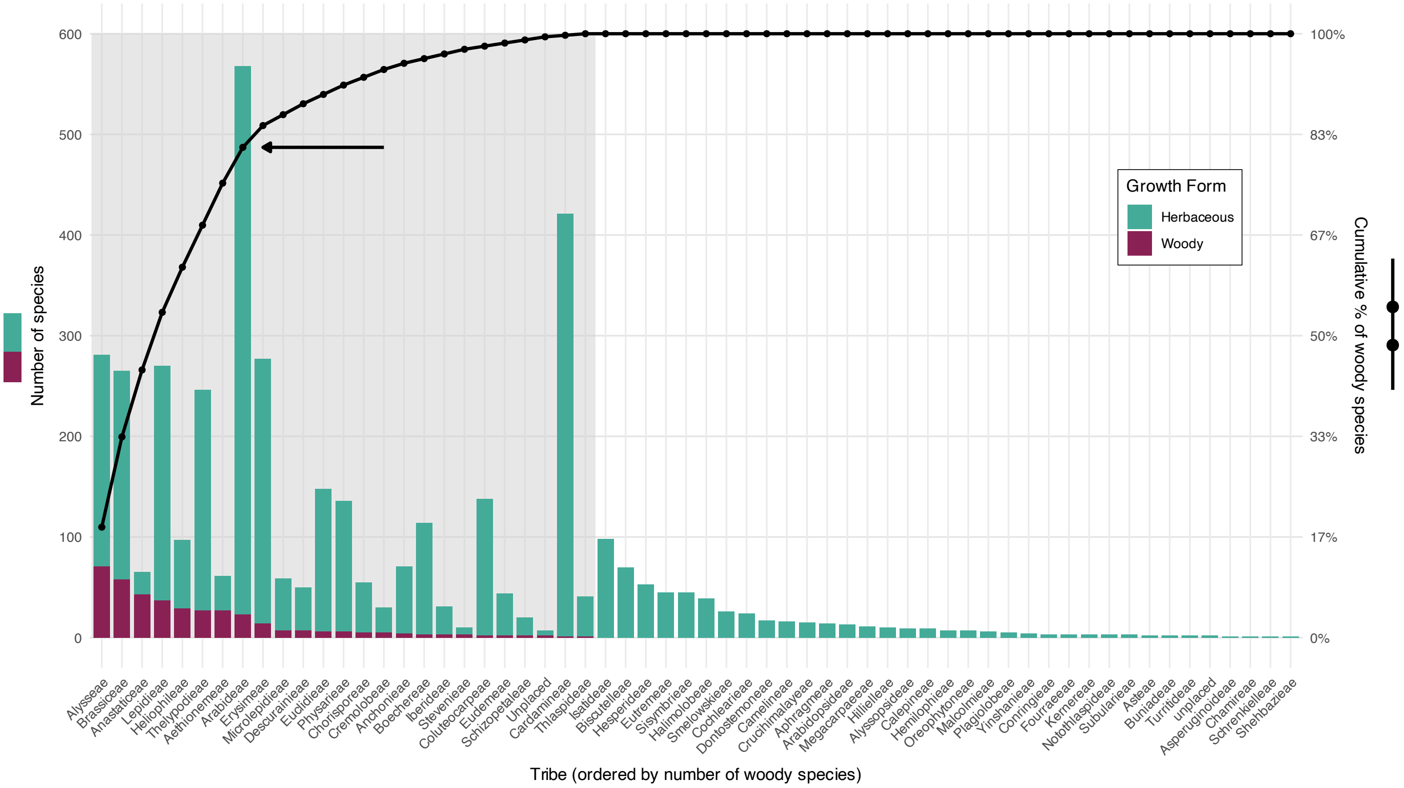


**Fig. S1. Derived woodiness is concentrated in a small subset of Brassicaceae tribes**
While 26 of the 58 Brassicaceae tribes contain at least one woody species, more than 80% of all woody species occur in just eight tribes. Tribes containing woody species are highlighted against a grey background.

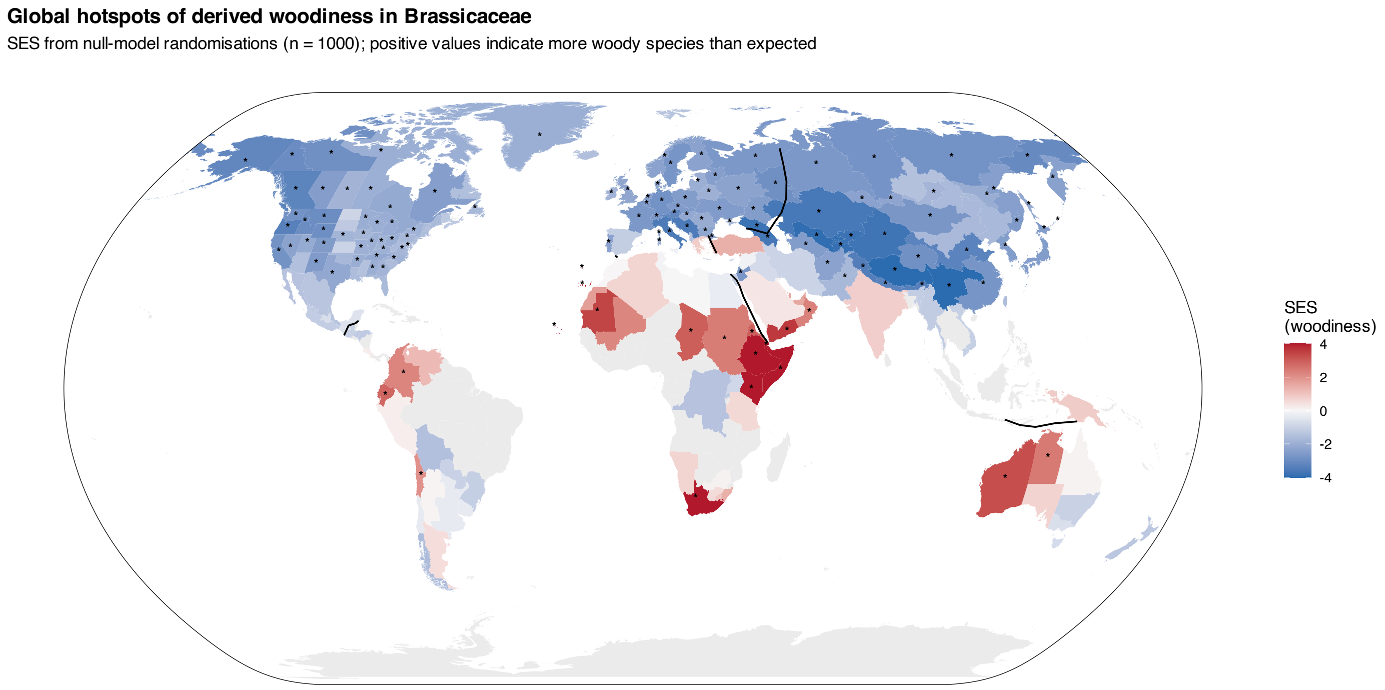


**Fig. S2. Geographic enrichment and depletion of woodiness across Brassicaceae** Botanical countries (WGSRPD Level 3) are coloured according to the standardised effect size (SES) of woody-species richness relative to null expectations based on total Brassicaceae richness. Positive values indicate regions containing more woody species than expected given their Brassicaceae diversity, whereas negative values indicate regions containing fewer woody species than expected. SES values were calculated from 1,000 randomisations, preserving total species richness per botanical country and the global number of woody species. Asterisks indicate significant deviations from null expectations (empirical $P \leq 0.05$). Countries containing fewer than 10 Brassicaceae species were excluded.

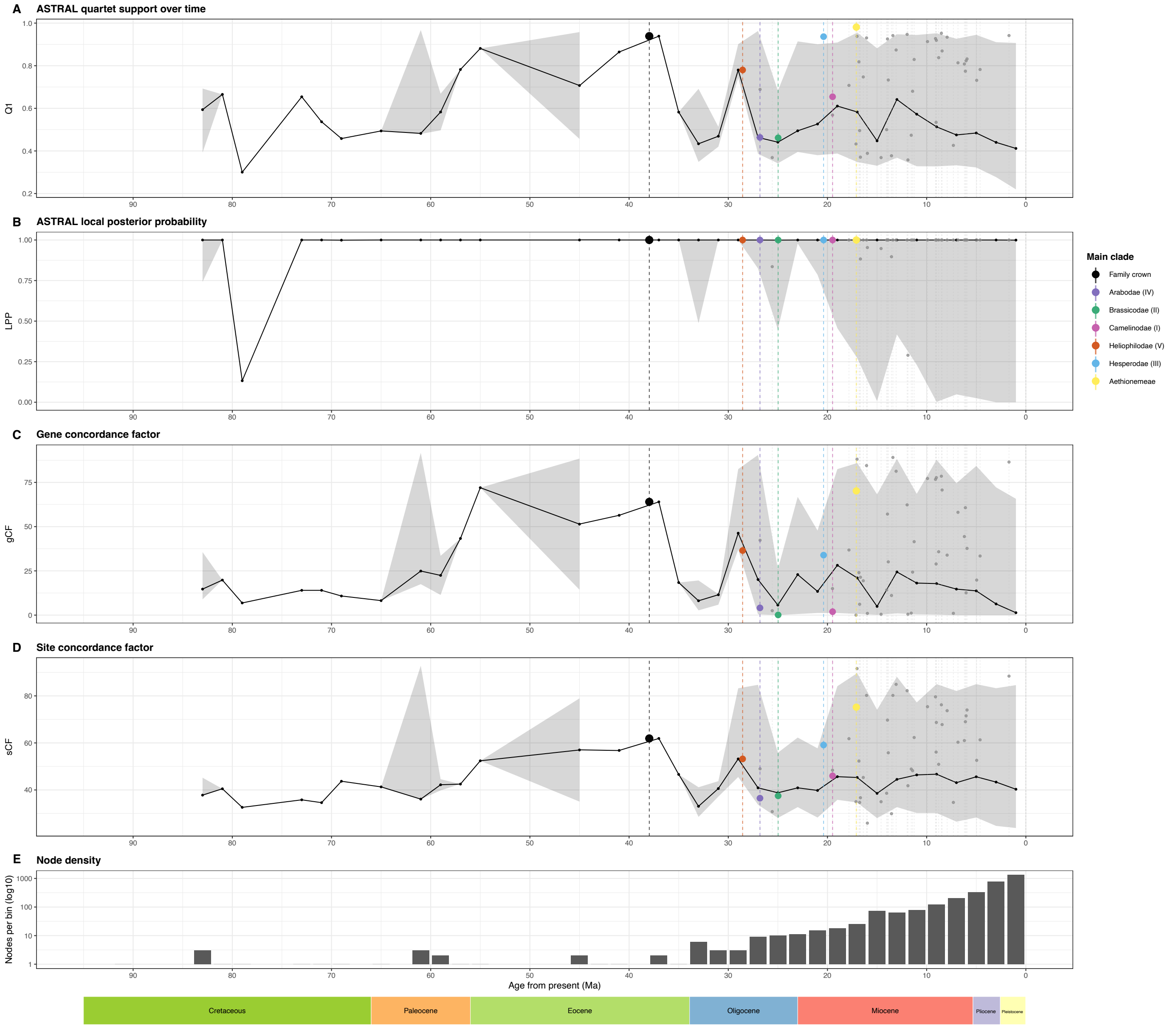


**Fig. S3. Support and concordance metrics across the Brassicaceae Tree of Life**
(A) Median ASTRAL quartet support (Q1), (B) local posterior probability (LPP), (C) gene concordance factor (gCF), and (D) site concordance factor (sCF) across non-overlapping 2-Myr age bins, with shaded areas indicating 95% quantile intervals. Vertical dashed lines and coloured points indicate crown nodes of the Brassicaceae family, the five supertribes, and tribe Aethionemeae, sister to the core Brassicaceae; grey lines and points indicate tribe crown nodes. (E) Number of internal nodes per 2-Myr age bin (log10 scale). Geological epochs are indicated below the panels.

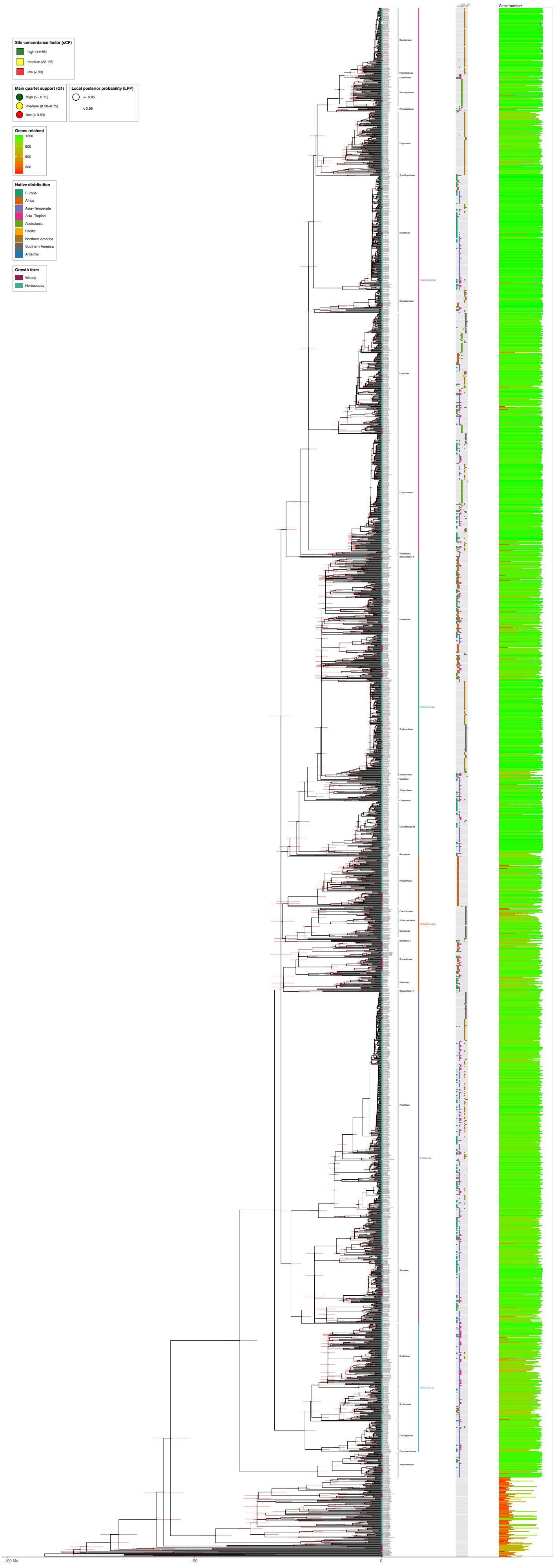


**Fig. S4. Time-calibrated Brassicaceae phylogeny**
Time-calibrated species phylogeny of Brassicaceae inferred from nuclear phylogenomic data. Branch lengths are proportional to time (Ma) and were calibrated using LSD2 on a constrained species-tree topology inferred from 1,052 nuclear loci. Red horizontal bars indicate 95% confidence intervals for node ages. Internal nodes are annotated using multilayer concordance and branch-certainty metrics. square symbols indicate site concordance factor (sCF), circle fill colour indicates the proportion of quartets supporting the main topology (Q1), and circle outline indicates local posterior probability (LPP). Tribes represented by multiple species are annotated along the right margin. Tip labels include accepted species names and sample identifiers; a red font indicates the sample derives from a type specimen, an * indicates samples derive from the backbone phylogeny of Hendriks et al. (2023)[2]. Additional side panels show the number of retained loci per sample and native continental distribution based on WGSRPD Level-1 regions.

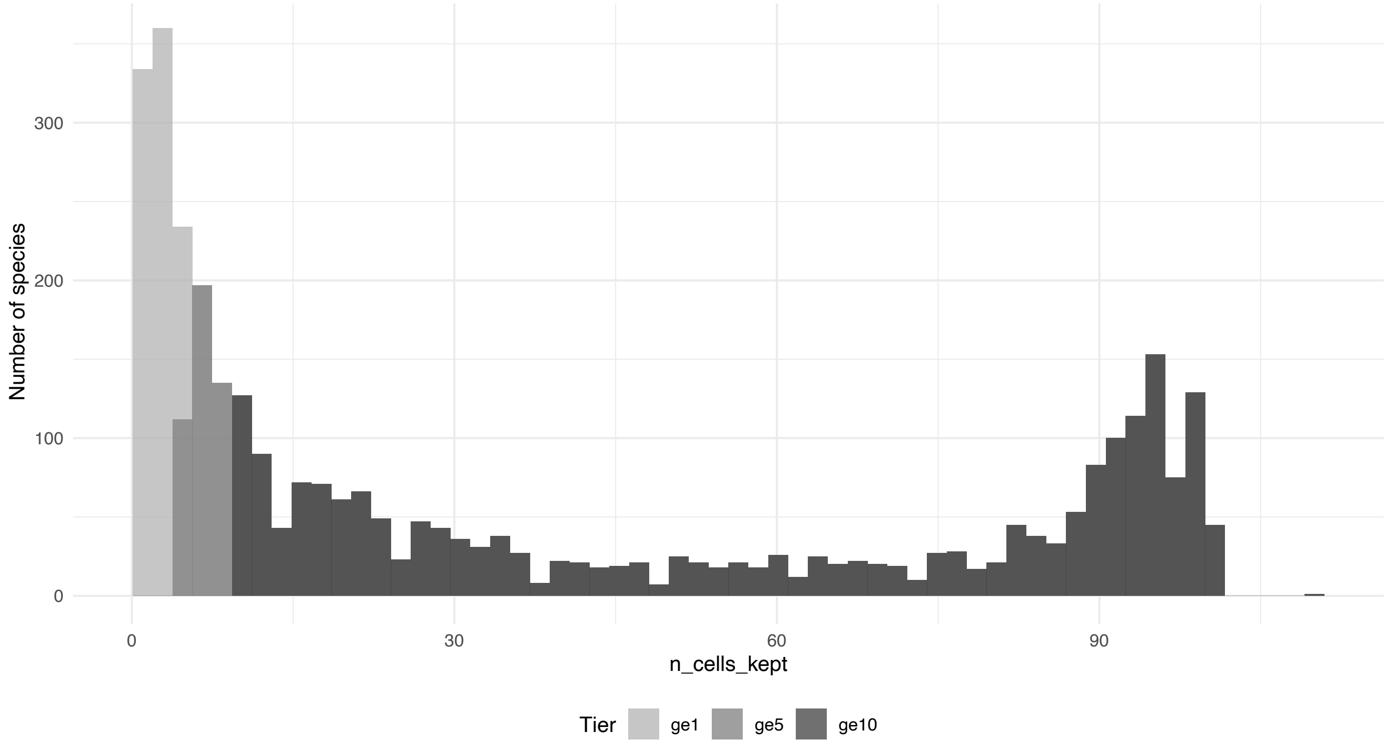


**Fig. S5. Species sampling depth for niche calculations**
Species niches were defined using climatic values extracted from global 30 arc-second (~1 km at the equator) raster cells (WorldClim), where each value represents the conditions of the corresponding grid cell (defined at its centre). Occurrence records were thinned to at most one per raster cell, and only records passing our automated and expert-curated pipeline were retained. Species were analysed under three sampling tiers based on the number of retained raster cells. ge1 (≥1 cell), ge5 (≥5 cells), and ge10 (≥10 cells).

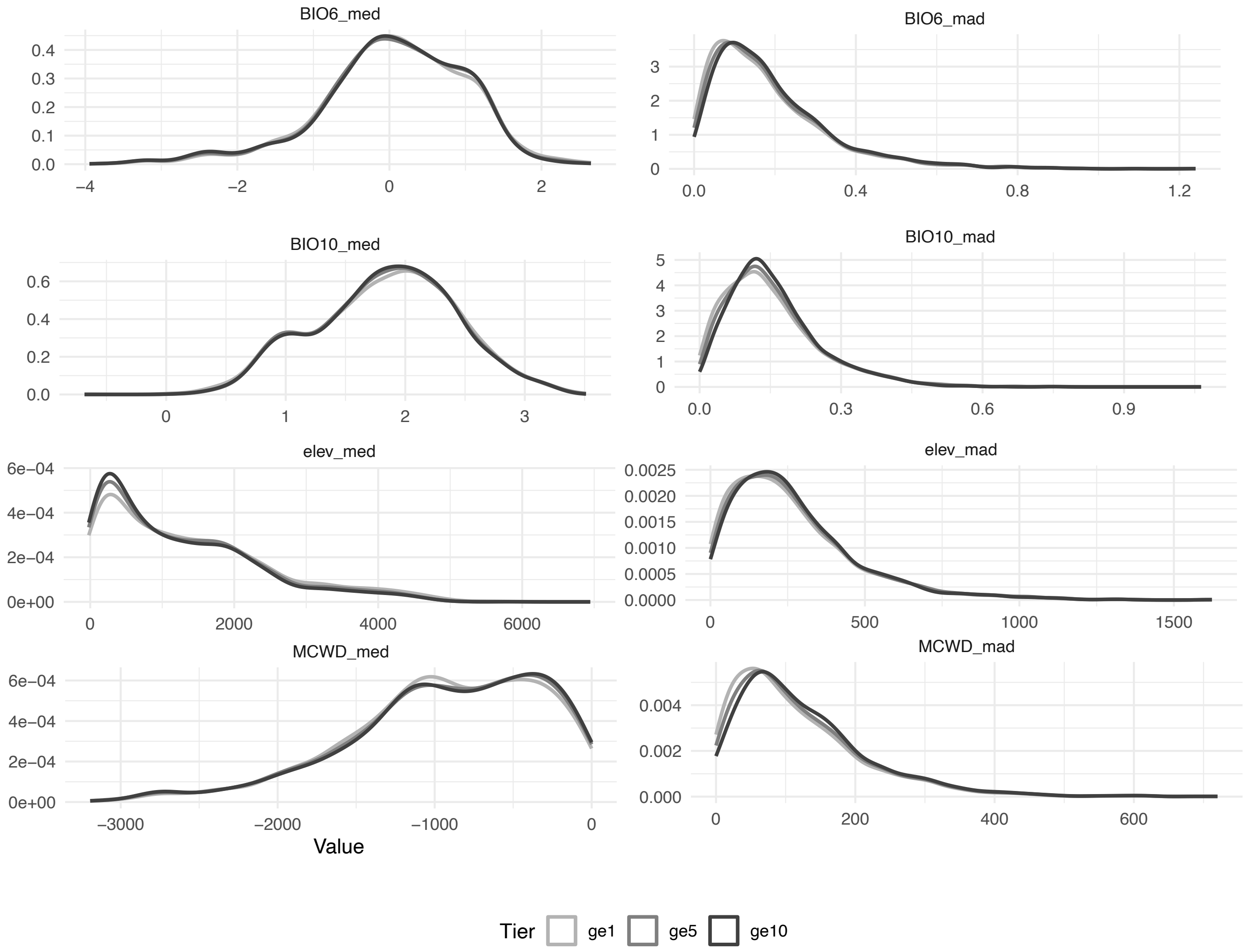


**Fig. S6. Species-level niche distributions are consistent across sampling thresholds**
Density distributions of species-level climatic niche summaries across the three retained-cell thresholds used in downstream analyses. ge1, ge5 and ge10. Each panel shows one niche variable, summarised per species from retained raster cells after occurrence filtering and thinning. The strong overlap among tiers indicates that increasing the minimum number of retained cells per species has relatively little effect on the overall distribution of inferred niche positions and breadths, supporting the robustness of downstream analyses to sampling threshold.

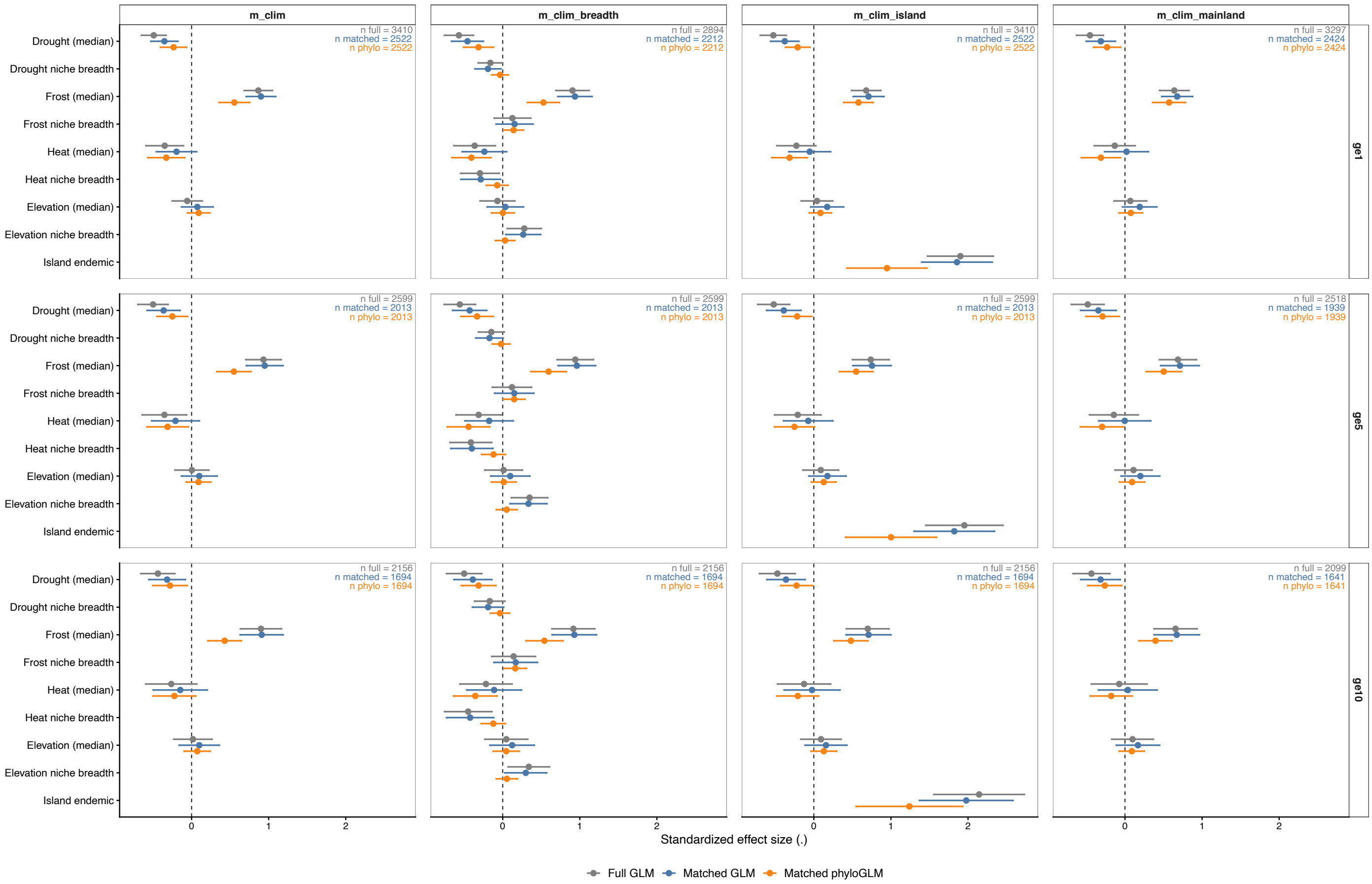


**Fig. S7. Integrated climate–woodiness model coefficients across thresholds and modelling frameworks**

Estimated standardised effect sizes (β ± 95% Wald CI) for climatic niche predictors in integrated models of woodiness across the three niche-data thresholds (ge1, ge5, ge10) and four model formulations. Results are shown for the full non-phylogenetic dataset (glm_full), the matched non-phylogenetic subset (glm_matched), and the matched phylogenetic subset (phyloglm_matched). Predictors include climatic niche medians and, where applicable, niche-breadth terms for drought (MCWD), frost (BIO6), heat (BIO10), and elevation, plus island endemism in the island model. This figure is intended to compare the overall consistency of effect directions and magnitudes across data subsets and phylogenetic treatments.

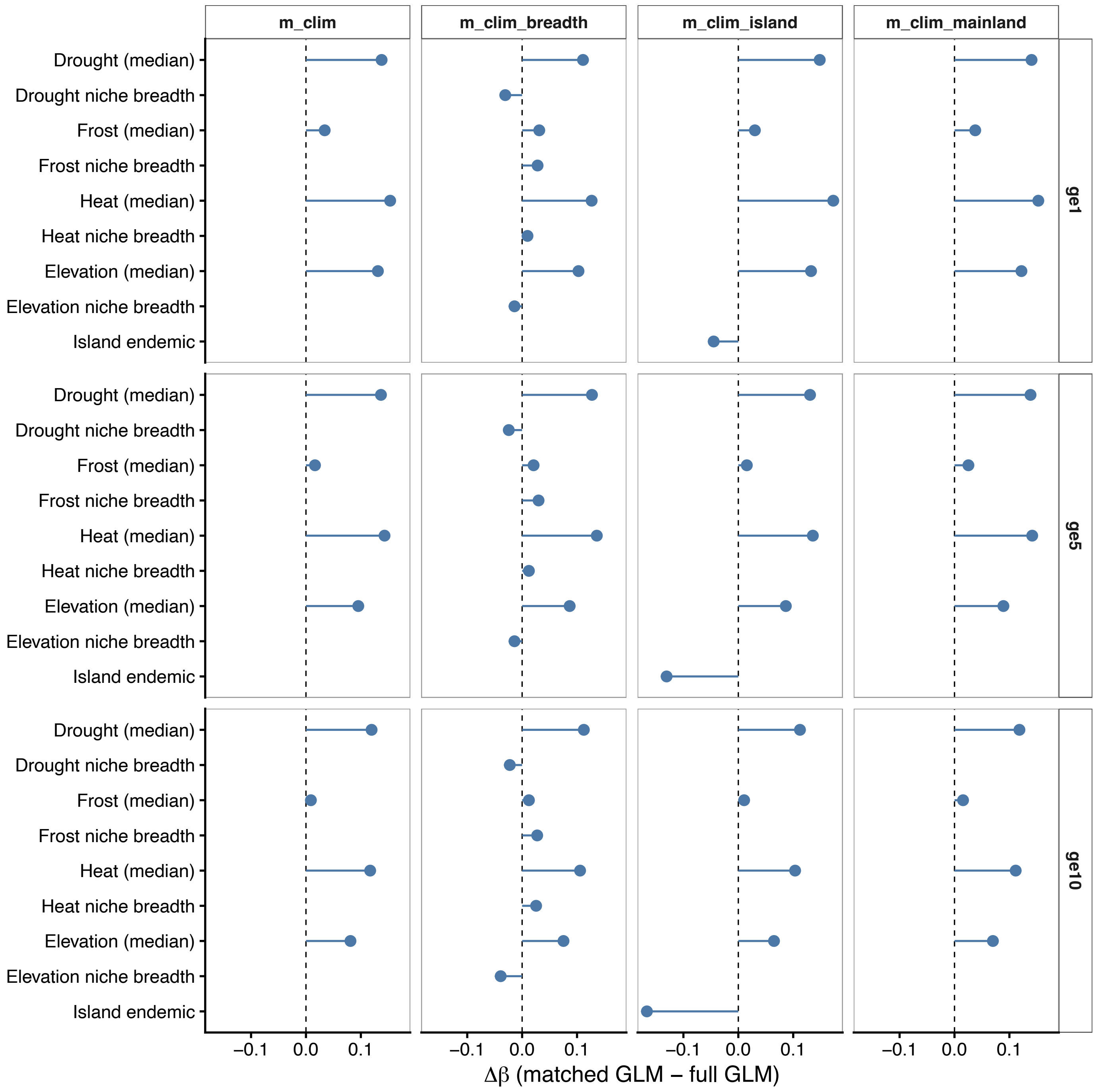


**Fig. S8. Effect of dataset restriction on estimated climate–woodiness relationships**
Differences in standardised regression coefficients (Δβ) between models fitted to the matched niche–tree subset and the full niche dataset (Δβ = β_{glm_matched} – β_{glm_full}). Positive values indicate stronger effects after restricting analyses to species included in the phylogenetic dataset, whereas negative values indicate weaker effects. Results are shown for all climatic predictors and niche breadth variables across ge1, ge5, and ge10 tiers and across all model formulations.

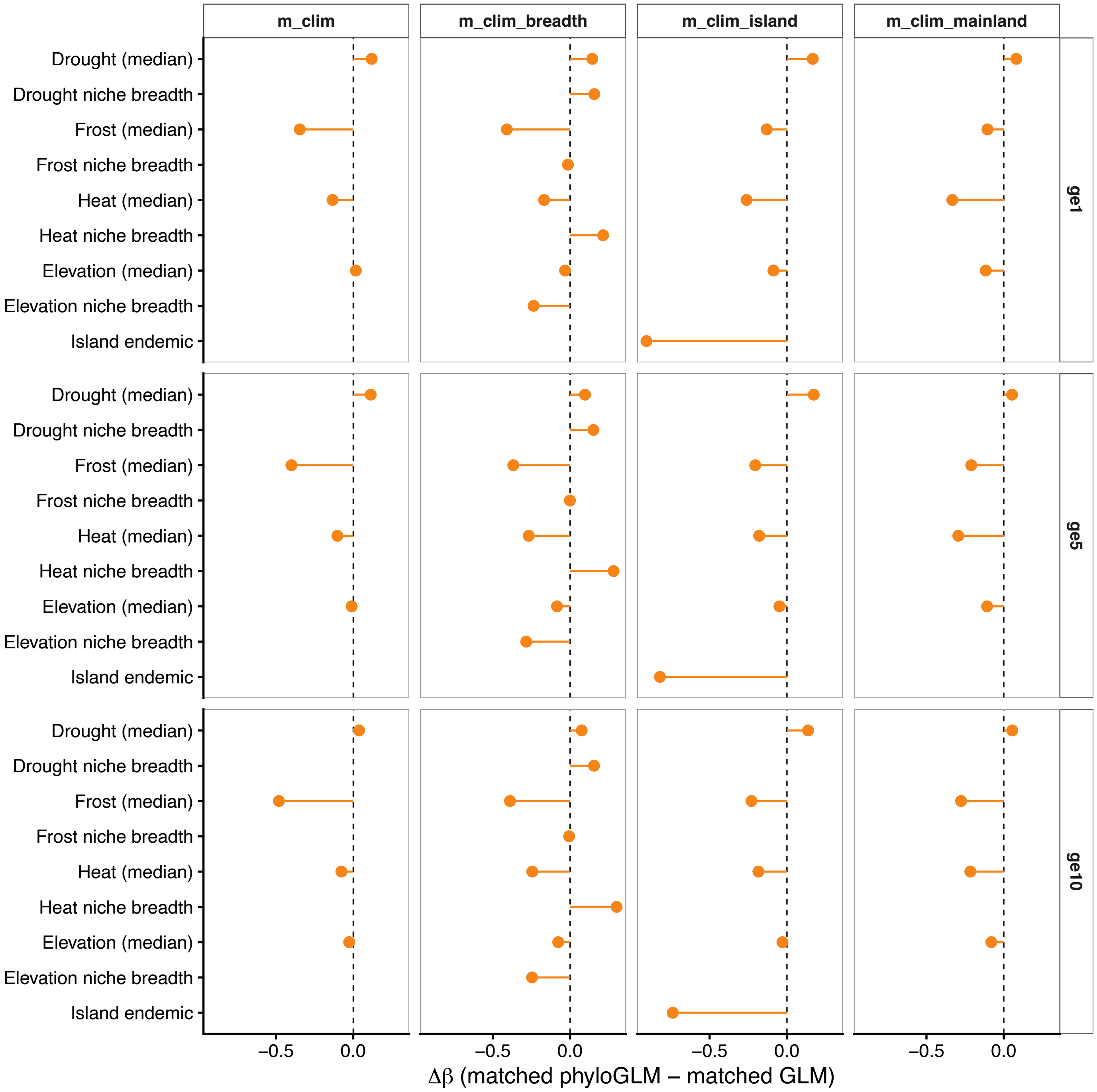


**Fig. S9. Effect of phylogenetic correction on estimated climate–woodiness relationships**
Differences in standardised regression coefficients (Δβ) between phylogenetic and non-phylogenetic models fitted to identical matched datasets (Δβ = β_{phyloglm_matched} – β_{glm_matched}). Positive values indicate stronger effects after accounting for phylogenetic relatedness, whereas negative values indicate weaker effects. Results are shown for all climatic predictors and niche breadth variables across ge1, ge5, and ge10 tiers and across all model formulations.

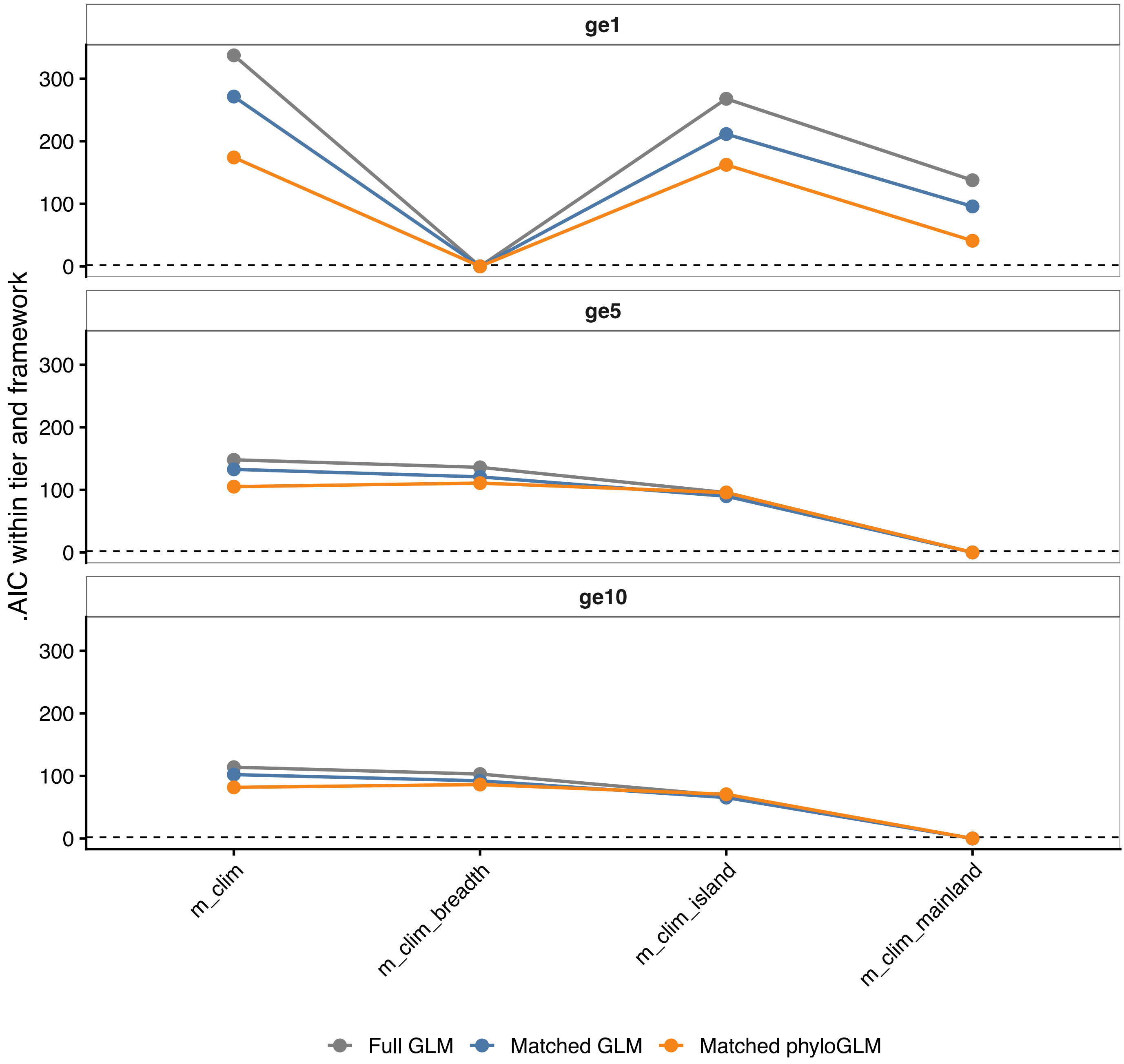


**Fig. S10. Model support across statistical frameworks and dataset tiers**
Comparison of model fit based on ΔAIC within each framework (GLM full, GLM matched, phylogenetic GLM) and ge tier (ge1, ge5, ge10). For each framework–tier combination, ΔAIC is calculated relative to the best-fitting model (ΔAIC = 0). Lower values indicate stronger support. Lines connect models within frameworks to highlight relative ranking consistency across model formulations.

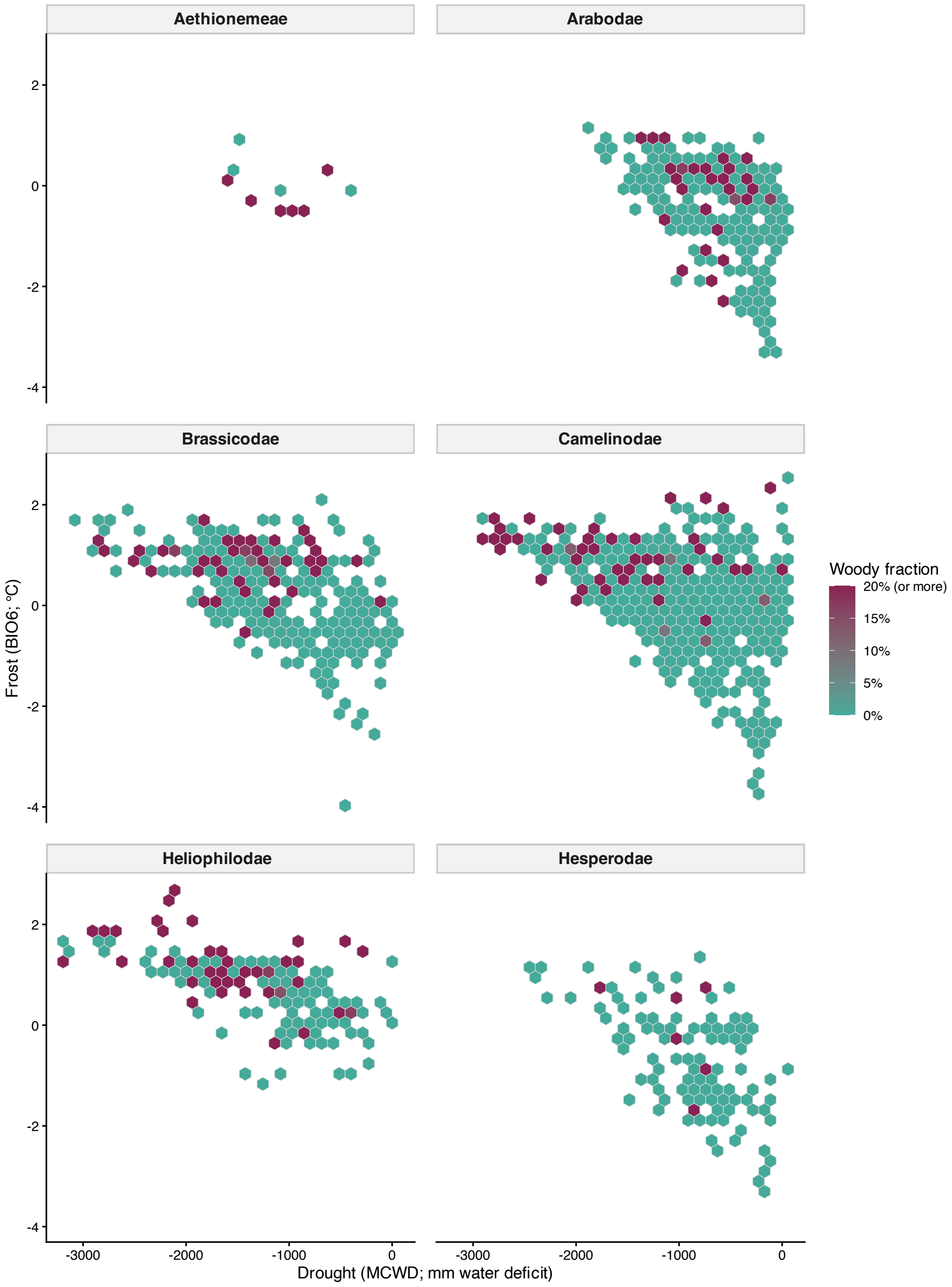


**Fig. S11. Distribution of woodiness across climatic niche space in the six main Brassicaceae lineages**

Hexagon-binned climatic niche space for the six main Brassicaceae lineages: tribe Aethionemeae and the five core Brassicaceae supertribes. Axes represent species-level median values of Maximum Cumulative Water Deficit (MCWD; mm water deficit) and BIO6 (minimum temperature of the coldest month; °C), calculated from retained occurrence cells in the ge10 dataset. Hexagon fill indicates the fraction of woody species within each niche-space bin. The colour scale is capped at 20% woody species to improve visual resolution across the observed range. More negative MCWD values indicate drier conditions, whereas higher BIO6 values indicate milder, less frost-prone winters.

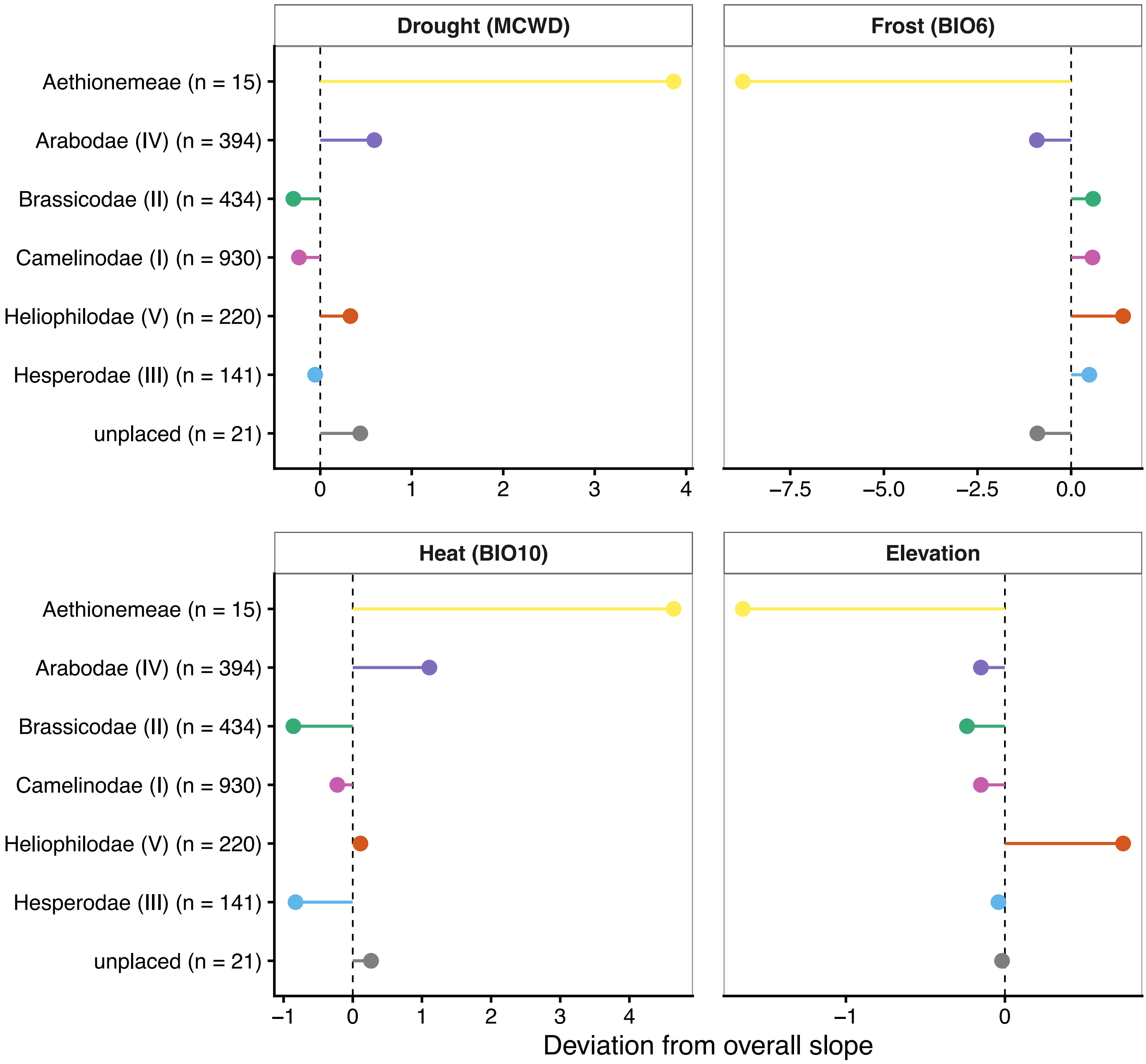


**Fig. S12. Main lineage deviations from overall climate–woodiness relationships in Brassicaceae**

For each climatic predictor, points show the deviation (Δβ) between the main lineage-specific slope and the corresponding overall slope estimated across all included main lineages in the ge10 analysis. Positive Δβ values indicate that the association between climate and woodiness is stronger within that main lineage than across Brassicaceae overall, whereas negative Δβ values indicate a weaker association. For example, positive drought Δβ values indicate that woodiness increases more strongly toward drier conditions within that lineage, while negative frost Δβ values indicate a weaker positive association between woodiness and mild winters relative to the family-wide trend. Slopes were estimated from a non-phylogenetic interaction model allowing supertribe-specific responses to drought (MCWD), frost (BIO6), heat (BIO10), and elevation, using standardised predictor values. Only main lineages exceeding the minimum sample-size threshold were included; sample sizes are indicated on the y-axis.

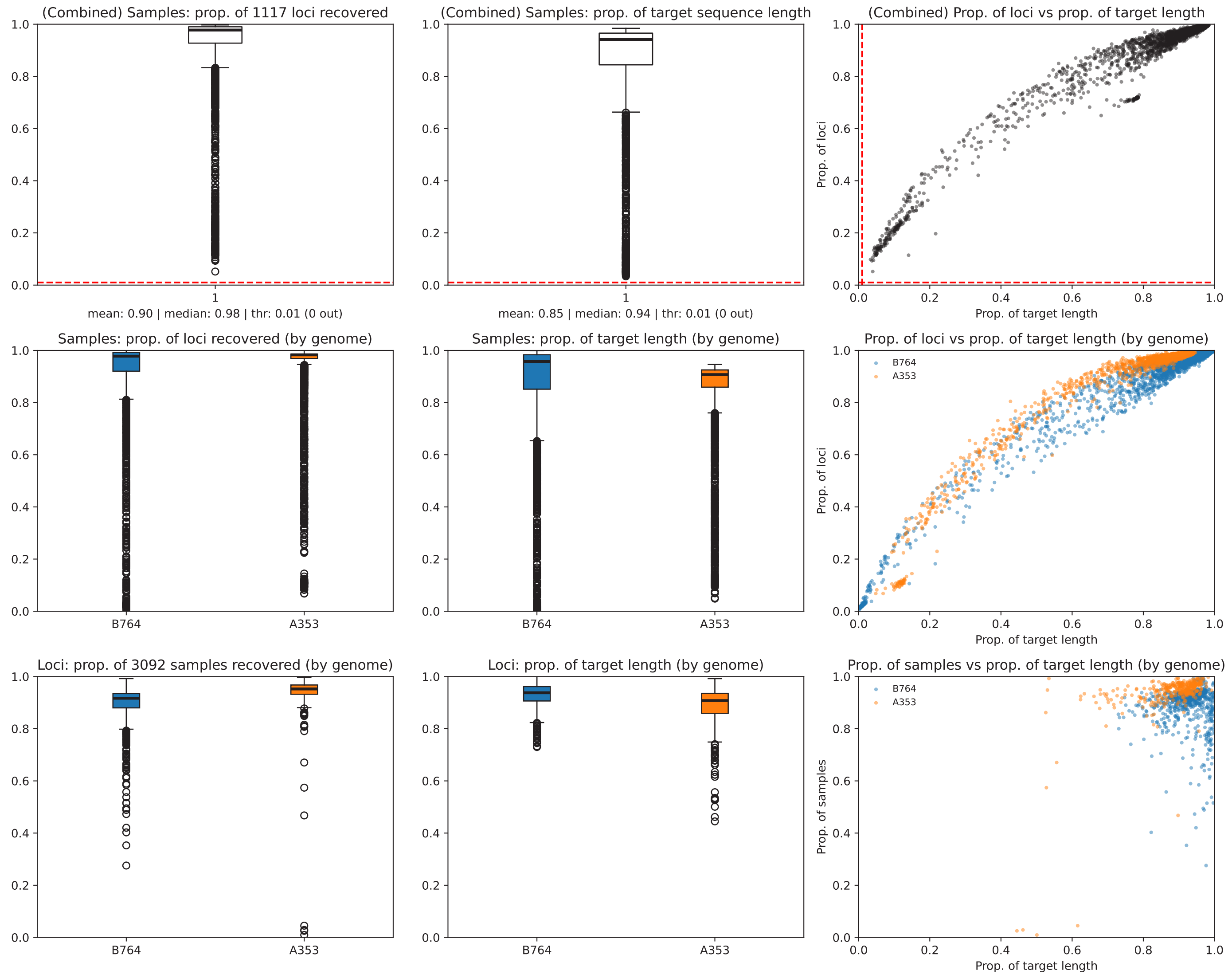


**Fig. S13. Overview of sequencing data recovery and locus assembly success**
Overview of sequencing and assembly success across all samples included in this study. The figure summarises key metrics such as total reads retained after trimming, number of loci recovered per sample, and sequence length distributions following HybPiper assembly of the Brassicaceae-specific baitset (B764) and the angiosperm-wide baitset (A353). Data are shown prior to HybPhaser filtering. This overview demonstrates overall dataset completeness and variation in recovery efficiency among samples. The red dotted lines indicate the minimum coverage threshold for the proportion of samples and the proportion of target length set in HybPhaser.

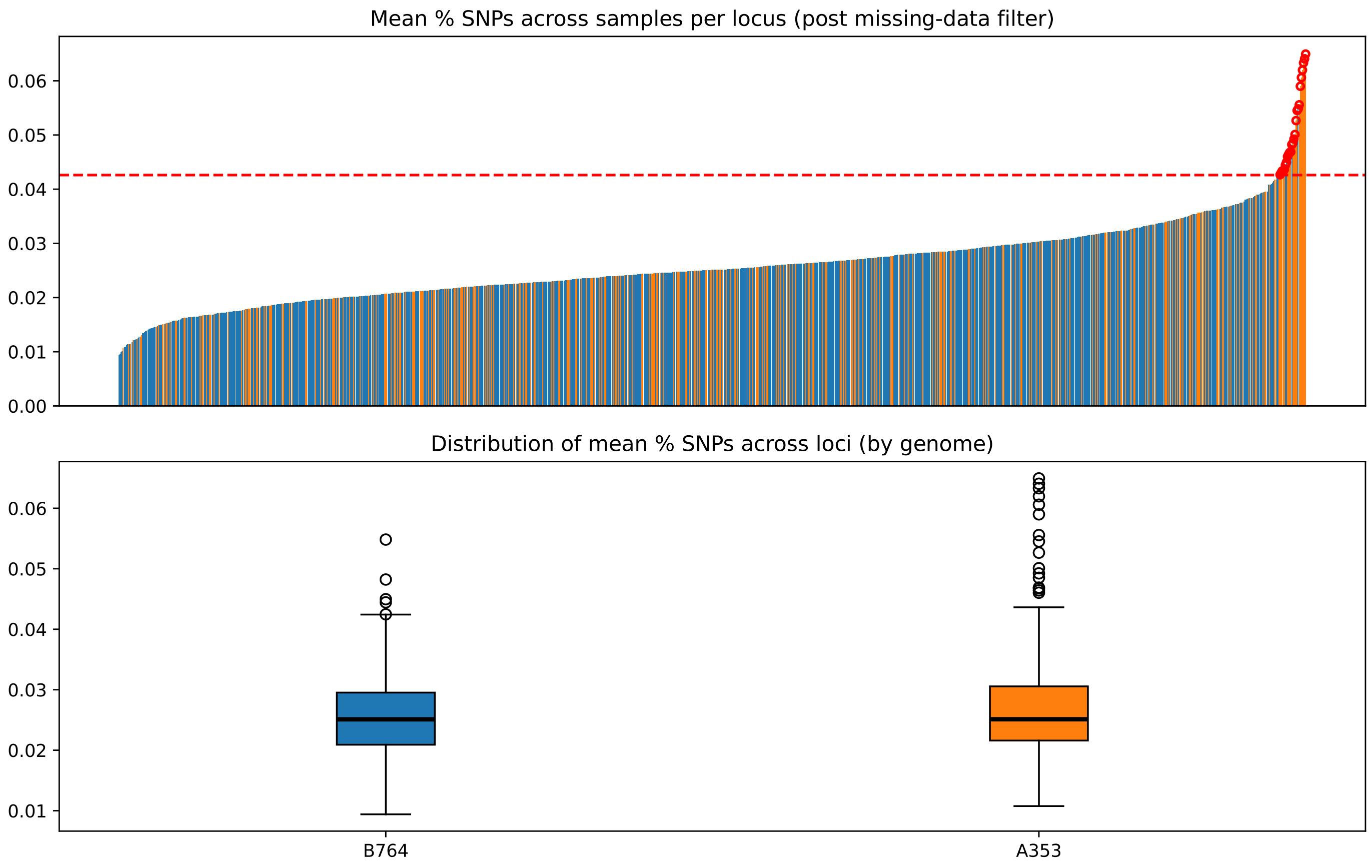


**Fig. S14. SNP distribution across genes**
Distribution of putative paralog occurrence across all samples for all genes, based on HybPhaser SNP assessment. Each bar represents a gene, and red circles highlight outlier loci flagged as paralogous, which were removed from subsequent analyses. The red dotted line indicates the global outlier threshold for SNP percentage set in HybPhaser to detect and remove putative paralogous genes.

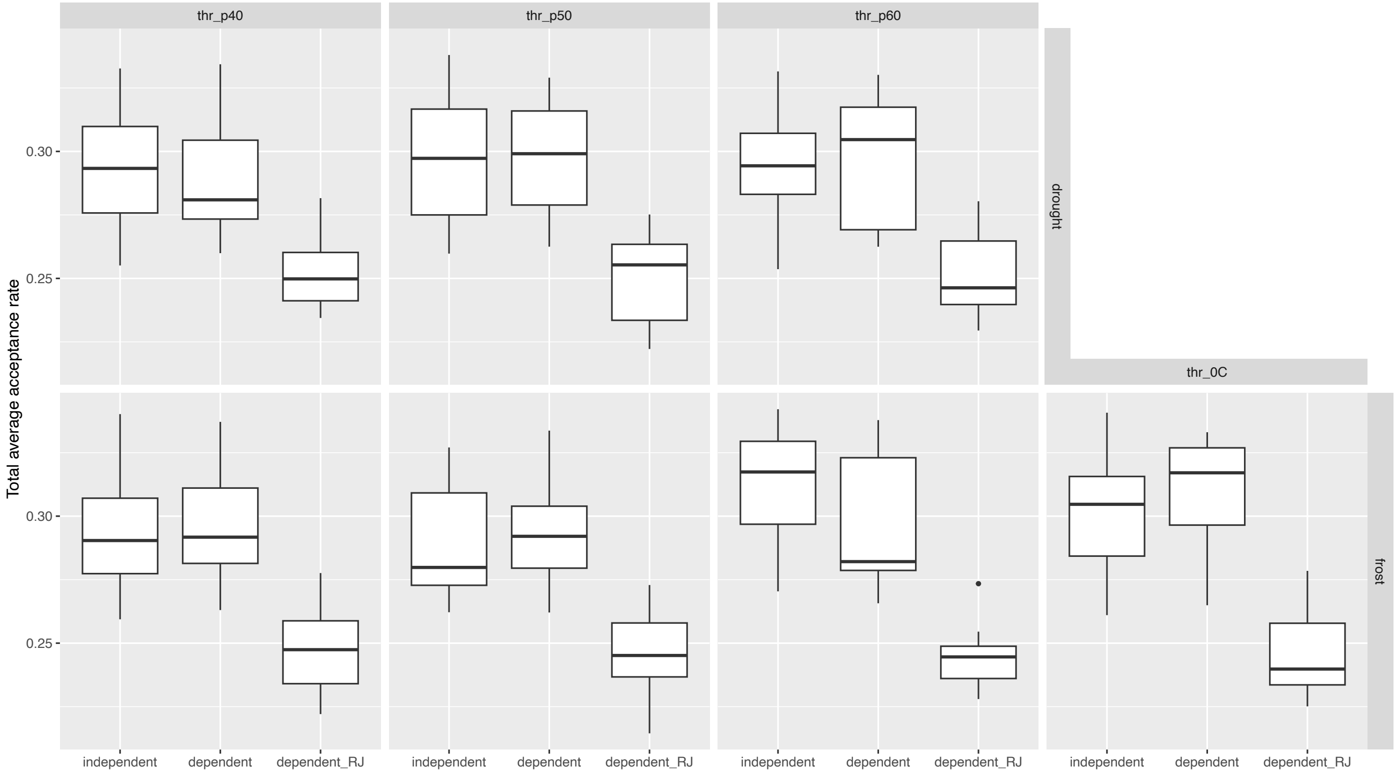


**Fig. S15. MCMC acceptance rates across BayesTraits models, analyses, and thresholds** Distribution of total average proposal acceptance rates across BayesTraits chains for all models (independent, dependent and reversible-jump dependent), analyses (drought, frost) and threshold definitions. Boxplots summarise acceptance rates across independent chains. Acceptance rates provide a diagnostic of MCMC performance, with values that are neither excessively low nor excessively high generally indicating efficient exploration of parameter space. The consistency of acceptance rates across analyses supports adequate mixing and convergence behaviour.

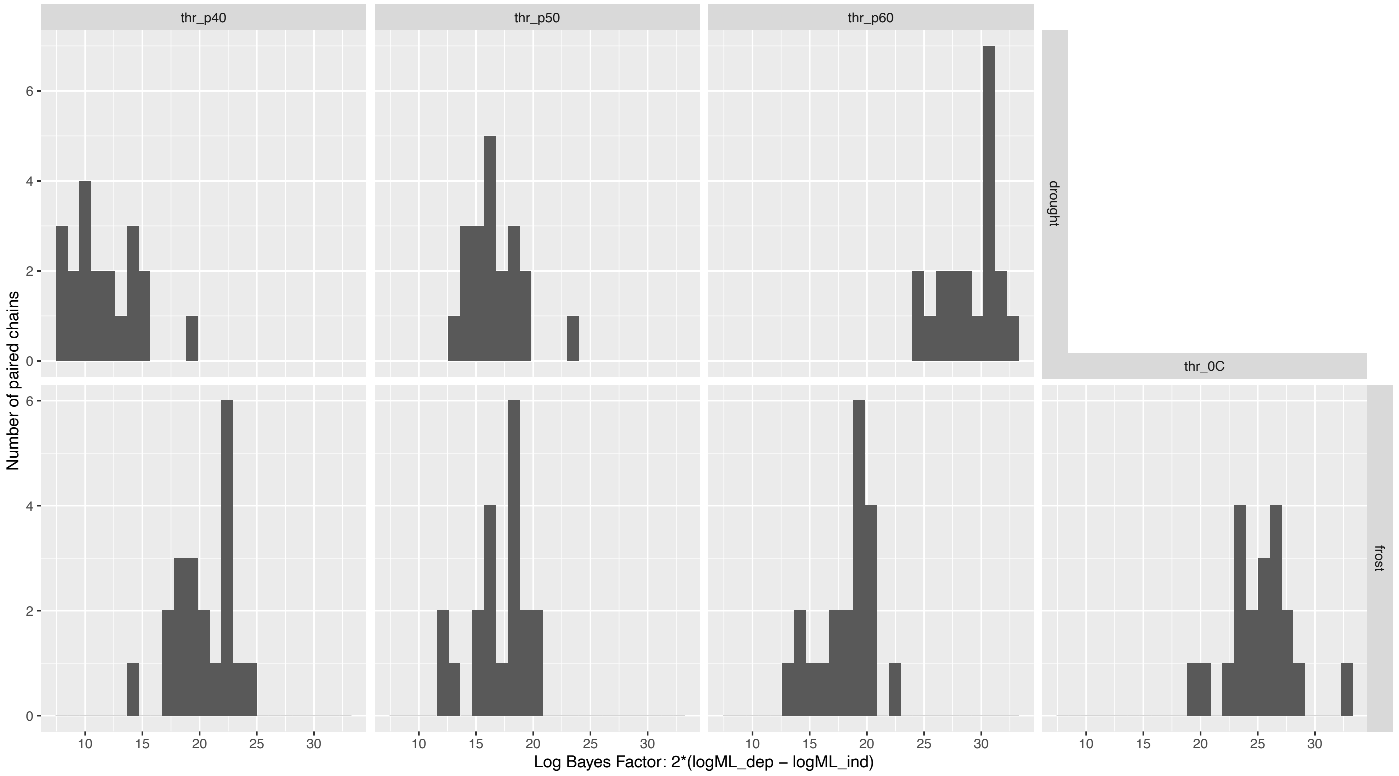


**Fig. S16. Bayes factor support for correlated evolution between woodiness and environment across analyses and thresholds**

Distributions of log Bayes factors (logBF = 2 × [log marginal likelihood dependent – log marginal likelihood independent]) comparing dependent and independent BayesTraits models across analyses (drought, frost) and threshold definitions. Each value represents a paired comparison from a single chain based on stepping-stone marginal likelihood estimates. Larger log Bayes factor values indicate stronger support for the dependent model, consistent with correlated evolution between woodiness and environmental regime. Variation among chains reflects Monte Carlo uncertainty in marginal likelihood estimation.

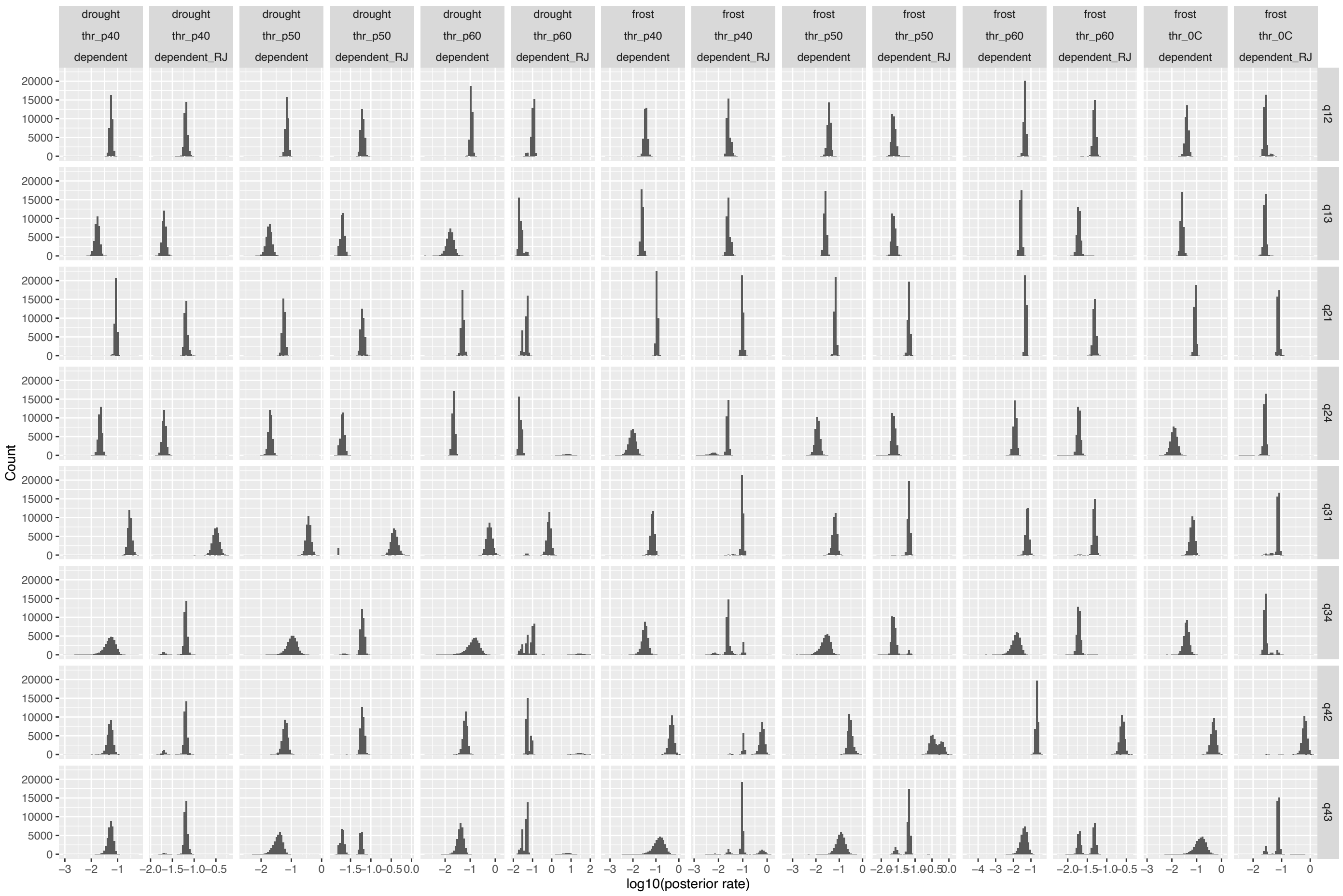


**Fig. S17. Posterior distributions of all transition rate parameters (q) across BayesTraits analyses, thresholds, and models**

Posterior distributions of all transition rate parameters (q) estimated under the dependent and reversible-jump (RJ) BayesTraits models, shown on a log10 scale for comparability. Panels are stratified by parameter (q), analysis (drought, frost), threshold definition, and model. Only finite, positive rate estimates are shown. These distributions provide a comprehensive view of parameter behaviour across the full model space and demonstrate that posterior estimates are well-behaved and not driven by extreme or poorly identified values. RJ results are included as a sensitivity analysis; under RJ, parameters may be set to zero in a subset of samples, reflecting model parsimony rather than effect size.

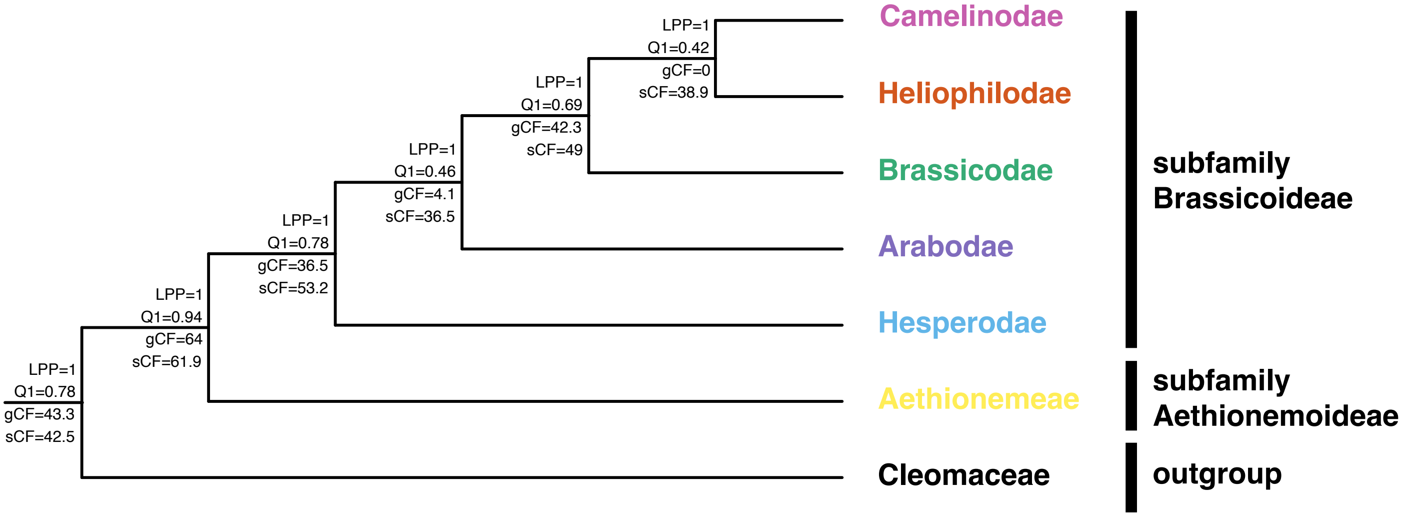


**Fig. S18. Phylogenetic relationships among the main Brassicaceae lineages**
Simplified backbone phylogeny summarising the main evolutionary lineages of Brassicaceae. The tree includes one representative sample from tribe Aethionemeae, one representative from each of the five core Brassicaceae supertribes, and a representative of the sister family Cleomaceae as outgroup. Branch lengths are arbitrary and shown only to illustrate topology. Node labels report ASTRAL local posterior probabilities (LPP), quartet support values (Q1), gene concordance factors (gCF), and site concordance factors (sCF) extracted from the full Brassicaceae Tree of Life reconstruction.

**Table S1. Crown and stem node ages and phylogenetic support values for main Brassicaceae lineages inferred from the calibrated Brassicaceae Tree of Life (BrassiToL)**

Estimated crown and stem ages, together with associated phylogenetic support metrics, for the Brassicaceae family, supertribes, and tribes identified in the time-calibrated Brassicaceae Tree of Life inferred using LSD2 dating on the fixed-topology ASTRAL species tree. Reported support metrics include ASTRAL quartet support (Q1), local posterior probability (LPP), gene concordance factor (gCF), and site concordance factor (sCF) for both crown and stem nodes where applicable. Crown ages correspond to the most recent common ancestor of all included representatives of a clade. For monospecific tribes and tribes represented here by a single species, only stem age can be calculated by defintion.

| rank | clade | n_tips | crown_node | crown_age_Ma | crown_q1 | crown_pp1 | crown_gCF | crown_sCF | stem_node | stem_age_Ma | stem_q1 | stem_pp1 | stem_gCF | stem_sCF |
|---|---|---|---|---|---|---|---|---|---|---|---|---|---|---|
| FAMILY | Brassicaceae | 2934 | 3104 | 36.9279 | 0.938765 | 1 | 64 | 61.9 | 3103 | 54.7924 | 0.782295 | 1 | 43.3 | 42.5 |
| SUPERTRIBE | Arabodae | 670 | 3106 | 26.05644 | 0.463397 | 1 | 4.09 | 36.5 | 3105 | 27.76377 | 0.779977 | 1 | 36.5 | 53.2 |
| SUPERTRIBE | Brassicodae | 595 | 3379 | 24.29342 | 0.461563 | 1 | 0.102 | 37.5 | 3108 | 26.05644 | 0.934275 | 1 | 86.2 | 79.9 |
| SUPERTRIBE | Camelinodae | 1081 | 3977 | 18.93836 | 0.654239 | 1 | 1.94 | 46 | 3976 | 20.91223 | 0.53461 | 1 | 1.46 | 40.4 |
| SUPERTRIBE | Heliophilodae | 273 | 3105 | 27.76377 | 0.779977 | 1 | 36.5 | 53.2 | 3104 | 36.9279 | 0.938765 | 1 | 64 | 61.9 |
| SUPERTRIBE | Hesperodae | 255 | 5732 | 19.83051 | 0.936458 | 1 | 33.9 | 59.1 | 3105 | 27.76377 | 0.779977 | 1 | 36.5 | 53.2 |
| TRIBE | Aethionemeae | 51 | 5987 | 16.6263 | 0.981142 | 1 | 70.2 | 75.2 | 3104 | 36.9279 | 0.938765 | 1 | 64 | 61.9 |
| TRIBE | Alysseae | 209 | 5074 | 17.34234 | 0.707697 | 1 | 36.8 | 61.8 | 5073 | 17.34234 | 0.582913 | 1 | 21 | 55.3 |
| TRIBE | Alyssopsideae | 4 | 4186 | 5.9314041 | 0.774227 | 1 | 60.7 | 69 | 4185 | 9.9509741 | 0.383232 | 0.972588 | 15.8 | 32 |
| TRIBE | Anastaticeae | 69 | 3105 | 27.76377 | 0.779977 | 1 | 36.5 | 53.2 | 3104 | 36.9279 | 0.938765 | 1 | 64 | 61.9 |
| TRIBE | Anchonieae | 63 | 5864 | 10.954835 | 0.829217 | 1 | 41.5 | 62.3 | 5863 | 15.603015 | 0.661619 | 1 | 26.6 | 45.9 |
| TRIBE | Aphragmeae | 1 | 868 | NA | NA | NA | NA | NA | 3379 | 24.29342 | 0.461563 | 1 | 0.102 | 37.5 |
| TRIBE | Arabideae | 451 | 5282 | 16.35684 | 0.818221 | 1 | 24 | 52.3 | 5071 | 23.4659 | 0.490769 | 1 | 14.6 | 45.5 |
| TRIBE | Arabidopsideae | 3 | 4313 | 8.7785451 | 0.919084 | 1 | 77.7 | 68.7 | 3981 | 16.9788951 | 0.426345 | 0.999935 | 1.43 | 36.9 |
| TRIBE | Asperuginoideae | 1 | 2176 | NA | NA | NA | NA | NA | 5072 | 20.74926 | 0.5588 | 1 | 8.93 | 37.2 |
| TRIBE | Asteae | 2 | 3276 | 18.964076 | 0.568441 | 1 | 15 | 48.4 | 3275 | 21.452616 | 0.392828 | 0.971524 | 1.27 | 32.4 |
| TRIBE | Biscutelleae | 8 | 3107 | 26.05644 | 0.688802 | 1 | 42.3 | 49 | 3106 | 26.05644 | 0.463397 | 1 | 4.09 | 36.5 |
| TRIBE | Boechereae | 127 | 3989 | 4.8028041 | 0.731507 | 1 | 19.8 | 52.6 | 3988 | 6.0812741 | 0.629997 | 1 | 21.6 | 54.4 |
| TRIBE | Brassiceae | 248 | 3578 | 15.484749 | 0.388805 | 0.953439 | 0.995 | 25.9 | 3387 | 15.484749 | 0.31178 | 0.002785 | 0.574 | 28.3 |
| TRIBE | Buniadeae | 1 | 2821 | NA | NA | NA | NA | NA | 5863 | 15.603015 | 0.661619 | 1 | 26.6 | 45.9 |
| TRIBE | Calepineae | 3 | 3867 | 9.642781 | 0.913328 | 1 | 77.2 | 75.4 | 3383 | 19.948981 | 0.526337 | 1 | 2.32 | 38.6 |
| TRIBE | Camelineae | 4 | 4123 | 6.6557741 | 0.813676 | 1 | 58.1 | 60.3 | 3986 | 9.7003741 | 0.489023 | 1 | 1.24 | 42.6 |
| TRIBE | Cardamineae | 235 | 4827 | 7.66896 | 0.933851 | 1 | 33.9 | 73.7 | 3977 | 18.93836 | 0.654239 | 1 | 1.94 | 46 |
| TRIBE | Chamireae | 1 | 99 | NA | NA | NA | NA | NA | 3112 | 24.252441 | 0.385572 | 0.995603 | 6.24 | 36 |
| TRIBE | Chorisporeae | 57 | 5929 | 8.80052 | 0.534275 | 1 | 28.9 | 56.1 | 5928 | 10.97639 | 0.798063 | 1 | 61.2 | 60.3 |
| TRIBE | Cochlearieae | 2 | 3974 | 14.200824 | 0.368754 | 0.947328 | 0.475 | 35 | 3380 | 23.748114 | 0.390496 | 0.955881 | 0.452 | 38.7 |
| TRIBE | Coluteocarpeae | 102 | 3871 | 13.657955 | 0.49974 | 1 | 14.2 | 38.6 | 3870 | 13.657955 | 0.684544 | 1 | 26.1 | 44 |
| TRIBE | Conringieae | 1 | 862 | NA | NA | NA | NA | NA | 3869 | 14.360131 | 0.911642 | 1 | 45.4 | 66.3 |
| TRIBE | Cremolobeae | 19 | 3240 | 16.306028 | 0.495549 | 1 | 6.14 | 34.7 | 3239 | 16.306028 | 0.360208 | 0.732934 | 1.15 | 35.3 |
| TRIBE | Crucihimalayeae | 3 | 4119 | 6.0261341 | 0.80818 | 1 | 44.4 | 61 | 3987 | 7.8854641 | 0.60746 | 1 | 3.36 | 60.8 |
| TRIBE | Descurainieae | 44 | 4544 | 15.9467021 | 0.747333 | 1 | 19.4 | 45.3 | 4542 | 17.3186221 | 0.351648 | 0.454137 | 7.28 | 48.3 |
| TRIBE | Dontostemoneae | 3 | 5985 | 8.21309 | 0.869259 | 1 | 70.7 | 67.8 | 5927 | 15.53781 | 0.793315 | 1 | 47 | 54.2 |
| TRIBE | Erysimeae | 227 | 4316 | 5.7640721 | 0.831867 | 1 | 37.7 | 74 | 4315 | 10.1019921 | 0.919775 | 1 | 47.5 | 71.8 |
| TRIBE | Euclidieae | 129 | 5735 | 16.208315 | 0.370823 | 0.883003 | 21.4 | 36.7 | 5734 | 18.049845 | 0.472988 | 1 | 9.12 | 37.8 |
| TRIBE | Eudemeae | 26 | 3214 | 13.492128 | 0.733004 | 1 | 30.1 | 55.8 | 3213 | 18.179418 | 0.900233 | 1 | 69 | 90.8 |
| TRIBE | Eutremeae | 1 | 718 | NA | NA | NA | NA | NA | 3830 | 16.940711 | 0.538334 | 1 | 18 | 41.3 |
| TRIBE | Fourraeeae | 2 | 3577 | 5.845451 | 0.823031 | 1 | 12.6 | 71.5 | 3388 | 14.526671 | 0.62066 | 1 | 2.5 | 62 |
| TRIBE | Halimolobeae | 5 | 4115 | 4.4779241 | 0.782177 | 1 | 33.4 | 61.3 | 3988 | 6.0812741 | 0.629997 | 1 | 21.6 | 54.4 |
| TRIBE | Heliophileae | 98 | 3113 | 16.558931 | 0.937917 | 1 | 88.1 | 91.6 | 3112 | 24.252441 | 0.385572 | 0.995603 | 6.24 | 36 |
| TRIBE | Hemilophieae | 2 | 4127 | 7.1030941 | 0.426078 | 0.999998 | 1.03 | 34.7 | 4126 | 9.1551341 | 0.435692 | 0.989374 | 4.25 | 56.7 |
| TRIBE | Hesperideae | 1 | 2822 | NA | NA | NA | NA | NA | 5733 | 18.35521 | 0.476393 | 1 | 10.1 | 39.1 |
| TRIBE | Hillielleae | 1 | 2175 | NA | NA | NA | NA | NA | 5073 | 17.34234 | 0.582913 | 1 | 21 | 55.3 |
| TRIBE | Iberideae | 25 | 3354 | 11.647114 | 0.947002 | 1 | 62.3 | 82.2 | 3286 | 25.491314 | 0.434542 | 0.999961 | 20.1 | 40.9 |
| TRIBE | Iberideae_II | 3 | 3278 | 13.042558 | 0.941507 | 1 | 89.1 | 80.2 | 3277 | 24.875958 | 0.615284 | 1 | 21.7 | 51.8 |
| TRIBE | Isatideae | 6 | 3825 | 8.272428 | 0.952397 | 1 | 78.5 | 76.2 | 3386 | 15.699528 | 0.525672 | 1 | 4.72 | 42.4 |
| TRIBE | Kernereae | 3 | 3972 | 8.860514 | 0.92754 | 1 | 76.7 | 79.6 | 3381 | 21.987314 | 0.459746 | 1 | 0.392 | 38.8 |
| TRIBE | Lepidieae | 240 | 4588 | 11.1107321 | 0.680146 | 1 | 24.3 | 48.2 | 4587 | 11.1107321 | 0.87538 | 1 | 30.8 | 58.4 |
| TRIBE | Malcolmieae | 1 | 1204 | NA | NA | NA | NA | NA | 4315 | 10.1019921 | 0.919775 | 1 | 47.5 | 71.8 |
| TRIBE | Megacarpaeeae | 2 | 3378 | 16.667054 | 0.432598 | 1 | 0 | 35 | 3285 | 25.491314 | 0.387658 | 0.995525 | 2.17 | 33.7 |
| TRIBE | Microlepidieae | 56 | 3984 | 11.2386161 | 0.473699 | 1 | 1.13 | 48.9 | 3983 | 13.9857461 | 0.489544 | 1 | 0.856 | 49 |
| TRIBE | Notothlaspideae | 1 | 100 | NA | NA | NA | NA | NA | 3111 | 24.252441 | 0.513564 | 1 | 5.11 | 39 |
| TRIBE | Oreophytoneae | 2 | 4189 | 1.6518361 | 0.941707 | 1 | 86.5 | 88.4 | 4182 | 11.6913361 | 0.604278 | 1 | 13.4 | 44.8 |
| TRIBE | Physarieae | 124 | 4190 | 15.6234161 | 0.930467 | 1 | 84.5 | 80.2 | 3982 | 16.6751461 | 0.395042 | 0.998576 | 1.24 | 35.6 |
| TRIBE | Plagiolobeae | 1 | 861 | NA | NA | NA | NA | NA | 3870 | 13.657955 | 0.684544 | 1 | 26.1 | 44 |
| TRIBE | Schizopetaleae | 16 | 3258 | 13.173828 | 0.377429 | 0.896891 | 11.1 | 29.9 | 3239 | 16.306028 | 0.360208 | 0.732934 | 1.15 | 35.3 |
| TRIBE | Schrenkielleae | 1 | 717 | NA | NA | NA | NA | NA | 3385 | 16.559151 | 0.810682 | 1 | 12.5 | 50.1 |
| TRIBE | Shehbazieae | 1 | 2823 | NA | NA | NA | NA | NA | 5928 | 10.97639 | 0.798063 | 1 | 61.2 | 60.3 |
| TRIBE | Sisymbrieae | 5 | 3573 | 8.544871 | 0.837835 | 1 | 35.8 | 50.9 | 3390 | 10.419361 | 0.434394 | 0.999988 | 0.414 | 41 |
| TRIBE | Smelowskieae | 1 | 1433 | NA | NA | NA | NA | NA | 4543 | 12.0620221 | 0.498198 | 1 | 1.79 | 42.3 |
| TRIBE | Stevenieae | 9 | 5061 | 12.71973 | 0.873783 | 1 | 81.3 | 84.9 | 3976 | 20.91223 | 0.53461 | 1 | 1.46 | 40.4 |
| TRIBE | Subularieae | 2 | 3210 | 24.875958 | 0.368848 | 0.835647 | 2.52 | 30.8 | 3110 | 25.679441 | 0.970209 | 1 | 91.6 | 85.9 |
| TRIBE | Thelypodieae | 184 | 3389 | 11.538441 | 0.357796 | 0.289647 | 0.517 | 41.8 | 3388 | 14.526671 | 0.62066 | 1 | 2.5 | 62 |
| TRIBE | Thlaspideae | 37 | 3831 | 13.561761 | 0.925576 | 1 | 57.1 | 69.7 | 3830 | 16.940711 | 0.538334 | 1 | 18 | 41.3 |
| TRIBE | Turritideae | 1 | 1068 | NA | NA | NA | NA | NA | 4183 | 10.3894461 | 0.455011 | 1 | 11.7 | 37.6 |
| TRIBE | Yinshanieae | 1 | 1432 | NA | NA | NA | NA | NA | 4543 | 12.0620221 | 0.498198 | 1 | 1.79 | 42.3 |

**Table S2. Summary statistics for phylogenetic signal in woodiness across the Brassicaceae tree**

Summary of phylogenetic signal analyses for binary growth form (herbaceous vs woody) across the coded Brassicaceae phylogeny. Pagel's λ was estimated by comparing an equal-rates discrete model with and without a λ transformation of branch lengths; the table reports the estimated λ, likelihood-ratio test statistic, associated p value, and interpretation. Fritz & Purvis' D was used as a complementary measure of phylogenetic structure for binary traits; the table reports the estimated D value together with tests against random trait distribution (D = 1) and Brownian-like phylogenetic structure (D = 0). Lower D values indicate stronger phylogenetic clustering of growth form across the tree. n_tips_coded gives the number of sampled tips with non-missing growth-form coding included in each analysis.

| metric | n_tips_coded | estimate | test_statistic | df | p_value_primary | p_value_secondary | primary_label | secondary_label | interpretation |
|---|---|---|---|---|---|---|---|---|---|
| Pagel_lambda | 3076 | 0.9449 | 82.4055 | 1 | 1.11E-19 | NA | LRT_lambda_vs_no_transform | NA | Trait shows strong phylogenetic dependence. |
| Fritz_Purvis_D | 3076 | 0.2023 | NA | NA | 0 | 0 | P_vs_random_D1 | P_vs_brownian_D0 | D indicates a phylogenetically clumped distribution; distribution differs from random expectation (D = 1); distribution is not distinguishable from Brownian expectation (D = 0). |

**Table S3. Mk model comparison for growth form evolution in Brassicaceae**
Comparison of equal-rates (ER) and all-rates-different (ARD) Mk models for binary growth form (herbaceous vs woody). Model support is assessed using log-likelihood, AIC/AICc, ΔIC, and Akaike weights. The best-supported model was used in downstream ancestral state reconstruction and SIMMAP analyses.

| model | logLik | k | AIC | AICc | deltaIC | weight |
|---|---|---|---|---|---|---|
| ARD | -885.72 | 2 | 1775.44 | 1775.44 | 0 | 1 |
| ER | -923.40 | 1 | 1848.80 | 1848.80 | 73.36 | 1.17E-16 |

**Table S4. Frequency and timing of woodiness gains and losses inferred from stochastic character mapping**

Summary of transitions between herbaceous and woody growth forms inferred from stochastic character mapping (SIMMAP) under the best-fitting Mk model, the ARD model. Values represent median estimates and 95% intervals across replicate simulations for the number of transitions (n_events) and their timing (in millions of years before present), reported separately for herbaceous-to-woody (H>W) and woody-to-herbaceous (W>H) shifts. Proportions indicate the relative contribution of each transition type to the total number of inferred events.

| direction | n_events_median | n_events_q025 | n_events_q975 | proportion_median | age_median | age_q025 | age_q975 |
|---|---|---|---|---|---|---|---|
| H→W | 231 | 213 | 250 | 0.5703 | 2.704 | 0.082 | 48.547 |
| W→H | 176 | 149 | 204 | 0.4297 | 2.608 | 0.083 | 54.103 |

**Table S5. Summary of confident growth-form shifts on the Brassicaceae–Cleomaceae tree**
Summary of confident shifts between herbaceous and woody growth forms identified on the reduced Brassicaceae–Cleomaceae tree. Confident shifts are defined as edge-specific transitions supported both by ancestral state reconstruction (ASR) and by directionally concordant stochastic character mapping (SIMMAP). Shifts are grouped by whether they occur on internal or terminal edges and by direction (H>W or W>H). For each category, the table reports the number of inferred shifts and summary statistics for directional SIMMAP support (p_event_dir).

| subclass | node_dir | n_edges | support_metric | support_median | support_q025 | support_q975 |
|---|---|---|---|---|---|---|
| Confident internal shift | H→W | 42 | p_event_dir | 0.779 | 0.199 | 1.000 |
| Confident internal shift | W→H | 12 | p_event_dir | 0.772 | 0.316 | 0.939 |
| Confident terminal shift | H→W | 101 | p_event_dir | 0.916 | 0.653 | 1.000 |
| Confident terminal shift | W→H | 63 | p_event_dir | 0.942 | 0.547 | 1.000 |

#### Table S6. Timing summary of confident internal growth-form shifts

Summary of the inferred timing of confident internal shifts between herbaceous and woody growth forms on the Brassicaceae–Cleomaceae tree. Only internal shifts are included, as these are less sensitive than terminal shifts to potential tip-level error in sample identification or growth-form coding. For each numbered shift, the table reports the inferred direction, clade label, clade size, and the median and 95% interval of shift ages estimated from directionally concordant SIMMAP events projected onto the corresponding edge. All times and ages in millions of years.

| shift_id | shift_label2 | shift_name_display | shift_name_display_axis | node_dir | family | tribe | supertribe | group | child_node | n_tribe_tips | n_events | age_med | age_lo | age_hi | tribe_crown_age | lag_from_tribe_crown |
|---|---|---|---|---|---|---|---|---|---|---|---|---|---|---|---|---|
| NA | C1 Andinocleome | Andinocleome clade | Andinocleome | H>W | Cleomaceae | NA | NA | Cleomaceae | 6007 | NA | 287 | 8.229 | 4.757 | 11.785 | NA | NA |
| NA | C2 Melidiscus | Melidiscus clade | Melidiscus | H>W | Cleomaceae | NA | NA | Cleomaceae | 6013 | NA | 468 | 11.576 | 4.832 | 21.067 | NA | NA |
| 1 | 1 Aethionema I | Aethionema clade I | Aethionema I | H>W | Brassicaceae | Aethionemeae | NA | Aethionemeae | 5927 | 51 | 423 | 24.183 | 17.532 | 37.458 | 17.091 | -7.092 |
| 2 | 2 Aethionema II | Aethionema clade II | Aethionema II | W>H | Brassicaceae | Aethionemeae | NA | Aethionemeae | 5970 | 51 | 409 | 8.166 | 5.686 | 10.203 | NA | NA |
| 3 | 3 Aethionema III | Aethionema clade III | Aethionema III | W>H | Brassicaceae | Aethionemeae | NA | Aethionemeae | 5955 | 51 | 395 | 2.640 | 0.838 | 4.491 | NA | NA |
| 4 | 4 Aethionema IV | Aethionema clade IV | Aethionema IV | W>H | Brassicaceae | Aethionemeae | NA | Aethionemeae | 5966 | 51 | 439 | 2.617 | 1.842 | 3.323 | NA | NA |
| 5 | 5 Parrya | Parrya clade | Parrya | H>W | Brassicaceae | Chorisporeae | Hesperodae | Hesperodae | 5884 | 57 | 460 | 2.141 | 1.582 | 2.694 | 9.046 | 6.905 |
| 6 | 6 Matthiola | Matthiola clade | Matthiola | H>W | Brassicaceae | Anchonieae | Hesperodae | Hesperodae | 5815 | 63 | 334 | 3.811 | 2.739 | 4.860 | 11.261 | 7.450 |
| 7 | 7 Rhammatophyllum | Rhammatophyllum clade | Rhammatophyllum | H>W | Brassicaceae | Euclidieae | Hesperodae | Hesperodae | 5768 | 129 | 346 | 7.340 | 6.454 | 8.190 | 16.676 | 9.336 |
| 8 | 8 Hormathophylla | Hormathophylla clade | Hormathophylla | H>W | Brassicaceae | Alysseae | Arabodae | Arabodae | 5046 | 208 | 475 | 8.649 | 4.844 | 14.050 | 17.832 | 9.184 |
| 9 | 9 Odontarrhena I | Odontarrhena clade I | Odontarrhena I | H>W | Brassicaceae | Alysseae | Arabodae | Arabodae | 5147 | 208 | 478 | 6.883 | 4.707 | 9.987 | 17.832 | 10.950 |
| 10 | 10 Odontarrhena II | Odontarrhena clade II | Odontarrhena II | W>H | Brassicaceae | Alysseae | Arabodae | Arabodae | 5204 | 208 | 379 | 2.096 | 1.387 | 2.696 | NA | NA |
| 11 | 11 Odontarrhena III | Odontarrhena clade III | Odontarrhena III | W>H | Brassicaceae | Alysseae | Arabodae | Arabodae | 5169 | 208 | 249 | 1.620 | 1.228 | 1.942 | NA | NA |
| 12 | 12 Odontarrhena IV | Odontarrhena clade IV | Odontarrhena IV | W>H | Brassicaceae | Alysseae | Arabodae | Arabodae | 5195 | 208 | 396 | 1.780 | 1.492 | 2.167 | NA | NA |
| 13 | 13 Odontarrhena V | Odontarrhena clade V | Odontarrhena V | W>H | Brassicaceae | Alysseae | Arabodae | Arabodae | 5191 | 208 | 468 | 1.089 | 0.774 | 1.442 | NA | NA |
| 14 | 14 Alyssum I | Alyssum clade I | Alyssum I | W>H | Brassicaceae | Alysseae | Arabodae | Arabodae | 5064 | 208 | 229 | 3.841 | 3.552 | 4.199 | NA | NA |
| 15 | 15 Alyssum II | Alyssum clade II | Alyssum II | H>W | Brassicaceae | Alysseae | Arabodae | Arabodae | 5062 | 208 | 100 | 6.291 | 4.723 | 7.747 | 17.832 | 11.541 |
| 16 | 16 Alyssum III | Alyssum clade III | Alyssum III | W>H | Brassicaceae | Alysseae | Arabodae | Arabodae | 5072 | 208 | 173 | 5.269 | 4.964 | 5.566 | NA | NA |
| 17 | 17 Alyssum IV | Alyssum clade IV | Alyssum IV | H>W | Brassicaceae | Alysseae | Arabodae | Arabodae | 5122 | 208 | 400 | 3.773 | 3.302 | 4.260 | 17.832 | 14.060 |
| 18 | 18 Draba I | Draba clade I | Draba I | H>W | Brassicaceae | Arabideae | Arabodae | Arabodae | 5341 | 450 | 446 | 1.198 | 1.194 | 1.203 | 16.827 | 15.629 |
| 19 | 19 Draba II | Draba clade II | Draba II | H>W | Brassicaceae | Arabideae | Arabodae | Arabodae | 5338 | 450 | 450 | 1.015 | 0.994 | 1.036 | 16.827 | 15.813 |
| 20 | 20 Draba III | Draba clade III | Draba III | H>W | Brassicaceae | Arabideae | Arabodae | Arabodae | 5334 | 450 | 129 | 0.475 | 0.451 | 0.500 | 16.827 | 16.353 |
| 21 | 21 Draba IV | Draba clade IV | Draba IV | H>W | Brassicaceae | Arabideae | Arabodae | Arabodae | 5320 | 450 | 500 | 0.601 | 0.472 | 0.715 | 16.827 | 16.227 |
| 22 | 22 Iberis I | Iberis clade I | Iberis I | H>W | Brassicaceae | Iberideae | Heliophilodae | Heliophilodae | 3327 | 25 | 215 | 8.523 | 5.546 | 11.684 | 11.967 | 3.444 |
| 23 | 23 Iberis II | Iberis clade II | Iberis II | W>H | Brassicaceae | Iberideae | Heliophilodae | Heliophilodae | 3329 | 25 | 171 | 2.616 | 2.257 | 3.004 | NA | NA |
| 24 | 24 Farsetia | Farsetia clade | Farsetia | H>W | Brassicaceae | Anastaticeae | Heliophilodae | Heliophilodae | 3242 | 68 | 292 | 16.724 | 13.450 | 21.748 | 28.554 | 11.830 |
| 25 | 25 Diceratella | Diceratella clade | Diceratella | H>W | Brassicaceae | Anastaticeae | Heliophilodae | Heliophilodae | 3268 | 68 | 118 | 20.091 | 18.563 | 21.809 | 28.554 | 8.463 |
| 26 | 26 Cremolobus | Cremolobus clade | Cremolobus | H>W | Brassicaceae | Cremolobeae | Heliophilodae | Heliophilodae | 3195 | 19 | 314 | 12.704 | 9.144 | 16.576 | 16.755 | 4.051 |
| 27 | 27 Heliophila | Heliophila clade | Heliophila | H>W | Brassicaceae | Heliophileae | Heliophilodae | Heliophilodae | 3067 | 98 | 309 | 15.311 | 13.556 | 16.896 | 17.018 | 1.707 |
| 28 | 28 Eunomia | Eunomia clade | Eunomia | H>W | Brassicaceae | Coluteocarpeae | Brassicodae | Brassicodae | 3921 | 102 | 218 | 3.630 | 3.181 | 4.123 | 14.035 | 10.404 |
| 29 | 29 Mostacillastrum I | Mostacillastrum clade I | Mostacillastrum I | H>W | Brassicaceae | Thelypodieae | Brassicodae | Brassicodae | 3497 | 184 | 377 | 3.494 | 3.243 | 3.789 | 11.886 | 8.392 |
| 30 | 30 Neuontobotrys | Neuontobotrys clade | Neuontobotrys | H>W | Brassicaceae | Thelypodieae | Brassicodae | Brassicodae | 3487 | 184 | 463 | 1.696 | 1.397 | 2.021 | 11.886 | 10.190 |
| 31 | 31 Mostacillastrum II | Mostacillastrum clade II | Mostacillastrum II | H>W | Brassicaceae | Thelypodieae | Brassicodae | Brassicodae | 3467 | 184 | 456 | 0.994 | 0.830 | 1.186 | 11.886 | 10.892 |
| 32 | 32 Hesperidanthus | Hesperidanthus clade | Hesperidanthus | H>W | Brassicaceae | Thelypodieae | Brassicodae | Brassicodae | 3436 | 184 | 472 | 1.545 | 1.065 | 1.985 | 11.886 | 10.341 |
| 33 | 33 Vella | Vella clade | Vella | H>W | Brassicaceae | Brassiceae | Brassicodae | Brassicodae | 3766 | 248 | 312 | 7.145 | 5.619 | 8.857 | 15.975 | 8.830 |
| 34 | 34 Douepea | Douepea clade | Douepea | H>W | Brassicaceae | Brassiceae | Brassicodae | Brassicodae | 3757 | 248 | 311 | 6.598 | 5.145 | 7.978 | 15.975 | 9.377 |
| 35 | 35 Crambe | Crambe clade | Crambe | H>W | Brassicaceae | Brassiceae | Brassicodae | Brassicodae | 3714 | 248 | 433 | 6.597 | 4.275 | 9.205 | 15.975 | 9.378 |
| 36 | 36 Moricandia | Moricandia clade | Moricandia | H>W | Brassicaceae | Brassiceae | Brassicodae | Brassicodae | 3686 | 248 | 88 | 9.221 | 8.284 | 10.164 | 15.975 | 6.754 |
| 37 | 37 Diplotaxis | Diplotaxis clade | Diplotaxis | H>W | Brassicaceae | Brassiceae | Brassicodae | Brassicodae | 3565 | 248 | 271 | 9.578 | 7.883 | 11.740 | 15.975 | 6.397 |
| 38 | 38 Hemicrambe | Hemicrambe clade | Hemicrambe | H>W | Brassicaceae | Brassiceae | Brassicodae | Brassicodae | 3684 | 248 | 275 | 2.674 | 2.027 | 3.332 | 15.975 | 13.301 |
| 39 | 39 Brassica I | Brassica clade I | Brassica I | H>W | Brassicaceae | Brassiceae | Brassicodae | Brassicodae | 3660 | 248 | 257 | 3.519 | 3.393 | 3.635 | 15.975 | 12.457 |
| 40 | 40 Brassica II | Brassica clade II | Brassica II | W>H | Brassicaceae | Brassiceae | Brassicodae | Brassicodae | 3664 | 248 | 153 | 2.562 | 2.394 | 2.716 | NA | NA |
| 41 | 41 Sinapidendron | Sinapidendron clade | Sinapidendron | H>W | Brassicaceae | Brassiceae | Brassicodae | Brassicodae | 3616 | 248 | 409 | 6.670 | 4.190 | 9.144 | 15.975 | 9.305 |
| 42 | 42 Stevenia | Stevenia clade | Stevenia | H>W | Brassicaceae | Stevenieae | Arabodae | Arabodae | 5008 | 9 | 438 | 5.201 | 1.548 | 10.201 | 13.078 | 7.877 |
| 43 | 43 Lepidium I | Lepidium clade I | Lepidium I | H>W | Brassicaceae | Lepidieae | Camelinodae | Camelinodae | 4754 | 239 | 378 | 8.197 | 5.367 | 11.228 | 11.418 | 3.221 |
| 44 | 44 Lepidium II | Lepidium clade II | Lepidium II | W>H | Brassicaceae | Lepidieae | Camelinodae | Camelinodae | 4765 | 239 | 470 | 2.859 | 1.769 | 3.871 | NA | NA |
| 45 | 45 Lepidium III | Lepidium clade III | Lepidium III | H>W | Brassicaceae | Lepidieae | Camelinodae | Camelinodae | 4572 | 239 | 496 | 1.009 | 0.706 | 1.336 | 11.418 | 10.409 |
| 46 | 46 Lepidium IV | Lepidium clade IV | Lepidium IV | H>W | Brassicaceae | Lepidieae | Camelinodae | Camelinodae | 4648 | 239 | 477 | 1.203 | 0.937 | 1.451 | 11.418 | 10.215 |
| 47 | 47 Descurainia | Descurainia clade | Descurainia | H>W | Brassicaceae | Descurainieae | Camelinodae | Camelinodae | 4518 | 44 | 432 | 2.470 | 1.765 | 3.093 | 16.391 | 13.922 |
| 48 | 48 Erysimum I | Erysimum clade I | Erysimum I | H>W | Brassicaceae | Erysimeae | Camelinodae | Camelinodae | 4440 | 227 | 499 | 1.851 | 1.351 | 2.344 | 5.928 | 4.077 |
| 49 | 49 Erysimum II | Erysimum clade II | Erysimum II | H>W | Brassicaceae | Erysimeae | Camelinodae | Camelinodae | 4353 | 227 | 500 | 1.169 | 0.947 | 1.399 | 5.928 | 4.759 |
| 50 | 50 Lyrocarpa | Lyrocarpa clade | Lyrocarpa | H>W | Brassicaceae | Physarieae | Camelinodae | Camelinodae | 4148 | 124 | 273 | 2.411 | 1.432 | 3.396 | 16.059 | 13.647 |
| 51 | 51 Pachycladon | Pachycladon clade | Pachycladon | H>W | Brassicaceae | Microlepidieae | Camelinodae | Camelinodae | 4080 | 56 | 474 | 0.959 | 0.649 | 1.281 | 11.556 | 10.597 |
| 52 | 52 Arabidella | Arabidella clade | Arabidella | H>W | Brassicaceae | Microlepidieae | Camelinodae | Camelinodae | 4117 | 56 | 495 | 5.006 | 2.872 | 7.389 | 11.556 | 6.549 |

**Table S7. Lag times between tribe origin and the emergence of woody lineages**

Lag times (in million years, Ma) between the origin of tribes (crown nodes) and the inferred origin of derived woody clades (H>W shifts), based on highly confident internal transitions supported by both ancestral state reconstruction and stochastic character mapping. Summary statistics are shown for all shifts and after excluding Aethionemeae and Anastaticeae, the only two tribes in which inferred woodiness shifts predate lineage origin.

<table>
<tr><th>subset</th><th>n_shifts</th><th>lag_min</th><th>lag_q025</th><th>lag_median</th><th>lag_mean</th><th>lag_q975</th><th>lag_max</th><th>n_zero_lag</th><th>prop_zero_lag</th><th>interpretation</th></tr>
<tr><td>All H→W shifts</td><td>40</td><td>-7.0923</td><td>1.4872</td><td>9.3773</td><td>9.1785</td><td>16.2300</td><td>16.3526</td><td>0</td><td>0</td><td>Median lag = 9.38 Ma (range -7.09–16.35 Ma); 0.0% effectively zero lag (|lag| <= 0.25 Ma).</td></tr>
<tr><td>Excluding tribe Aethionemeae</td><td>39</td><td>1.7072</td><td>3.1455</td><td>9.3777</td><td>9.5957</td><td>16.2331</td><td>16.3526</td><td>0</td><td>0</td><td>Median lag = 9.38 Ma (range 1.71–16.35 Ma); 0.0% effectively zero lag (|lag| <= 0.25 Ma).</td></tr>
</table>

**Table S8. Temporal variation in the tempo and propensity of woodiness shifts across the Brassicaceae–Cleomaceae tree**

Median and 95% interval estimates of two complementary process-level summaries of growth-form evolution through time, based on all Brassicaceae–Cleomaceae (BC) projected SIMMAP events. For each 2-Myr time bin, tempo was calculated as the shift rate per unit lineage time (events · $Myr^{-1}$ lineage time), whereas propensity was calculated as the shift rate per time spent in the origin state (events · $Myr^{-1}$ spent in the origin state), providing a hazard-like measure of state-specific transition tendency. Estimates are shown separately for herbaceous-to-woody (H>W) and woody-to-herbaceous (W>H) transitions and are summarized across SIMMAP posterior replicates. Ages are given as time before present (Myr BP). This table summarizes broad temporal patterning across all BC-projected events, rather than only the confident internal shifts highlighted elsewhere.

| bin | bin_mid | bin_lo | bin_hi | node_dir | rate_med | rate_lo | rate_hi | prop_med | prop_lo | prop_hi |
|---|---|---|---|---|---|---|---|---|---|---|
| 1 | 1 | 0 | 2 | H>W | 0.01969895 | 0.01768886 | 0.02201559 | 0.02282627 | 0.02044410 | 0.02536209 |
| 1 | 1 | 0 | 2 | W>H | 0.01507573 | 0.01185957 | 0.01779439 | 0.11194348 | 0.09058786 | 0.13033589 |
| 2 | 3 | 2 | 4 | H>W | 0.01221018 | 0.00877607 | 0.01564430 | 0.01449651 | 0.01030584 | 0.01834780 |
| 2 | 3 | 2 | 4 | W>H | 0.01259175 | 0.00839450 | 0.01678900 | 0.08494132 | 0.05831786 | 0.11059121 |
| 3 | 5 | 4 | 6 | H>W | 0.01076281 | 0.00633107 | 0.01519456 | 0.01227779 | 0.00728003 | 0.01734258 |
| 3 | 5 | 4 | 6 | W>H | 0.01012971 | 0.00506485 | 0.01456145 | 0.08081984 | 0.04402190 | 0.11900832 |
| 4 | 7 | 6 | 8 | H>W | 0.01222135 | 0.00752083 | 0.01929564 | 0.01386091 | 0.00836805 | 0.02164577 |
| 4 | 7 | 6 | 8 | W>H | 0.00940104 | 0.00376042 | 0.01598177 | 0.08290288 | 0.03690230 | 0.13385782 |
| 5 | 9 | 8 | 10 | H>W | 0.01337367 | 0.00668683 | 0.02273523 | 0.01530734 | 0.00751546 | 0.02559620 |
| 5 | 9 | 8 | 10 | W>H | 0.00936157 | 0.00267473 | 0.01738577 | 0.08598486 | 0.02587106 | 0.15074546 |
| 6 | 11 | 10 | 12 | H>W | 0.01361076 | 0.00362954 | 0.02359199 | 0.01540280 | 0.00421186 | 0.02597792 |
| 6 | 11 | 10 | 12 | W>H | 0.00907384 | 0.00181477 | 0.01996245 | 0.09004788 | 0.01876147 | 0.17549475 |
| 7 | 13 | 12 | 14 | H>W | 0.01315796 | 0.00478471 | 0.02631593 | 0.01481795 | 0.00504610 | 0.02902114 |
| 7 | 13 | 12 | 14 | W>H | 0.00717707 | 0.00000000 | 0.01913886 | 0.08054615 | 0.00000000 | 0.19121941 |
| 8 | 15 | 14 | 16 | H>W | 0.01446930 | 0.00361733 | 0.02893861 | 0.01624835 | 0.00399588 | 0.03352875 |
| 8 | 15 | 14 | 16 | W>H | 0.00723465 | 0.00000000 | 0.02170396 | 0.07282737 | 0.00000000 | 0.22866478 |
| 9 | 17 | 16 | 18 | H>W | 0.01639256 | 0.00000000 | 0.03018963 | 0.01751995 | 0.00000000 | 0.03502858 |
| 9 | 17 | 16 | 18 | W>H | 0.01092837 | 0.00000000 | 0.02732094 | 0.08220584 | 0.00000000 | 0.24994482 |
| 10 | 19 | 18 | 20 | H>W | 0.01403916 | 0.00000000 | 0.03509790 | 0.01582983 | 0.00000000 | 0.04009828 |
| 10 | 19 | 18 | 20 | W>H | 0.00701958 | 0.00000000 | 0.03509790 | 0.07817944 | 0.00000000 | 0.27814668 |
| 11 | 21 | 20 | 22 | H>W | 0.01825210 | 0.00000000 | 0.04563026 | 0.01913767 | 0.00000000 | 0.04865785 |
| 11 | 21 | 20 | 22 | W>H | 0.00912605 | 0.00000000 | 0.03650421 | 0.09141202 | 0.00000000 | 0.43341421 |
| 12 | 23 | 22 | 24 | H>W | 0.01333060 | 0.00000000 | 0.05332239 | 0.01461389 | 0.00000000 | 0.05966217 |
| 12 | 23 | 22 | 24 | W>H | 0.00000000 | 0.00000000 | 0.03999180 | 0.00000000 | 0.00000000 | 0.54600745 |
| 13 | 25 | 24 | 26 | H>W | 0.01582935 | 0.00000000 | 0.04748805 | 0.01658715 | 0.00000000 | 0.05560887 |
| 13 | 25 | 24 | 26 | W>H | 0.00000000 | 0.00000000 | 0.04748805 | 0.00000000 | 0.00000000 | 0.73380556 |
| 14 | 27 | 26 | 28 | H>W | 0.00000000 | 0.00000000 | 0.05127767 | 0.00000000 | 0.00000000 | 0.06963972 |
| 14 | 27 | 26 | 28 | W>H | 0.00000000 | 0.00000000 | 0.05127767 | 0.00000000 | 0.00000000 | 1.38074167 |
| 15 | 29 | 28 | 30 | H>W | 0.00000000 | 0.00000000 | 0.06672179 | 0.00000000 | 0.00000000 | 0.07862267 |
| 15 | 29 | 28 | 30 | W>H | 0.00000000 | 0.00000000 | 0.06672179 | 0.00000000 | 0.00000000 | 1.43542528 |
| 16 | 31 | 30 | 32 | H>W | 0.00000000 | 0.00000000 | 0.08053002 | 0.00000000 | 0.00000000 | 0.09874706 |
| 16 | 31 | 30 | 32 | W>H | 0.00000000 | 0.00000000 | 0.08053002 | 0.00000000 | 0.00000000 | 1.43077719 |
| 17 | 33 | 32 | 34 | H>W | 0.00000000 | 0.00000000 | 0.05934413 | 0.00000000 | 0.00000000 | 0.12836593 |
| 17 | 33 | 32 | 34 | W>H | 0.00000000 | 0.00000000 | 0.11868826 | 0.00000000 | 0.00000000 | 2.02616477 |
| 18 | 35 | 34 | 36 | H>W | 0.00000000 | 0.00000000 | 0.11050408 | 0.00000000 | 0.00000000 | 0.15475398 |
| 18 | 35 | 34 | 36 | W>H | 0.00000000 | 0.00000000 | 0.11050408 | 0.00000000 | 0.00000000 | 1.90927859 |
| 19 | 37 | 36 | 38 | H>W | 0.00000000 | 0.00000000 | 0.12555212 | 0.00000000 | 0.00000000 | 0.16862368 |
| 19 | 37 | 36 | 38 | W>H | 0.00000000 | 0.00000000 | 0.12555212 | 0.00000000 | 0.00000000 | 1.25084734 |
| 20 | 39 | 38 | 40 | H>W | 0.00000000 | 0.00000000 | 0.16666667 | 0.00000000 | 0.00000000 | 0.22812572 |
| 20 | 39 | 38 | 40 | W>H | 0.00000000 | 0.00000000 | 0.16666667 | 0.00000000 | 0.00000000 | 3.21843486 |
| 21 | 41 | 40 | 42 | H>W | 0.00000000 | 0.00000000 | 0.16666667 | 0.00000000 | 0.00000000 | 0.21276364 |
| 21 | 41 | 40 | 42 | W>H | 0.00000000 | 0.00000000 | 0.16666667 | 0.00000000 | 0.00000000 | 1.93500754 |

**Table S9. Proportion of island endemics within woody and herbaceous species and distribution of woodiness across island endemics and mainland species**

(A) Counts of mainland and island-endemic species within herbaceous (H) and woody (W) growth forms for each sampling tier. The proportion of island endemics (p_island) is calculated within each growth-form category; (B) Counts of herbaceous (H) and woody (W) species across mainland and island-endemic taxa for each sampling tier (ge1, ge5, ge10). The proportion of woody species (p_woody) is calculated within each island category.

**(A)**

| tier | woody_state | mainland | island_endemic | total | p_mainland | p_island |
|---|---|---|---|---|---|---|
| ge1 | H | 3,025 | 57 | 3,082 | 0.982 | 0.018 |
| ge1 | W | 276 | 57 | 333 | 0.829 | 0.171 |
| ge5 | H | 2,317 | 41 | 2,358 | 0.983 | 0.017 |
| ge5 | W | 204 | 40 | 244 | 0.836 | 0.164 |
| ge10 | H | 1,937 | 28 | 1,965 | 0.986 | 0.014 |
| ge10 | W | 165 | 29 | 194 | 0.851 | 0.149 |

**(B)**

| tier | island_f | H | W | total | p_herbaceous | p_woody |
|---|---|---|---|---|---|---|
| ge1 | mainland | 3,025 | 276 | 3,301 | 0.916 | 0.084 |
| ge1 | island_endemic | 57 | 57 | 114 | 0.500 | 0.500 |
| ge5 | mainland | 2,317 | 204 | 2,521 | 0.919 | 0.081 |
| ge5 | island_endemic | 41 | 40 | 81 | 0.506 | 0.494 |
| ge10 | mainland | 1,937 | 165 | 2,102 | 0.922 | 0.078 |
| ge10 | island_endemic | 28 | 29 | 57 | 0.491 | 0.509 |

**Table S10. Summary statistics for woodiness–climate models across datasets and modelling frameworks**
Model-level summary statistics for all fitted generalized linear models (GLM full, GLM matched) and phylogenetic generalized linear models (phyloGLM matched) across ge tiers (ge1, ge5, ge10) and model formulations. Reported values include sample size (n), model formula, log-likelihood, AIC, and the phylogenetic signal parameter (α) for phyloGLM models.

| tier | model | framework | n | formula | logLik | AIC | alpha |
|---|---|---|---|---|---|---|---|
| ge1 | m_clim | glm_full | 3,412 | woody_bin ~ MCWD_med + BIO10_med + BIO6_med + elev_med | -963.85 | 1,937.71 | NA |
| ge1 | m_clim | glm_matched | 2,527 | woody_bin ~ MCWD_med + BIO10_med + BIO6_med + elev_med | -861.76 | 1,733.52 | NA |
| ge1 | m_clim | phyloglm_matched | 2,527 | woody_bin ~ MCWD_med + BIO10_med + BIO6_med + elev_med | -612.38 | 1,236.75 | 0.058 |
| ge1 | m_clim_breadth | glm_full | 2,896 | woody_bin ~ MCWD_med + BIO10_med + BIO6_med + elev_med + MCWD_mad + BIO10_mad + BIO6_mad + elev_mad | -791.25 | 1,600.49 | NA |
| ge1 | m_clim_breadth | glm_matched | 2,217 | woody_bin ~ MCWD_med + BIO10_med + BIO6_med + elev_med + MCWD_mad + BIO10_mad + BIO6_mad + elev_mad | -722.21 | 1,462.42 | NA |
| ge1 | m_clim_breadth | phyloglm_matched | 2,217 | woody_bin ~ MCWD_med + BIO10_med + BIO6_med + elev_med + MCWD_mad + BIO10_mad + BIO6_mad + elev_mad | -522.66 | 1,065.32 | 0.061 |
| ge1 | m_clim_island | glm_full | 3,412 | woody_bin ~ MCWD_med + BIO10_med + BIO6_med + elev_med + is_island | -928.14 | 1,868.27 | NA |
| ge1 | m_clim_island | glm_matched | 2,527 | woody_bin ~ MCWD_med + BIO10_med + BIO6_med + elev_med + is_island | -831.09 | 1,674.18 | NA |
| ge1 | m_clim_island | phyloglm_matched | 2,527 | woody_bin ~ MCWD_med + BIO10_med + BIO6_med + elev_med + is_island | -606.35 | 1,226.69 | 0.059 |
| ge1 | m_clim_mainland | glm_full | 3,299 | woody_bin ~ MCWD_med + BIO10_med + BIO6_med + elev_med | -864.01 | 1,738.03 | NA |
| ge1 | m_clim_mainland | glm_matched | 2,429 | woody_bin ~ MCWD_med + BIO10_med + BIO6_med + elev_med | -774.13 | 1,558.25 | NA |
| ge1 | m_clim_mainland | phyloglm_matched | 2,429 | woody_bin ~ MCWD_med + BIO10_med + BIO6_med + elev_med | -547.52 | 1,107.04 | 0.056 |
| ge5 | m_clim | glm_full | 2,601 | woody_bin ~ MCWD_med + BIO10_med + BIO6_med + elev_med | -713.12 | 1,436.24 | NA |
| ge5 | m_clim | glm_matched | 2,018 | woody_bin ~ MCWD_med + BIO10_med + BIO6_med + elev_med | -656.76 | 1,323.51 | NA |
| ge5 | m_clim | phyloglm_matched | 2,018 | woody_bin ~ MCWD_med + BIO10_med + BIO6_med + elev_med | -482.27 | 976.54 | 0.060 |
| ge5 | m_clim_breadth | glm_full | 2,601 | woody_bin ~ MCWD_med + BIO10_med + BIO6_med + elev_med + MCWD_mad + BIO10_mad + BIO6_mad + elev_mad | -703.19 | 1,424.38 | NA |
| ge5 | m_clim_breadth | glm_matched | 2,018 | woody_bin ~ MCWD_med + BIO10_med + BIO6_med + elev_med + MCWD_mad + BIO10_mad + BIO6_mad + elev_mad | -646.79 | 1,311.58 | NA |
| ge5 | m_clim_breadth | phyloglm_matched | 2,018 | woody_bin ~ MCWD_med + BIO10_med + BIO6_med + elev_med + MCWD_mad + BIO10_mad + BIO6_mad + elev_mad | -481.02 | 982.03 | 0.063 |
| ge5 | m_clim_island | glm_full | 2,601 | woody_bin ~ MCWD_med + BIO10_med + BIO6_med + elev_med + is_island | -685.94 | 1,383.88 | NA |
| ge5 | m_clim_island | glm_matched | 2,018 | woody_bin ~ MCWD_med + BIO10_med + BIO6_med + elev_med + is_island | -634.57 | 1,281.14 | NA |
| ge5 | m_clim_island | phyloglm_matched | 2,018 | woody_bin ~ MCWD_med + BIO10_med + BIO6_med + elev_med + is_island | -476.76 | 967.52 | 0.064 |
| ge5 | m_clim_mainland | glm_full | 2,520 | woody_bin ~ MCWD_med + BIO10_med + BIO6_med + elev_med | -639.08 | 1,288.16 | NA |
| ge5 | m_clim_mainland | glm_matched | 1,944 | woody_bin ~ MCWD_med + BIO10_med + BIO6_med + elev_med | -590.55 | 1,191.09 | NA |
| ge5 | m_clim_mainland | phyloglm_matched | 1,944 | woody_bin ~ MCWD_med + BIO10_med + BIO6_med + elev_med | -429.91 | 871.83 | 0.055 |
| ge10 | m_clim | glm_full | 2,158 | woody_bin ~ MCWD_med + BIO10_med + BIO6_med + elev_med | -582.12 | 1,174.24 | NA |
| ge10 | m_clim | glm_matched | 1,699 | woody_bin ~ MCWD_med + BIO10_med + BIO6_med + elev_med | -538.80 | 1,087.61 | NA |
| ge10 | m_clim | phyloglm_matched | 1,699 | woody_bin ~ MCWD_med + BIO10_med + BIO6_med + elev_med | -406.14 | 824.29 | 0.057 |
| ge10 | m_clim_breadth | glm_full | 2,158 | woody_bin ~ MCWD_med + BIO10_med + BIO6_med + elev_med + MCWD_mad + BIO10_mad + BIO6_mad + elev_mad | -572.73 | 1,163.46 | NA |
| ge10 | m_clim_breadth | glm_matched | 1,699 | woody_bin ~ MCWD_med + BIO10_med + BIO6_med + elev_med + MCWD_mad + BIO10_mad + BIO6_mad + elev_mad | -529.82 | 1,077.64 | NA |
| ge10 | m_clim_breadth | phyloglm_matched | 1,699 | woody_bin ~ MCWD_med + BIO10_med + BIO6_med + elev_med + MCWD_mad + BIO10_mad + BIO6_mad + elev_mad | -403.91 | 827.81 | 0.061 |
| ge10 | m_clim_island | glm_full | 2,158 | woody_bin ~ MCWD_med + BIO10_med + BIO6_med + elev_med + is_island | -558.41 | 1,128.82 | NA |
| ge10 | m_clim_island | glm_matched | 1,699 | woody_bin ~ MCWD_med + BIO10_med + BIO6_med + elev_med + is_island | -519.73 | 1,051.45 | NA |
| ge10 | m_clim_island | phyloglm_matched | 1,699 | woody_bin ~ MCWD_med + BIO10_med + BIO6_med + elev_med + is_island | -399.20 | 812.40 | 0.060 |
| ge10 | m_clim_mainland | glm_full | 2,101 | woody_bin ~ MCWD_med + BIO10_med + BIO6_med + elev_med | -525.11 | 1,060.22 | NA |
| ge10 | m_clim_mainland | glm_matched | 1,646 | woody_bin ~ MCWD_med + BIO10_med + BIO6_med + elev_med | -487.92 | 985.84 | NA |
| ge10 | m_clim_mainland | phyloglm_matched | 1,646 | woody_bin ~ MCWD_med + BIO10_med + BIO6_med + elev_med | -365.03 | 742.05 | 0.053 |

**Table S11. Effect of dataset restriction on model coefficients**
Comparison of standardized regression coefficients between models fitted to the full niche dataset and the matched niche–tree subset. For each predictor, the table reports coefficients, confidence intervals, odds ratios, and P-values for both models, as well as differences in coefficients (Δβ) and odds ratios (ΔOR). This quantifies the impact of restricting analyses to species represented in the phylogenetic dataset.

| tier | model | term | estimate__glm_full | estimate__glm_matched | conf. low__glm_full | conf. high__glm_full | conf. low__glm_matched | conf. high__glm_matched | OR__glm_full | OR__glm_matched | p__glm_full | p__glm_matched | delta_beta_subset | delta_OR_subset |
|---|---|---|---|---|---|---|---|---|---|---|---|---|---|---|
| ge1 | m_clim | BIO10_med | -0.349 | -0.198 | -0.593 | -0.106 | -0.458 | 0.062 | 0.705 | 0.820 | 0.005 | 0.136 | 0.151 | 1.163 |
| ge1 | m_clim | BIO6_med | 0.866 | 0.894 | 0.684 | 1.049 | 0.703 | 1.084 | 2.378 | 2.444 | 0.000 | 0.000 | 0.028 | 1.028 |
| ge1 | m_clim | MCWD_med | -0.491 | -0.343 | -0.651 | -0.330 | -0.519 | -0.168 | 0.612 | 0.709 | 0.000 | 0.000 | 0.147 | 1.159 |
| ge1 | m_clim | elev_med | -0.056 | 0.063 | -0.251 | 0.139 | -0.141 | 0.267 | 0.945 | 1.065 | 0.572 | 0.544 | 0.119 | 1.127 |
| ge1 | m_clim_breadth | BIO10_mad | -0.294 | -0.286 | -0.545 | -0.042 | -0.542 | -0.029 | 0.746 | 0.751 | 0.022 | 0.029 | 0.008 | 1.008 |
| ge1 | m_clim_breadth | BIO10_med | -0.365 | -0.238 | -0.635 | -0.095 | -0.525 | 0.049 | 0.694 | 0.788 | 0.008 | 0.105 | 0.128 | 1.136 |
| ge1 | m_clim_breadth | BIO6_mad | 0.125 | 0.145 | -0.112 | 0.363 | -0.094 | 0.384 | 1.134 | 1.156 | 0.301 | 0.236 | 0.019 | 1.019 |
| ge1 | m_clim_breadth | BIO6_med | 0.906 | 0.928 | 0.690 | 1.123 | 0.707 | 1.150 | 2.475 | 2.530 | 0.000 | 0.000 | 0.022 | 1.022 |
| ge1 | m_clim_breadth | MCWD_mad | -0.160 | -0.187 | -0.319 | -0.002 | -0.355 | -0.019 | 0.852 | 0.830 | 0.048 | 0.030 | -0.027 | 0.974 |
| ge1 | m_clim_breadth | MCWD_med | -0.568 | -0.444 | -0.757 | -0.380 | -0.650 | -0.238 | 0.566 | 0.642 | 0.000 | 0.000 | 0.125 | 1.133 |
| ge1 | m_clim_breadth | elev_mad | 0.279 | 0.268 | 0.058 | 0.500 | 0.042 | 0.494 | 1.322 | 1.308 | 0.014 | 0.020 | -0.011 | 0.989 |
| ge1 | m_clim_breadth | elev_med | -0.069 | 0.022 | -0.295 | 0.157 | -0.213 | 0.257 | 0.934 | 1.022 | 0.551 | 0.854 | 0.091 | 1.095 |
| ge1 | m_clim_island | BIO10_med | -0.227 | -0.058 | -0.480 | 0.026 | -0.328 | 0.212 | 0.797 | 0.944 | 0.078 | 0.675 | 0.170 | 1.185 |
| ge1 | m_clim_island | BIO6_med | 0.679 | 0.704 | 0.489 | 0.870 | 0.505 | 0.904 | 1.972 | 2.023 | 0.000 | 0.000 | 0.025 | 1.025 |
| ge1 | m_clim_island | MCWD_med | -0.526 | -0.367 | -0.694 | -0.357 | -0.551 | -0.183 | 0.591 | 0.693 | 0.000 | 0.000 | 0.159 | 1.172 |
| ge1 | m_clim_island | elev_med | 0.041 | 0.159 | -0.164 | 0.245 | -0.054 | 0.372 | 1.041 | 1.173 | 0.697 | 0.143 | 0.119 | 1.126 |
| ge1 | m_clim_island | is_island | 1.902 | 1.844 | 1.475 | 2.329 | 1.386 | 2.301 | 6.701 | 6.319 | 0.000 | 0.000 | -0.059 | 0.943 |
| ge1 | m_clim_mainland | BIO10_med | -0.133 | 0.015 | -0.399 | 0.132 | -0.268 | 0.298 | 0.875 | 1.015 | 0.325 | 0.918 | 0.148 | 1.160 |
| ge1 | m_clim_mainland | BIO6_med | 0.639 | 0.672 | 0.448 | 0.831 | 0.473 | 0.872 | 1.895 | 1.959 | 0.000 | 0.000 | 0.033 | 1.033 |
| ge1 | m_clim_mainland | MCWD_med | -0.454 | -0.303 | -0.629 | -0.280 | -0.493 | -0.114 | 0.635 | 0.738 | 0.000 | 0.002 | 0.151 | 1.163 |
| ge1 | m_clim_mainland | elev_med | 0.069 | 0.177 | -0.143 | 0.282 | -0.044 | 0.398 | 1.072 | 1.193 | 0.522 | 0.117 | 0.107 | 1.113 |
| ge5 | m_clim | BIO10_med | -0.353 | -0.210 | -0.642 | -0.064 | -0.517 | 0.097 | 0.703 | 0.810 | 0.017 | 0.180 | 0.143 | 1.154 |
| ge5 | m_clim | BIO6_med | 0.933 | 0.939 | 0.702 | 1.163 | 0.702 | 1.176 | 2.541 | 2.558 | 0.000 | 0.000 | 0.007 | 1.007 |
| ge5 | m_clim | MCWD_med | -0.500 | -0.350 | -0.697 | -0.303 | -0.563 | -0.138 | 0.607 | 0.705 | 0.000 | 0.001 | 0.150 | 1.162 |
| ge5 | m_clim | elev_med | 0.004 | 0.086 | -0.216 | 0.224 | -0.144 | 0.315 | 1.004 | 1.090 | 0.970 | 0.464 | 0.082 | 1.085 |
| ge5 | m_clim_breadth | BIO10_mad | -0.412 | -0.401 | -0.683 | -0.140 | -0.674 | -0.129 | 0.662 | 0.669 | 0.003 | 0.004 | 0.011 | 1.011 |
| ge5 | m_clim_breadth | BIO10_med | -0.314 | -0.176 | -0.607 | -0.020 | -0.487 | 0.135 | 0.731 | 0.838 | 0.036 | 0.267 | 0.137 | 1.147 |
| ge5 | m_clim_breadth | BIO6_mad | 0.120 | 0.139 | -0.135 | 0.374 | -0.114 | 0.393 | 1.127 | 1.150 | 0.358 | 0.281 | 0.020 | 1.020 |
| ge5 | m_clim_breadth | BIO6_med | 0.941 | 0.951 | 0.705 | 1.177 | 0.709 | 1.193 | 2.564 | 2.588 | 0.000 | 0.000 | 0.010 | 1.010 |
| ge5 | m_clim_breadth | MCWD_mad | -0.147 | -0.167 | -0.316 | 0.021 | -0.345 | 0.010 | 0.863 | 0.846 | 0.087 | 0.064 | -0.020 | 0.980 |
| ge5 | m_clim_breadth | MCWD_med | -0.556 | -0.414 | -0.760 | -0.353 | -0.634 | -0.194 | 0.573 | 0.661 | 0.000 | 0.000 | 0.143 | 1.153 |
| ge5 | m_clim_breadth | elev_mad | 0.347 | 0.336 | 0.110 | 0.583 | 0.097 | 0.574 | 1.415 | 1.399 | 0.004 | 0.006 | -0.011 | 0.989 |
| ge5 | m_clim_breadth | elev_med | 0.011 | 0.085 | -0.234 | 0.256 | -0.168 | 0.337 | 1.011 | 1.088 | 0.930 | 0.512 | 0.074 | 1.076 |
| ge5 | m_clim_island | BIO10_med | -0.209 | -0.075 | -0.510 | 0.092 | -0.393 | 0.242 | 0.811 | 0.927 | 0.173 | 0.642 | 0.134 | 1.143 |
| ge5 | m_clim_island | BIO6_med | 0.739 | 0.747 | 0.500 | 0.978 | 0.500 | 0.993 | 2.093 | 2.110 | 0.000 | 0.000 | 0.008 | 1.008 |
| ge5 | m_clim_island | MCWD_med | -0.521 | -0.376 | -0.728 | -0.315 | -0.598 | -0.154 | 0.594 | 0.687 | 0.000 | 0.001 | 0.146 | 1.157 |
| ge5 | m_clim_island | elev_med | 0.090 | 0.160 | -0.142 | 0.322 | -0.079 | 0.400 | 1.094 | 1.174 | 0.447 | 0.189 | 0.071 | 1.073 |
| ge5 | m_clim_island | is_island | 1.954 | 1.806 | 1.453 | 2.454 | 1.286 | 2.327 | 7.054 | 6.087 | 0.000 | 0.000 | -0.147 | 0.863 |
| ge5 | m_clim_mainland | BIO10_med | -0.146 | -0.007 | -0.462 | 0.171 | -0.341 | 0.327 | 0.865 | 0.993 | 0.367 | 0.967 | 0.139 | 1.149 |
| ge5 | m_clim_mainland | BIO6_med | 0.687 | 0.706 | 0.447 | 0.928 | 0.457 | 0.954 | 1.988 | 2.025 | 0.000 | 0.000 | 0.018 | 1.018 |
| ge5 | m_clim_mainland | MCWD_med | -0.484 | -0.330 | -0.697 | -0.271 | -0.560 | -0.101 | 0.616 | 0.719 | 0.000 | 0.005 | 0.154 | 1.166 |
| ge5 | m_clim_mainland | elev_med | 0.111 | 0.182 | -0.130 | 0.351 | -0.066 | 0.431 | 1.117 | 1.200 | 0.368 | 0.151 | 0.072 | 1.075 |
| ge10 | m_clim | BIO10_med | -0.265 | -0.148 | -0.596 | 0.067 | -0.495 | 0.199 | 0.767 | 0.863 | 0.118 | 0.404 | 0.117 | 1.124 |
| ge10 | m_clim | BIO6_med | 0.901 | 0.897 | 0.634 | 1.168 | 0.620 | 1.174 | 2.462 | 2.453 | 0.000 | 0.000 | -0.004 | 0.996 |
| ge10 | m_clim | MCWD_med | -0.438 | -0.303 | -0.658 | -0.217 | -0.539 | -0.067 | 0.646 | 0.739 | 0.000 | 0.012 | 0.135 | 1.144 |
| ge10 | m_clim | elev_med | 0.018 | 0.083 | -0.231 | 0.267 | -0.175 | 0.341 | 1.018 | 1.087 | 0.887 | 0.528 | 0.065 | 1.067 |
| ge10 | m_clim_breadth | BIO10_mad | -0.446 | -0.425 | -0.753 | -0.139 | -0.730 | -0.120 | 0.640 | 0.654 | 0.004 | 0.006 | 0.021 | 1.021 |
| ge10 | m_clim_breadth | BIO10_med | -0.219 | -0.110 | -0.557 | 0.119 | -0.463 | 0.242 | 0.803 | 0.895 | 0.205 | 0.539 | 0.108 | 1.114 |
| ge10 | m_clim_breadth | BIO6_mad | 0.141 | 0.154 | -0.143 | 0.425 | -0.127 | 0.435 | 1.151 | 1.166 | 0.331 | 0.282 | 0.013 | 1.013 |
| ge10 | m_clim_breadth | BIO6_med | 0.917 | 0.913 | 0.639 | 1.196 | 0.627 | 1.200 | 2.502 | 2.492 | 0.000 | 0.000 | -0.004 | 0.996 |
| ge10 | m_clim_breadth | MCWD_mad | -0.168 | -0.184 | -0.365 | 0.030 | -0.386 | 0.018 | 0.846 | 0.832 | 0.097 | 0.073 | -0.016 | 0.984 |
| ge10 | m_clim_breadth | MCWD_med | -0.501 | -0.370 | -0.729 | -0.272 | -0.615 | -0.125 | 0.606 | 0.691 | 0.000 | 0.003 | 0.131 | 1.139 |
| ge10 | m_clim_breadth | elev_mad | 0.336 | 0.302 | 0.067 | 0.605 | 0.032 | 0.572 | 1.399 | 1.353 | 0.014 | 0.028 | -0.034 | 0.967 |
| ge10 | m_clim_breadth | elev_med | 0.046 | 0.107 | -0.232 | 0.325 | -0.180 | 0.393 | 1.047 | 1.113 | 0.745 | 0.466 | 0.061 | 1.062 |
| ge10 | m_clim_island | BIO10_med | -0.128 | -0.026 | -0.473 | 0.217 | -0.385 | 0.332 | 0.880 | 0.974 | 0.466 | 0.886 | 0.102 | 1.107 |
| ge10 | m_clim_island | BIO6_med | 0.699 | 0.700 | 0.422 | 0.976 | 0.412 | 0.988 | 2.012 | 2.013 | 0.000 | 0.000 | 0.001 | 1.001 |
| ge10 | m_clim_island | MCWD_med | -0.474 | -0.344 | -0.706 | -0.242 | -0.591 | -0.098 | 0.623 | 0.709 | 0.000 | 0.006 | 0.130 | 1.138 |
| ge10 | m_clim_island | elev_med | 0.093 | 0.139 | -0.169 | 0.354 | -0.129 | 0.408 | 1.097 | 1.150 | 0.488 | 0.309 | 0.047 | 1.048 |
| ge10 | m_clim_island | is_island | 2.145 | 1.957 | 1.557 | 2.733 | 1.352 | 2.562 | 8.540 | 7.078 | 0.000 | 0.000 | -0.188 | 0.829 |
| ge10 | m_clim_mainland | BIO10_med | -0.076 | 0.032 | -0.437 | 0.285 | -0.344 | 0.409 | 0.927 | 1.033 | 0.680 | 0.866 | 0.108 | 1.114 |
| ge10 | m_clim_mainland | BIO6_med | 0.656 | 0.663 | 0.376 | 0.936 | 0.372 | 0.953 | 1.927 | 1.940 | 0.000 | 0.000 | 0.007 | 1.007 |
| ge10 | m_clim_mainland | MCWD_med | -0.436 | -0.300 | -0.674 | -0.198 | -0.554 | -0.046 | 0.647 | 0.741 | 0.000 | 0.020 | 0.136 | 1.145 |
| ge10 | m_clim_mainland | elev_med | 0.098 | 0.149 | -0.173 | 0.368 | -0.129 | 0.427 | 1.103 | 1.160 | 0.478 | 0.295 | 0.051 | 1.052 |

#### Table S12. Effect of phylogenetic correction on model coefficients

Comparison of standardized regression coefficients between non-phylogenetic (GLM matched) and phylogenetic (phyloGLM matched) models fitted to identical datasets. For each predictor, the table reports coefficients, confidence intervals, odds ratios, and P-values, along with differences (Δβ and ΔOR), quantifying the impact of accounting for phylogenetic relatedness.

| tier | model | term | estimate__glm_matched | estimate__phyloglm_matched | conf. low__glm_matched | conf. high__glm_matched | conf. low__phyloglm_matched | conf. high__phyloglm_matched | OR__glm_matched | OR__phyloglm_matched | p__glm_matched | p__phyloglm_matched | delta_beta_phylogeny | delta_OR_phylogeny |
|---|---|---|---|---|---|---|---|---|---|---|---|---|---|---|
| ge1 | m_clim | BIO10_med | -0.198 | -0.348 | -0.458 | 0.062 | -0.590 | -0.106 | 0.820 | 0.706 | 0.136 | 0.005 | -0.150 | 0.861 |
| ge1 | m_clim | BIO6_med | 0.894 | 0.620 | 0.703 | 1.084 | 0.415 | 0.825 | 2.444 | 1.859 | 0.000 | 0.000 | -0.273 | 0.761 |
| ge1 | m_clim | MCWD_med | -0.343 | -0.179 | -0.519 | -0.168 | -0.343 | -0.016 | 0.709 | 0.836 | 0.000 | 0.032 | 0.164 | 1.178 |
| ge1 | m_clim | elev_med | 0.063 | 0.006 | -0.141 | 0.267 | -0.141 | 0.152 | 1.065 | 1.006 | 0.544 | 0.940 | -0.057 | 0.944 |
| ge1 | m_clim_breadth | BIO10_mad | -0.286 | -0.074 | -0.542 | -0.029 | -0.215 | 0.067 | 0.751 | 0.928 | 0.029 | 0.302 | 0.212 | 1.236 |
| ge1 | m_clim_breadth | BIO10_med | -0.238 | -0.389 | -0.525 | 0.049 | -0.640 | -0.138 | 0.788 | 0.678 | 0.105 | 0.002 | -0.151 | 0.859 |
| ge1 | m_clim_breadth | BIO6_mad | 0.145 | 0.132 | -0.094 | 0.384 | 0.004 | 0.261 | 1.156 | 1.141 | 0.236 | 0.043 | -0.012 | 0.988 |
| ge1 | m_clim_breadth | BIO6_med | 0.928 | 0.515 | 0.707 | 1.150 | 0.309 | 0.721 | 2.530 | 1.674 | 0.000 | 0.000 | -0.413 | 0.661 |
| ge1 | m_clim_breadth | MCWD_mad | -0.187 | -0.028 | -0.355 | -0.019 | -0.134 | 0.079 | 0.830 | 0.973 | 0.030 | 0.610 | 0.159 | 1.172 |
| ge1 | m_clim_breadth | MCWD_med | -0.444 | -0.287 | -0.650 | -0.238 | -0.477 | -0.096 | 0.642 | 0.751 | 0.000 | 0.003 | 0.157 | 1.170 |
| ge1 | m_clim_breadth | elev_mad | 0.268 | 0.035 | 0.042 | 0.494 | -0.089 | 0.159 | 1.308 | 1.035 | 0.020 | 0.581 | -0.233 | 0.792 |
| ge1 | m_clim_breadth | elev_med | 0.022 | 0.011 | -0.213 | 0.257 | -0.136 | 0.157 | 1.022 | 1.011 | 0.854 | 0.884 | -0.011 | 0.989 |
| ge1 | m_clim_island | BIO10_med | -0.058 | -0.309 | -0.328 | 0.212 | -0.542 | -0.075 | 0.944 | 0.734 | 0.675 | 0.010 | -0.251 | 0.778 |
| ge1 | m_clim_island | BIO6_med | 0.704 | 0.605 | 0.505 | 0.904 | 0.408 | 0.803 | 2.023 | 1.832 | 0.000 | 0.000 | -0.099 | 0.906 |
| ge1 | m_clim_island | MCWD_med | -0.367 | -0.174 | -0.551 | -0.183 | -0.333 | -0.015 | 0.693 | 0.840 | 0.000 | 0.032 | 0.192 | 1.212 |
| ge1 | m_clim_island | elev_med | 0.159 | 0.090 | -0.054 | 0.372 | -0.057 | 0.237 | 1.173 | 1.095 | 0.143 | 0.228 | -0.069 | 0.933 |
| ge1 | m_clim_island | is_island | 1.844 | 0.986 | 1.386 | 2.301 | 0.461 | 1.511 | 6.319 | 2.681 | 0.000 | 0.000 | -0.858 | 0.424 |
| ge1 | m_clim_mainland | BIO10_med | 0.015 | -0.329 | -0.268 | 0.298 | -0.586 | -0.072 | 1.015 | 0.720 | 0.918 | 0.012 | -0.344 | 0.709 |
| ge1 | m_clim_mainland | BIO6_med | 0.672 | 0.548 | 0.473 | 0.872 | 0.337 | 0.760 | 1.959 | 1.730 | 0.000 | 0.000 | -0.124 | 0.883 |
| ge1 | m_clim_mainland | MCWD_med | -0.303 | -0.245 | -0.493 | -0.114 | -0.424 | -0.065 | 0.738 | 0.783 | 0.002 | 0.007 | 0.059 | 1.060 |
| ge1 | m_clim_mainland | elev_med | 0.177 | 0.079 | -0.044 | 0.398 | -0.074 | 0.232 | 1.193 | 1.082 | 0.117 | 0.311 | -0.097 | 0.907 |
| ge5 | m_clim | BIO10_med | -0.210 | -0.325 | -0.517 | 0.097 | -0.589 | -0.060 | 0.810 | 0.723 | 0.180 | 0.016 | -0.115 | 0.892 |
| ge5 | m_clim | BIO6_med | 0.939 | 0.530 | 0.702 | 1.176 | 0.316 | 0.745 | 2.558 | 1.699 | 0.000 | 0.000 | -0.409 | 0.664 |
| ge5 | m_clim | MCWD_med | -0.350 | -0.254 | -0.563 | -0.138 | -0.451 | -0.056 | 0.705 | 0.776 | 0.001 | 0.012 | 0.096 | 1.101 |
| ge5 | m_clim | elev_med | 0.086 | 0.080 | -0.144 | 0.315 | -0.075 | 0.235 | 1.090 | 1.083 | 0.464 | 0.313 | -0.006 | 0.994 |
| ge5 | m_clim_breadth | BIO10_mad | -0.401 | -0.123 | -0.674 | -0.129 | -0.278 | 0.033 | 0.669 | 0.884 | 0.004 | 0.121 | 0.279 | 1.321 |
| ge5 | m_clim_breadth | BIO10_med | -0.176 | -0.391 | -0.487 | 0.135 | -0.662 | -0.120 | 0.838 | 0.676 | 0.267 | 0.005 | -0.215 | 0.807 |
| ge5 | m_clim_breadth | BIO6_mad | 0.139 | 0.134 | -0.114 | 0.393 | -0.009 | 0.278 | 1.150 | 1.144 | 0.281 | 0.066 | -0.005 | 0.995 |
| ge5 | m_clim_breadth | BIO6_med | 0.951 | 0.579 | 0.709 | 1.193 | 0.351 | 0.806 | 2.588 | 1.784 | 0.000 | 0.000 | -0.372 | 0.689 |
| ge5 | m_clim_breadth | MCWD_mad | -0.167 | -0.008 | -0.345 | 0.010 | -0.123 | 0.106 | 0.846 | 0.992 | 0.064 | 0.886 | 0.159 | 1.172 |
| ge5 | m_clim_breadth | MCWD_med | -0.414 | -0.290 | -0.634 | -0.194 | -0.496 | -0.084 | 0.661 | 0.748 | 0.000 | 0.006 | 0.124 | 1.132 |
| ge5 | m_clim_breadth | elev_mad | 0.336 | 0.060 | 0.097 | 0.574 | -0.075 | 0.196 | 1.399 | 1.062 | 0.006 | 0.382 | -0.275 | 0.759 |
| ge5 | m_clim_breadth | elev_med | 0.085 | 0.032 | -0.168 | 0.337 | -0.131 | 0.195 | 1.088 | 1.033 | 0.512 | 0.699 | -0.052 | 0.949 |
| ge5 | m_clim_island | BIO10_med | -0.075 | -0.226 | -0.393 | 0.242 | -0.480 | 0.029 | 0.927 | 0.798 | 0.642 | 0.082 | -0.150 | 0.860 |
| ge5 | m_clim_island | BIO6_med | 0.747 | 0.516 | 0.500 | 0.993 | 0.305 | 0.727 | 2.110 | 1.676 | 0.000 | 0.000 | -0.230 | 0.794 |
| ge5 | m_clim_island | MCWD_med | -0.376 | -0.179 | -0.598 | -0.154 | -0.366 | 0.008 | 0.687 | 0.836 | 0.001 | 0.061 | 0.197 | 1.218 |
| ge5 | m_clim_island | elev_med | 0.160 | 0.126 | -0.079 | 0.400 | -0.033 | 0.285 | 1.174 | 1.134 | 0.189 | 0.122 | -0.035 | 0.966 |
| ge5 | m_clim_island | is_island | 1.806 | 1.043 | 1.286 | 2.327 | 0.448 | 1.639 | 6.087 | 2.838 | 0.000 | 0.001 | -0.763 | 0.466 |
| ge5 | m_clim_mainland | BIO10_med | -0.007 | -0.266 | -0.341 | 0.327 | -0.536 | 0.004 | 0.993 | 0.766 | 0.967 | 0.054 | -0.259 | 0.772 |
| ge5 | m_clim_mainland | BIO6_med | 0.706 | 0.462 | 0.457 | 0.954 | 0.247 | 0.678 | 2.025 | 1.588 | 0.000 | 0.000 | -0.243 | 0.784 |
| ge5 | m_clim_mainland | MCWD_med | -0.330 | -0.246 | -0.560 | -0.101 | -0.450 | -0.041 | 0.719 | 0.782 | 0.005 | 0.019 | 0.085 | 1.089 |
| ge5 | m_clim_mainland | elev_med | 0.182 | 0.075 | -0.066 | 0.431 | -0.082 | 0.233 | 1.200 | 1.078 | 0.151 | 0.349 | -0.107 | 0.898 |
| ge10 | m_clim | BIO10_med | -0.148 | -0.297 | -0.495 | 0.199 | -0.578 | -0.016 | 0.863 | 0.743 | 0.404 | 0.038 | -0.149 | 0.862 |
| ge10 | m_clim | BIO6_med | 0.897 | 0.504 | 0.620 | 1.174 | 0.277 | 0.730 | 2.453 | 1.655 | 0.000 | 0.000 | -0.394 | 0.675 |
| ge10 | m_clim | MCWD_med | -0.303 | -0.297 | -0.539 | -0.067 | -0.518 | -0.075 | 0.739 | 0.743 | 0.012 | 0.009 | 0.006 | 1.006 |
| ge10 | m_clim | elev_med | 0.083 | 0.102 | -0.175 | 0.341 | -0.063 | 0.267 | 1.087 | 1.107 | 0.528 | 0.228 | 0.019 | 1.019 |
| ge10 | m_clim_breadth | BIO10_mad | -0.425 | -0.119 | -0.730 | -0.120 | -0.274 | 0.035 | 0.654 | 0.888 | 0.006 | 0.131 | 0.306 | 1.358 |
| ge10 | m_clim_breadth | BIO10_med | -0.110 | -0.319 | -0.463 | 0.242 | -0.592 | -0.045 | 0.895 | 0.727 | 0.539 | 0.023 | -0.208 | 0.812 |
| ge10 | m_clim_breadth | BIO6_mad | 0.154 | 0.154 | -0.127 | 0.435 | 0.011 | 0.297 | 1.166 | 1.167 | 0.282 | 0.034 | 0.000 | 1.000 |
| ge10 | m_clim_breadth | BIO6_med | 0.913 | 0.514 | 0.627 | 1.200 | 0.282 | 0.746 | 2.492 | 1.672 | 0.000 | 0.000 | -0.399 | 0.671 |
| ge10 | m_clim_breadth | MCWD_mad | -0.184 | -0.040 | -0.386 | 0.018 | -0.163 | 0.083 | 0.832 | 0.961 | 0.073 | 0.521 | 0.144 | 1.155 |
| ge10 | m_clim_breadth | MCWD_med | -0.370 | -0.286 | -0.615 | -0.125 | -0.504 | -0.068 | 0.691 | 0.751 | 0.003 | 0.010 | 0.084 | 1.087 |
| ge10 | m_clim_breadth | elev_mad | 0.302 | 0.056 | 0.032 | 0.572 | -0.081 | 0.192 | 1.353 | 1.057 | 0.028 | 0.422 | -0.246 | 0.782 |
| ge10 | m_clim_breadth | elev_med | 0.107 | 0.041 | -0.180 | 0.393 | -0.126 | 0.208 | 1.113 | 1.042 | 0.466 | 0.627 | -0.065 | 0.937 |
| ge10 | m_clim_island | BIO10_med | -0.026 | -0.199 | -0.385 | 0.332 | -0.463 | 0.066 | 0.974 | 0.820 | 0.886 | 0.141 | -0.172 | 0.842 |
| ge10 | m_clim_island | BIO6_med | 0.700 | 0.461 | 0.412 | 0.988 | 0.246 | 0.676 | 2.013 | 1.585 | 0.000 | 0.000 | -0.239 | 0.788 |
| ge10 | m_clim_island | MCWD_med | -0.344 | -0.211 | -0.591 | -0.098 | -0.414 | -0.009 | 0.709 | 0.810 | 0.006 | 0.041 | 0.133 | 1.143 |
| ge10 | m_clim_island | elev_med | 0.139 | 0.117 | -0.129 | 0.408 | -0.046 | 0.281 | 1.150 | 1.125 | 0.309 | 0.160 | -0.022 | 0.978 |
| ge10 | m_clim_island | is_island | 1.957 | 1.255 | 1.352 | 2.562 | 0.569 | 1.941 | 7.078 | 3.507 | 0.000 | 0.000 | -0.702 | 0.495 |
| ge10 | m_clim_mainland | BIO10_med | 0.032 | -0.227 | -0.344 | 0.409 | -0.508 | 0.054 | 1.033 | 0.797 | 0.866 | 0.114 | -0.259 | 0.772 |
| ge10 | m_clim_mainland | BIO6_med | 0.663 | 0.374 | 0.372 | 0.953 | 0.160 | 0.589 | 1.940 | 1.454 | 0.000 | 0.001 | -0.288 | 0.750 |
| ge10 | m_clim_mainland | MCWD_med | -0.300 | -0.287 | -0.554 | -0.046 | -0.517 | -0.057 | 0.741 | 0.751 | 0.020 | 0.014 | 0.013 | 1.013 |
| ge10 | m_clim_mainland | elev_med | 0.149 | 0.065 | -0.129 | 0.427 | -0.098 | 0.229 | 1.160 | 1.068 | 0.295 | 0.434 | -0.083 | 0.920 |

#### Table S13. Model comparison based on AIC across frameworks and tiers

AIC values and ΔAIC (relative to the best model within each framework–tier combination) for all model formulations across ge tiers and statistical frameworks. This table provides the numerical basis for model selection and comparison of support among competing models.

| tier | model | framework | n | formula | logLik | AIC | alpha | deltaAIC_within_framework |
|---|---|---|---|---|---|---|---|---|
| ge1 | m_clim_breadth | glm_full | 2,896 | woody_bin ~ MCWD_med + BIO10_med + BIO6_med + elev_med + MCWD_mad + BIO10_mad + BIO6_mad + elev_mad | -791.25 | 1,600.49 | NA | 0.00 |
| ge1 | m_clim_mainland | glm_full | 3,299 | woody_bin ~ MCWD_med + BIO10_med + BIO6_med + elev_med | -864.01 | 1,738.03 | NA | 137.53 |
| ge1 | m_clim_island | glm_full | 3,412 | woody_bin ~ MCWD_med + BIO10_med + BIO6_med + elev_med + is_island | -928.14 | 1,868.27 | NA | 267.78 |
| ge1 | m_clim | glm_full | 3,412 | woody_bin ~ MCWD_med + BIO10_med + BIO6_med + elev_med | -963.85 | 1,937.71 | NA | 337.22 |
| ge1 | m_clim_breadth | glm_matched | 2,217 | woody_bin ~ MCWD_med + BIO10_med + BIO6_med + elev_med + MCWD_mad + BIO10_mad + BIO6_mad + elev_mad | -722.21 | 1,462.42 | NA | 0.00 |
| ge1 | m_clim_mainland | glm_matched | 2,429 | woody_bin ~ MCWD_med + BIO10_med + BIO6_med + elev_med | -774.13 | 1,558.25 | NA | 95.83 |
| ge1 | m_clim_island | glm_matched | 2,527 | woody_bin ~ MCWD_med + BIO10_med + BIO6_med + elev_med + is_island | -831.09 | 1,674.18 | NA | 211.76 |
| ge1 | m_clim | glm_matched | 2,527 | woody_bin ~ MCWD_med + BIO10_med + BIO6_med + elev_med | -861.76 | 1,733.52 | NA | 271.10 |
| ge1 | m_clim_breadth | phyloglm_matched | 2,217 | woody_bin ~ MCWD_med + BIO10_med + BIO6_med + elev_med + MCWD_mad + BIO10_mad + BIO6_mad + elev_mad | -522.66 | 1,065.32 | 0.061 | 0.00 |
| ge1 | m_clim_mainland | phyloglm_matched | 2,429 | woody_bin ~ MCWD_med + BIO10_med + BIO6_med + elev_med | -547.52 | 1,107.04 | 0.056 | 41.72 |
| ge1 | m_clim_island | phyloglm_matched | 2,527 | woody_bin ~ MCWD_med + BIO10_med + BIO6_med + elev_med + is_island | -606.35 | 1,226.69 | 0.059 | 161.38 |
| ge1 | m_clim | phyloglm_matched | 2,527 | woody_bin ~ MCWD_med + BIO10_med + BIO6_med + elev_med | -612.38 | 1,236.75 | 0.058 | 171.43 |
| ge5 | m_clim_mainland | glm_full | 2,520 | woody_bin ~ MCWD_med + BIO10_med + BIO6_med + elev_med | -639.08 | 1,288.16 | NA | 0.00 |
| ge5 | m_clim_island | glm_full | 2,601 | woody_bin ~ MCWD_med + BIO10_med + BIO6_med + elev_med + is_island | -685.94 | 1,383.88 | NA | 95.72 |
| ge5 | m_clim_breadth | glm_full | 2,601 | woody_bin ~ MCWD_med + BIO10_med + BIO6_med + elev_med + MCWD_mad + BIO10_mad + BIO6_mad + elev_mad | -703.19 | 1,424.38 | NA | 136.22 |
| ge5 | m_clim | glm_full | 2,601 | woody_bin ~ MCWD_med + BIO10_med + BIO6_med + elev_med | -713.12 | 1,436.24 | NA | 148.08 |
| ge5 | m_clim_mainland | glm_matched | 1,944 | woody_bin ~ MCWD_med + BIO10_med + BIO6_med + elev_med | -590.55 | 1,191.09 | NA | 0.00 |
| ge5 | m_clim_island | glm_matched | 2,018 | woody_bin ~ MCWD_med + BIO10_med + BIO6_med + elev_med + is_island | -634.57 | 1,281.14 | NA | 90.05 |
| ge5 | m_clim_breadth | glm_matched | 2,018 | woody_bin ~ MCWD_med + BIO10_med + BIO6_med + elev_med + MCWD_mad + BIO10_mad + BIO6_mad + elev_mad | -646.79 | 1,311.58 | NA | 120.49 |
| ge5 | m_clim | glm_matched | 2,018 | woody_bin ~ MCWD_med + BIO10_med + BIO6_med + elev_med | -656.76 | 1,323.51 | NA | 132.42 |
| ge5 | m_clim_mainland | phyloglm_matched | 1,944 | woody_bin ~ MCWD_med + BIO10_med + BIO6_med + elev_med | -429.91 | 871.83 | 0.055 | 0.00 |
| ge5 | m_clim_island | phyloglm_matched | 2,018 | woody_bin ~ MCWD_med + BIO10_med + BIO6_med + elev_med + is_island | -476.76 | 967.52 | 0.064 | 95.69 |
| ge5 | m_clim | phyloglm_matched | 2,018 | woody_bin ~ MCWD_med + BIO10_med + BIO6_med + elev_med | -482.27 | 976.54 | 0.060 | 104.72 |
| ge5 | m_clim_breadth | phyloglm_matched | 2,018 | woody_bin ~ MCWD_med + BIO10_med + BIO6_med + elev_med + MCWD_mad + BIO10_mad + BIO6_mad + elev_mad | -481.02 | 982.03 | 0.063 | 110.21 |
| ge10 | m_clim_mainland | glm_full | 2,101 | woody_bin ~ MCWD_med + BIO10_med + BIO6_med + elev_med | -525.11 | 1,060.22 | NA | 0.00 |
| ge10 | m_clim_island | glm_full | 2,158 | woody_bin ~ MCWD_med + BIO10_med + BIO6_med + elev_med + is_island | -558.41 | 1,128.82 | NA | 68.60 |
| ge10 | m_clim_breadth | glm_full | 2,158 | woody_bin ~ MCWD_med + BIO10_med + BIO6_med + elev_med + MCWD_mad + BIO10_mad + BIO6_mad + elev_mad | -572.73 | 1,163.46 | NA | 103.24 |
| ge10 | m_clim | glm_full | 2,158 | woody_bin ~ MCWD_med + BIO10_med + BIO6_med + elev_med | -582.12 | 1,174.24 | NA | 114.02 |
| ge10 | m_clim_mainland | glm_matched | 1,646 | woody_bin ~ MCWD_med + BIO10_med + BIO6_med + elev_med | -487.92 | 985.84 | NA | 0.00 |
| ge10 | m_clim_island | glm_matched | 1,699 | woody_bin ~ MCWD_med + BIO10_med + BIO6_med + elev_med + is_island | -519.73 | 1,051.45 | NA | 65.61 |
| ge10 | m_clim_breadth | glm_matched | 1,699 | woody_bin ~ MCWD_med + BIO10_med + BIO6_med + elev_med + MCWD_mad + BIO10_mad + BIO6_mad + elev_mad | -529.82 | 1,077.64 | NA | 91.80 |
| ge10 | m_clim | glm_matched | 1,699 | woody_bin ~ MCWD_med + BIO10_med + BIO6_med + elev_med | -538.80 | 1,087.61 | NA | 101.77 |
| ge10 | m_clim_mainland | phyloglm_matched | 1,646 | woody_bin ~ MCWD_med + BIO10_med + BIO6_med + elev_med | -365.03 | 742.05 | 0.053 | 0.00 |
| ge10 | m_clim_island | phyloglm_matched | 1,699 | woody_bin ~ MCWD_med + BIO10_med + BIO6_med + elev_med + is_island | -399.20 | 812.40 | 0.060 | 70.34 |
| ge10 | m_clim | phyloglm_matched | 1,699 | woody_bin ~ MCWD_med + BIO10_med + BIO6_med + elev_med | -406.14 | 824.29 | 0.057 | 82.23 |
| ge10 | m_clim_breadth | phyloglm_matched | 1,699 | woody_bin ~ MCWD_med + BIO10_med + BIO6_med + elev_med + MCWD_mad + BIO10_mad + BIO6_mad + elev_mad | -403.91 | 827.81 | 0.061 | 85.76 |

**Table S14. Likelihood-based model performance and phylogenetic signal for matched datasets**

Model performance metrics for matched datasets, including log-likelihood of full and null models, McFadden's pseudo-$R^2$, and (for phylogenetic models) the estimated phylogenetic signal parameter (α). Values are reported for each ge tier and model formulation, allowing comparison of explanatory power and the contribution of phylogenetic structure.

| tier | model | framework | n | logLik_full | logLik_null | R2 | alpha |
|---|---|---|---|---|---|---|---|
| ge1 | m_clim | glm_matched | 2,527 | -861.759 | -967.79 | 0.11 | NA |
| ge1 | m_clim | phyloglm_matched | 2,527 | -612.376 | -665.56 | 0.08 | 0.058 |
| ge1 | m_clim_breadth | glm_matched | 2,217 | -722.210 | -821.33 | 0.12 | NA |
| ge1 | m_clim_breadth | phyloglm_matched | 2,217 | -522.659 | -561.87 | 0.07 | 0.061 |
| ge1 | m_clim_island | glm_matched | 2,527 | -831.088 | -967.79 | 0.14 | NA |
| ge1 | m_clim_island | phyloglm_matched | 2,527 | -606.347 | -665.56 | 0.09 | 0.059 |
| ge1 | m_clim_mainland | glm_matched | 2,429 | -774.125 | -845.46 | 0.08 | NA |
| ge1 | m_clim_mainland | phyloglm_matched | 2,429 | -547.520 | -601.82 | 0.09 | 0.056 |
| ge5 | m_clim | glm_matched | 2,018 | -656.757 | -738.14 | 0.11 | NA |
| ge5 | m_clim | phyloglm_matched | 2,018 | -482.271 | -517.89 | 0.07 | 0.060 |
| ge5 | m_clim_breadth | glm_matched | 2,018 | -646.789 | -738.14 | 0.12 | NA |
| ge5 | m_clim_breadth | phyloglm_matched | 2,018 | -481.017 | -517.89 | 0.07 | 0.063 |
| ge5 | m_clim_island | glm_matched | 2,018 | -634.571 | -738.14 | 0.14 | NA |
| ge5 | m_clim_island | phyloglm_matched | 2,018 | -476.760 | -517.89 | 0.08 | 0.064 |
| ge5 | m_clim_mainland | glm_matched | 1,944 | -590.546 | -648.50 | 0.09 | NA |
| ge5 | m_clim_mainland | phyloglm_matched | 1,944 | -429.913 | -465.29 | 0.08 | 0.055 |
| ge10 | m_clim | glm_matched | 1,699 | -538.804 | -597.27 | 0.10 | NA |
| ge10 | m_clim | phyloglm_matched | 1,699 | -406.143 | -434.96 | 0.07 | 0.057 |
| ge10 | m_clim_breadth | glm_matched | 1,699 | -529.820 | -597.27 | 0.11 | NA |
| ge10 | m_clim_breadth | phyloglm_matched | 1,699 | -403.906 | -434.96 | 0.07 | 0.061 |
| ge10 | m_clim_island | glm_matched | 1,699 | -519.725 | -597.27 | 0.13 | NA |
| ge10 | m_clim_island | phyloglm_matched | 1,699 | -399.198 | -434.96 | 0.08 | 0.060 |
| ge10 | m_clim_mainland | glm_matched | 1,646 | -487.919 | -531.56 | 0.08 | NA |
| ge10 | m_clim_mainland | phyloglm_matched | 1,646 | -365.026 | -392.14 | 0.07 | 0.053 |

**Table S15. Pairwise correlations among climatic niche predictors across niche-data tiers**
Pairwise correlations among explanatory variables used in woodiness–climate models, calculated separately for each niche-data tier (ge1, ge5, ge10). Both Spearman's rank correlation (ρ) and Pearson's correlation (r) are reported, along with corresponding p-values and sample sizes (n). Correlations were generally moderate rather than extreme. In particular, drought (Maximum Cumulative Water Deficit; MCWD) and frost (minimum temperature of the coldest month; BIO6) were consistently negatively correlated, but not strongly so, indicating overlapping but non-redundant climatic gradients.

| tier | var1 | var2 | spearman_rho | spearman_p | spearman_n | pearson_r | pearson_p | pearson_n |
|---|---|---|---|---|---|---|---|---|
| ge1 | MCWD_med | BIO10_med | -0.616547415 | 0 | 3,417 | -0.639784153 | 0 | 3,417 |
| ge1 | MCWD_med | BIO6_med | -0.473886965 | 7.15E-191 | 3,417 | -0.493240256 | 5.22E-209 | 3,417 |
| ge1 | MCWD_med | elev_med | 0.049200971 | 0.004018089 | 3,417 | 0.129820921 | 2.57E-14 | 3,417 |
| ge1 | MCWD_med | MCWD_mad | -0.319949576 | 5.06E-70 | 2,900 | -0.314112877 | 2.00E-67 | 2,900 |
| ge1 | MCWD_med | BIO10_mad | 0.051728083 | 0.005331236 | 2,900 | 0.05087883 | 0.006134531 | 2,900 |
| ge1 | MCWD_med | BIO6_mad | 0.189723351 | 6.60E-25 | 2,900 | 0.191717251 | 2.08E-25 | 2,900 |
| ge1 | MCWD_med | elev_mad | 0.055924754 | 0.002589351 | 2,900 | 0.097356093 | 1.50E-07 | 2,900 |
| ge1 | MCWD_med | is_island | -0.071493425 | 2.88E-05 | 3,416 | -0.052089153 | 0.002323827 | 3,416 |
| ge1 | BIO10_med | BIO6_med | 0.618390669 | 0 | 3,417 | 0.621504771 | 0 | 3,417 |
| ge1 | BIO10_med | elev_med | -0.568221334 | 1.77E-291 | 3,417 | -0.628762605 | 0 | 3,417 |
| ge1 | BIO10_med | MCWD_mad | 0.182110733 | 4.81E-23 | 2,900 | 0.217789709 | 1.78E-32 | 2,900 |
| ge1 | BIO10_med | BIO10_mad | -0.180116012 | 1.44E-22 | 2,900 | -0.163477426 | 8.04E-19 | 2,900 |
| ge1 | BIO10_med | BIO6_mad | -0.163490085 | 7.99E-19 | 2,900 | -0.16321109 | 9.17E-19 | 2,900 |
| ge1 | BIO10_med | elev_mad | -0.267882076 | 7.59E-49 | 2,900 | -0.250017459 | 1.42E-42 | 2,900 |
| ge1 | BIO10_med | is_island | 0.06967638 | 4.59E-05 | 3,416 | 0.062999153 | 2.29E-04 | 3,416 |
| ge1 | BIO6_med | elev_med | -0.390115177 | 1.30E-124 | 3,417 | -0.320494936 | 1.80E-82 | 3,417 |
| ge1 | BIO6_med | MCWD_mad | 0.101179627 | 4.75E-08 | 2,900 | 0.149933951 | 4.79E-16 | 2,900 |
| ge1 | BIO6_med | BIO10_mad | -0.231091015 | 1.85E-36 | 2,900 | -0.177938924 | 4.67E-22 | 2,900 |
| ge1 | BIO6_med | BIO6_mad | -0.288070665 | 1.56E-56 | 2,900 | -0.320525037 | 2.79E-70 | 2,900 |
| ge1 | BIO6_med | elev_mad | -0.241917101 | 6.83E-40 | 2,900 | -0.194817635 | 3.38E-26 | 2,900 |
| ge1 | BIO6_med | is_island | 0.199682048 | 4.65E-32 | 3,416 | 0.192995329 | 5.08E-30 | 3,416 |
| ge1 | elev_med | MCWD_mad | 0.031852262 | 0.086346609 | 2,900 | -0.005861815 | 0.752355463 | 2,900 |
| ge1 | elev_med | BIO10_mad | 0.324597567 | 3.93E-72 | 2,900 | 0.321550406 | 9.59E-71 | 2,900 |
| ge1 | elev_med | BIO6_mad | 0.137839744 | 8.97E-14 | 2,900 | 0.08872633 | 1.71E-06 | 2,900 |
| ge1 | elev_med | elev_mad | 0.529888091 | 1.09E-209 | 2,900 | 0.427679258 | 2.62E-129 | 2,900 |
| ge1 | elev_med | is_island | -0.088078752 | 2.52E-07 | 3,416 | -0.097132004 | 1.28E-08 | 3,416 |
| ge1 | MCWD_mad | BIO10_mad | 0.566202497 | 1.52E-245 | 2,900 | 0.465394875 | 7.78E-156 | 2,900 |
| ge1 | MCWD_mad | BIO6_mad | 0.454362459 | 1.00E-147 | 2,900 | 0.313315438 | 4.48E-67 | 2,900 |
| ge1 | MCWD_mad | elev_mad | 0.410293019 | 3.62E-118 | 2,900 | 0.301169602 | 7.16E-62 | 2,900 |
| ge1 | MCWD_mad | is_island | -0.125898847 | 1.03E-11 | 2,899 | -0.105853792 | 1.11E-08 | 2,899 |
| ge1 | BIO10_mad | BIO6_mad | 0.717262964 | 0 | 2,900 | 0.625889274 | 0.00E-02 | 2,900 |
| ge1 | BIO10_mad | elev_mad | 0.726281871 | 0 | 2,900 | 0.742187633 | 0 | 2,900 |
| ge1 | BIO10_mad | is_island | -0.114253254 | 6.86E-10 | 2,899 | -0.102878686 | 2.83E-08 | 2,899 |
| ge1 | BIO6_mad | elev_mad | 0.616429781 | 3.72E-303 | 2,900 | 0.513834501 | 3.49E-195 | 2,900 |
| ge1 | BIO6_mad | is_island | -0.071708705 | 1.11E-04 | 2,899 | -0.074341381 | 6.16E-05 | 2,899 |
| ge1 | elev_mad | is_island | -0.039316405 | 0.03427737 | 2,899 | -0.049838063 | 0.007276677 | 2,899 |
| ge5 | MCWD_med | BIO10_med | -0.617748644 | 8.65E-274 | 2,603 | -0.652130721 | 0.00E-02 | 2,603 |
| ge5 | MCWD_med | BIO6_med | -0.498420165 | 1.58E-163 | 2,603 | -0.51398835 | 1.65E-175 | 2,603 |
| ge5 | MCWD_med | elev_med | 0.004827525 | 0.805541685 | 2,603 | 0.094504805 | 1.36E-06 | 2,603 |
| ge5 | MCWD_med | MCWD_mad | -0.352786067 | 3.77E-77 | 2,603 | -0.336344483 | 7.24E-70 | 2,603 |
| ge5 | MCWD_med | BIO10_mad | 0.048046275 | 0.014224625 | 2,603 | 0.052845878 | 0.007001663 | 2,603 |
| ge5 | MCWD_med | BIO6_mad | 0.193110629 | 2.75E-23 | 2,603 | 0.196233262 | 5.23E-24 | 2,603 |
| ge5 | MCWD_med | elev_mad | 0.053833196 | 0.006010257 | 2,603 | 0.098541951 | 4.72E-07 | 2,603 |
| ge5 | MCWD_med | is_island | -0.074211043 | 1.51E-04 | 2,602 | -0.05265183 | 0.00722412 | 2,602 |
| ge5 | BIO10_med | BIO6_med | 0.647368433 | 0.00E+00 | 2,603 | 0.652049174 | 0.00E-02 | 2,603 |
| ge5 | BIO10_med | elev_med | -0.53225194 | 2.19E-190 | 2,603 | -0.580107494 | 5.17E-234 | 2,603 |
| ge5 | BIO10_med | MCWD_mad | 0.197167952 | 3.17E-24 | 2,603 | 0.228816533 | 2.87E-32 | 2,603 |
| ge5 | BIO10_med | BIO10_mad | -0.198220289 | 1.79E-24 | 2,603 | -0.182632439 | 5.86E-21 | 2,603 |
| ge5 | BIO10_med | BIO6_mad | -0.174981712 | 2.41E-19 | 2,603 | -0.169772223 | 2.76E-18 | 2,603 |
| ge5 | BIO10_med | elev_mad | -0.296637071 | 4.99E-54 | 2,603 | -0.272938626 | 1.08E-45 | 2,603 |
| ge5 | BIO10_med | is_island | 0.052525396 | 0.007365018 | 2,602 | 0.045250478 | 0.020982946 | 2,602 |
| ge5 | BIO6_med | elev_med | -0.378299161 | 2.42E-89 | 2,603 | -0.30095893 | 1.23E-55 | 2,603 |
| ge5 | BIO6_med | MCWD_mad | 0.112605292 | 8.38E-09 | 2,603 | 0.16127246 | 1.25E-16 | 2,603 |
| ge5 | BIO6_med | BIO10_mad | -0.247569866 | 1.19E-37 | 2,603 | -0.196819663 | 3.82E-24 | 2,603 |
| ge5 | BIO6_med | BIO6_mad | -0.307752316 | 3.21E-58 | 2,603 | -0.33697176 | 3.89E-70 | 2,603 |
| ge5 | BIO6_med | elev_mad | -0.262465648 | 2.88E-42 | 2,603 | -0.210289233 | 2.10E-27 | 2,603 |
| ge5 | BIO6_med | is_island | 0.183726769 | 3.46E-21 | 2,602 | 0.171664931 | 1.17E-18 | 2,602 |
| ge5 | elev_med | MCWD_mad | 0.060057928 | 0.002173593 | 2,603 | 0.016087076 | 0.411979497 | 2,603 |
| ge5 | elev_med | BIO10_mad | 0.366272017 | 1.86E-83 | 2,603 | 0.367722441 | 3.75E-84 | 2,603 |
| ge5 | elev_med | BIO6_mad | 0.15520339 | 1.68E-15 | 2,603 | 0.09491604 | 1.23E-06 | 2,603 |
| ge5 | elev_med | elev_mad | 0.586928515 | 7.78E-241 | 2,603 | 0.474338087 | 3.61E-146 | 2,603 |
| ge5 | elev_med | is_island | -0.052627039 | 0.007251556 | 2,602 | -0.072191367 | 2.28E-04 | 2,602 |
| ge5 | MCWD_mad | BIO10_mad | 0.546537677 | 1.13E-202 | 2,603 | 0.457142865 | 1.34E-134 | 2,603 |
| ge5 | MCWD_mad | BIO6_mad | 0.422715472 | 2.43E-113 | 2,603 | 0.293616132 | 6.39E-53 | 2,603 |
| ge5 | MCWD_mad | elev_mad | 0.383822834 | 3.95E-92 | 2,603 | 0.28569344 | 4.43E-50 | 2,603 |
| ge5 | MCWD_mad | is_island | -0.130361235 | 2.47E-11 | 2,602 | -0.109685146 | 2.03E-08 | 2,602 |
| ge5 | BIO10_mad | BIO6_mad | 0.699190979 | 0 | 2,603 | 0.603827983 | 1.86E-258 | 2,603 |
| ge5 | BIO10_mad | elev_mad | 0.715214391 | 0 | 2,603 | 0.738159891 | 0 | 2,603 |
| ge5 | BIO10_mad | is_island | -0.109478885 | 2.16E-08 | 2,602 | -0.099206175 | 3.97E-07 | 2,602 |
| ge5 | BIO6_mad | elev_mad | 0.593170789 | 3.19E-247 | 2,603 | 0.485391813 | 5.79E-154 | 2,603 |
| ge5 | BIO6_mad | is_island | -0.065836925 | 7.78E-04 | 2,602 | -0.069910379 | 3.59E-04 | 2,602 |
| ge5 | elev_mad | is_island | -0.025479791 | 0.193839102 | 2,602 | -0.039822882 | 0.042236029 | 2,602 |
| ge10 | MCWD_med | BIO10_med | -0.610497883 | 1.05E-220 | 2,159 | -0.646855235 | 3.58E-256 | 2,159 |
| ge10 | MCWD_med | BIO6_med | -0.505496314 | 2.09E-140 | 2,159 | -0.517727168 | 2.34E-148 | 2,159 |
| ge10 | MCWD_med | elev_med | -0.033322052 | 0.121659948 | 2,159 | 0.059588316 | 0.005611978 | 2,159 |
| ge10 | MCWD_med | MCWD_mad | -0.383578817 | 1.28E-76 | 2,159 | -0.360028062 | 4.50E-67 | 2,159 |
| ge10 | MCWD_med | BIO10_mad | 0.046228143 | 0.031721948 | 2,159 | 0.051192379 | 0.017367296 | 2,159 |
| ge10 | MCWD_med | BIO6_mad | 0.218134405 | 1.14E-24 | 2,159 | 0.217732283 | 1.39E-24 | 2,159 |
| ge10 | MCWD_med | elev_mad | 0.048403696 | 0.024506419 | 2,159 | 0.097211203 | 6.04E-06 | 2,159 |
| ge10 | MCWD_med | is_island | -0.050164697 | 0.019780696 | 2,158 | -0.029503094 | 0.170671585 | 2,158 |
| ge10 | BIO10_med | BIO6_med | 0.672843827 | 1.17E-284 | 2,159 | 0.680106783 | 3.77E-293 | 2,159 |
| ge10 | BIO10_med | elev_med | -0.507337386 | 1.39E-141 | 2,159 | -0.558792145 | 1.44E-177 | 2,159 |
| ge10 | BIO10_med | MCWD_mad | 0.213808767 | 9.64E-24 | 2,159 | 0.244942731 | 7.29E-31 | 2,159 |
| ge10 | BIO10_med | BIO10_mad | -0.204900748 | 6.75E-22 | 2,159 | -0.189849402 | 5.72E-19 | 2,159 |
| ge10 | BIO10_med | BIO6_mad | -0.183832666 | 7.29E-18 | 2,159 | -0.179434171 | 4.43E-17 | 2,159 |
| ge10 | BIO10_med | elev_mad | -0.314319522 | 1.03E-50 | 2,159 | -0.290567468 | 2.85E-43 | 2,159 |
| ge10 | BIO10_med | is_island | 0.047585294 | 0.027069651 | 2,158 | 0.039568658 | 0.066093628 | 2,158 |
| ge10 | BIO6_med | elev_med | -0.377507008 | 4.40E-74 | 2,159 | -0.298959832 | 8.02E-46 | 2,159 |
| ge10 | BIO6_med | MCWD_mad | 0.137426388 | 1.43E-10 | 2,159 | 0.188486126 | 1.03E-18 | 2,159 |
| ge10 | BIO6_med | BIO10_mad | -0.260451171 | 8.21E-35 | 2,159 | -0.216916355 | 2.09E-24 | 2,159 |
| ge10 | BIO6_med | BIO6_mad | -0.339137335 | 2.98E-59 | 2,159 | -0.377653692 | 3.83E-74 | 2,159 |
| ge10 | BIO6_med | elev_mad | -0.277795207 | 1.49E-39 | 2,159 | -0.235523441 | 1.35E-28 | 2,159 |
| ge10 | BIO6_med | is_island | 0.166925979 | 5.96E-15 | 2,158 | 0.153177158 | 8.44E-13 | 2,158 |
| ge10 | elev_med | MCWD_mad | 0.077195527 | 3.30E-04 | 2,159 | 0.028803845 | 0.180937411 | 2,159 |
| ge10 | elev_med | BIO10_mad | 0.373182734 | 2.62E-72 | 2,159 | 0.374942466 | 5.00E-73 | 2,159 |
| ge10 | elev_med | BIO6_mad | 0.128099823 | 2.32E-09 | 2,159 | 0.060072334 | 0.005235605 | 2,159 |
| ge10 | elev_med | elev_mad | 0.627847199 | 4.84E-237 | 2,159 | 0.499464149 | 1.33E-136 | 2,159 |
| ge10 | elev_med | is_island | -0.047102876 | 0.028664005 | 2,158 | -0.066378586 | 0.002034251 | 2,158 |
| ge10 | MCWD_mad | BIO10_mad | 0.542178501 | 2.91E-165 | 2,159 | 0.45990903 | 1.83E-113 | 2,159 |
| ge10 | MCWD_mad | BIO6_mad | 0.401138327 | 2.86E-84 | 2,159 | 0.278772385 | 7.85E-40 | 2,159 |
| ge10 | MCWD_mad | elev_mad | 0.372286223 | 6.07E-72 | 2,159 | 0.282715361 | 5.80E-41 | 2,159 |
| ge10 | MCWD_mad | is_island | -0.121547197 | 1.48E-08 | 2,158 | -0.10006015 | 3.21E-06 | 2,158 |
| ge10 | BIO10_mad | BIO6_mad | 0.672948934 | 8.82E-285 | 2,159 | 0.568718068 | 2.92E-185 | 2,159 |
| ge10 | BIO10_mad | elev_mad | 0.689845632 | 6.25E-305 | 2,159 | 0.716867639 | 0 | 2,159 |
| ge10 | BIO10_mad | is_island | -0.11829183 | 3.56E-08 | 2,158 | -0.102875462 | 1.68E-06 | 2,158 |
| ge10 | BIO6_mad | elev_mad | 0.546646453 | 1.67E-168 | 2,159 | 0.435849967 | 8.24E-101 | 2,159 |
| ge10 | BIO6_mad | is_island | -0.073708603 | 6.11E-04 | 2,158 | -0.071173067 | 9.38E-04 | 2,158 |
| ge10 | elev_mad | is_island | -0.030153394 | 0.161435182 | 2,158 | -0.039165963 | 0.068900702 | 2,158 |

**Table S16. Variance inflation factors for predictors in woodiness–climate models**
Variance inflation factors (VIF), tolerance (1 – $R^2$), and predictor-specific $R^2$ values calculated for each explanatory variable across niche-data tiers (ge1, ge5, ge10), dataset variants (full and tree-matched subsets), and model formulations. VIF values remained low across all analyses (all < 4.4), indicating no strong multicollinearity among predictors. The focal variables drought (MCWD) and frost (BIO6) showed only modest collinearity, supporting their joint inclusion in generalized linear and phylogenetic models.

| tier | dataset | model | term | n | r2_against_others | tolerance | vif | vif_flag |
|---|---|---|---|---|---|---|---|---|
| ge1 | glm_full_input | m_clim | MCWD_med | 3,412 | 0.538813233 | 0.461186767 | 2.168318935 | low |
| ge1 | glm_full_input | m_clim | BIO10_med | 3,412 | 0.751022024 | 0.248977976 | 4.016419505 | low |
| ge1 | glm_full_input | m_clim | BIO6_med | 3,412 | 0.403685677 | 0.596314323 | 1.676967936 | low |
| ge1 | glm_full_input | m_clim | elev_med | 3,412 | 0.522377082 | 0.477622918 | 2.093701877 | low |
| ge1 | matched_input | m_clim | MCWD_med | 2,527 | 0.577626076 | 0.422373924 | 2.367570397 | low |
| ge1 | matched_input | m_clim | BIO10_med | 2,527 | 0.760975543 | 0.239024457 | 4.183672298 | low |
| ge1 | matched_input | m_clim | BIO6_med | 2,527 | 0.430924838 | 0.569075162 | 1.757237123 | low |
| ge1 | matched_input | m_clim | elev_med | 2,527 | 0.512588526 | 0.487411474 | 2.051654613 | low |
| ge1 | glm_full_input | m_clim_breadth | MCWD_med | 2,896 | 0.577868329 | 0.422131671 | 2.368929099 | low |
| ge1 | glm_full_input | m_clim_breadth | BIO10_med | 2,896 | 0.743902433 | 0.256097567 | 3.904761815 | low |
| ge1 | glm_full_input | m_clim_breadth | BIO6_med | 2,896 | 0.472812591 | 0.527187409 | 1.896858657 | low |
| ge1 | glm_full_input | m_clim_breadth | elev_med | 2,896 | 0.569870318 | 0.430129682 | 2.324880244 | low |
| ge1 | glm_full_input | m_clim_breadth | MCWD_mad | 2,896 | 0.359615459 | 0.640384541 | 1.561561743 | low |
| ge1 | glm_full_input | m_clim_breadth | BIO10_mad | 2,896 | 0.679975242 | 0.320024758 | 3.124758242 | low |
| ge1 | glm_full_input | m_clim_breadth | BIO6_mad | 2,896 | 0.491089117 | 0.508910883 | 1.964980574 | low |
| ge1 | glm_full_input | m_clim_breadth | elev_mad | 2,896 | 0.601474999 | 0.398525001 | 2.509252861 | low |
| ge1 | matched_input | m_clim_breadth | MCWD_med | 2,217 | 0.610810139 | 0.389189861 | 2.569440009 | low |
| ge1 | matched_input | m_clim_breadth | BIO10_med | 2,217 | 0.759073855 | 0.240926145 | 4.150649569 | low |
| ge1 | matched_input | m_clim_breadth | BIO6_med | 2,217 | 0.495107423 | 0.504892577 | 1.980619336 | low |
| ge1 | matched_input | m_clim_breadth | elev_med | 2,217 | 0.568789108 | 0.431210892 | 2.319050881 | low |
| ge1 | matched_input | m_clim_breadth | MCWD_mad | 2,217 | 0.371493806 | 0.628506194 | 1.591074217 | low |
| ge1 | matched_input | m_clim_breadth | BIO10_mad | 2,217 | 0.66186929 | 0.33813071 | 2.957436195 | low |
| ge1 | matched_input | m_clim_breadth | BIO6_mad | 2,217 | 0.494485745 | 0.505514255 | 1.978183581 | low |
| ge1 | matched_input | m_clim_breadth | elev_mad | 2,217 | 0.588372882 | 0.411627118 | 2.429383186 | low |
| ge1 | glm_full_input | m_clim_island | MCWD_med | 3,412 | 0.539260597 | 0.460739403 | 2.170424307 | low |
| ge1 | glm_full_input | m_clim_island | BIO10_med | 3,412 | 0.754134696 | 0.245865304 | 4.067267654 | low |
| ge1 | glm_full_input | m_clim_island | BIO6_med | 3,412 | 0.428225927 | 0.571774073 | 1.748942542 | low |
| ge1 | glm_full_input | m_clim_island | elev_med | 3,412 | 0.527428548 | 0.472571452 | 2.116082122 | low |
| ge1 | glm_full_input | m_clim_island | is_island | 3,412 | 0.053120529 | 0.946879471 | 1.056100624 | low |
| ge1 | matched_input | m_clim_island | MCWD_med | 2,527 | 0.57799984 | 0.42200016 | 2.36966735 | low |
| ge1 | matched_input | m_clim_island | BIO10_med | 2,527 | 0.764874753 | 0.235125247 | 4.253052408 | low |
| ge1 | matched_input | m_clim_island | BIO6_med | 2,527 | 0.460270004 | 0.539729996 | 1.852778254 | low |
| ge1 | matched_input | m_clim_island | elev_med | 2,527 | 0.518165913 | 0.481834087 | 2.075403188 | low |
| ge1 | matched_input | m_clim_island | is_island | 2,527 | 0.065489493 | 0.934510507 | 1.070078927 | low |
| ge1 | glm_full_input | m_clim_mainland | MCWD_med | 3,299 | 0.540846728 | 0.459153272 | 2.177921973 | low |
| ge1 | glm_full_input | m_clim_mainland | BIO10_med | 3,299 | 0.75632956 | 0.24367044 | 4.103903618 | low |
| ge1 | glm_full_input | m_clim_mainland | BIO6_med | 3,299 | 0.405984842 | 0.594015158 | 1.683458724 | low |
| ge1 | glm_full_input | m_clim_mainland | elev_med | 3,299 | 0.527229424 | 0.472770576 | 2.115190859 | low |
| ge1 | matched_input | m_clim_mainland | MCWD_med | 2,429 | 0.57834979 | 0.42165021 | 2.371634064 | low |
| ge1 | matched_input | m_clim_mainland | BIO10_med | 2,429 | 0.7671629 | 0.2328371 | 4.294848204 | low |
| ge1 | matched_input | m_clim_mainland | BIO6_med | 2,429 | 0.434690745 | 0.565309255 | 1.768943266 | low |
| ge1 | matched_input | m_clim_mainland | elev_med | 2,429 | 0.517468467 | 0.482531533 | 2.072403421 | low |
| ge5 | glm_full_input | m_clim | MCWD_med | 2,601 | 0.552125178 | 0.447874822 | 2.232766724 | low |
| ge5 | glm_full_input | m_clim | BIO10_med | 2,601 | 0.74731235 | 0.25268765 | 3.957454977 | low |
| ge5 | glm_full_input | m_clim | BIO6_med | 2,601 | 0.441431316 | 0.558568684 | 1.790290125 | low |
| ge5 | glm_full_input | m_clim | elev_med | 2,601 | 0.478826858 | 0.521173142 | 1.918748146 | low |
| ge5 | matched_input | m_clim | MCWD_med | 2,018 | 0.591197467 | 0.408802533 | 2.446168796 | low |
| ge5 | matched_input | m_clim | BIO10_med | 2,018 | 0.763946747 | 0.236053253 | 4.23633222 | low |
| ge5 | matched_input | m_clim | BIO6_med | 2,018 | 0.468807337 | 0.531192663 | 1.882556121 | low |
| ge5 | matched_input | m_clim | elev_med | 2,018 | 0.478056938 | 0.521943062 | 1.915917794 | low |
| ge5 | glm_full_input | m_clim_breadth | MCWD_med | 2,601 | 0.58875226 | 0.41124774 | 2.431624308 | low |
| ge5 | glm_full_input | m_clim_breadth | BIO10_med | 2,601 | 0.752554457 | 0.247445543 | 4.041293238 | low |
| ge5 | glm_full_input | m_clim_breadth | BIO6_med | 2,601 | 0.49573227 | 0.50426773 | 1.983073554 | low |
| ge5 | glm_full_input | m_clim_breadth | elev_med | 2,601 | 0.586072135 | 0.413927865 | 2.415879875 | low |
| ge5 | glm_full_input | m_clim_breadth | MCWD_mad | 2,601 | 0.368450874 | 0.631549126 | 1.583408098 | low |
| ge5 | glm_full_input | m_clim_breadth | BIO10_mad | 2,601 | 0.677455027 | 0.322544973 | 3.100342845 | low |
| ge5 | glm_full_input | m_clim_breadth | BIO6_mad | 2,601 | 0.474039165 | 0.525960835 | 1.901282251 | low |
| ge5 | glm_full_input | m_clim_breadth | elev_mad | 2,601 | 0.602192843 | 0.397807157 | 2.51378082 | low |
| ge5 | matched_input | m_clim_breadth | MCWD_med | 2,018 | 0.619657398 | 0.380342602 | 2.629208493 | low |
| ge5 | matched_input | m_clim_breadth | BIO10_med | 2,018 | 0.767604552 | 0.232395448 | 4.303010274 | low |
| ge5 | matched_input | m_clim_breadth | BIO6_med | 2,018 | 0.52246492 | 0.47753508 | 2.094086994 | low |
| ge5 | matched_input | m_clim_breadth | elev_med | 2,018 | 0.580302536 | 0.419697464 | 2.382668676 | low |
| ge5 | matched_input | m_clim_breadth | MCWD_mad | 2,018 | 0.370391522 | 0.629608478 | 1.588288651 | low |
| ge5 | matched_input | m_clim_breadth | BIO10_mad | 2,018 | 0.656330624 | 0.343669376 | 2.909773374 | low |
| ge5 | matched_input | m_clim_breadth | BIO6_mad | 2,018 | 0.4835019 | 0.5164981 | 1.936115545 | low |
| ge5 | matched_input | m_clim_breadth | elev_mad | 2,018 | 0.586802397 | 0.413197603 | 2.420149566 | low |
| ge5 | glm_full_input | m_clim_island | MCWD_med | 2,601 | 0.552950877 | 0.447049123 | 2.236890642 | low |
| ge5 | glm_full_input | m_clim_island | BIO10_med | 2,601 | 0.750947638 | 0.249052362 | 4.015219893 | low |
| ge5 | glm_full_input | m_clim_island | BIO6_med | 2,601 | 0.461890145 | 0.538109855 | 1.858356598 | low |
| ge5 | glm_full_input | m_clim_island | elev_med | 2,601 | 0.483280394 | 0.516719606 | 1.935285574 | low |
| ge5 | glm_full_input | m_clim_island | is_island | 2,601 | 0.045546363 | 0.954453637 | 1.047719828 | low |
| ge5 | matched_input | m_clim_island | MCWD_med | 2,018 | 0.59152583 | 0.40847417 | 2.448135213 | low |
| ge5 | matched_input | m_clim_island | BIO10_med | 2,018 | 0.767591577 | 0.232408423 | 4.302770049 | low |
| ge5 | matched_input | m_clim_island | BIO6_med | 2,018 | 0.493090027 | 0.506909973 | 1.972736884 | low |
| ge5 | matched_input | m_clim_island | elev_med | 2,018 | 0.482574953 | 0.517425047 | 1.932647068 | low |
| ge5 | matched_input | m_clim_island | is_island | 2,018 | 0.055119182 | 0.944880818 | 1.058334533 | low |
| ge5 | glm_full_input | m_clim_mainland | MCWD_med | 2,520 | 0.552234588 | 0.447765412 | 2.233312297 | low |
| ge5 | glm_full_input | m_clim_mainland | BIO10_med | 2,520 | 0.753285349 | 0.246714651 | 4.053265574 | low |
| ge5 | glm_full_input | m_clim_mainland | BIO6_med | 2,520 | 0.446382812 | 0.553617188 | 1.806302299 | low |
| ge5 | glm_full_input | m_clim_mainland | elev_med | 2,520 | 0.484509778 | 0.515490222 | 1.939901005 | low |
| ge5 | matched_input | m_clim_mainland | MCWD_med | 1,944 | 0.592005495 | 0.407994505 | 2.4510134 | low |
| ge5 | matched_input | m_clim_mainland | BIO10_med | 1,944 | 0.770590323 | 0.229409677 | 4.359014018 | low |
| ge5 | matched_input | m_clim_mainland | BIO6_med | 1,944 | 0.475451315 | 0.524548685 | 1.906400739 | low |
| ge5 | matched_input | m_clim_mainland | elev_med | 1,944 | 0.483851056 | 0.516148944 | 1.937425256 | low |
| ge10 | glm_full_input | m_clim | MCWD_med | 2,158 | 0.554037237 | 0.445962763 | 2.242339681 | low |
| ge10 | glm_full_input | m_clim | BIO10_med | 2,158 | 0.753805926 | 0.246194074 | 4.061836198 | low |
| ge10 | glm_full_input | m_clim | BIO6_med | 2,158 | 0.476317439 | 0.523682561 | 1.909553753 | low |
| ge10 | glm_full_input | m_clim | elev_med | 2,158 | 0.472091022 | 0.527908978 | 1.894265948 | low |
| ge10 | matched_input | m_clim | MCWD_med | 1,699 | 0.585187273 | 0.414812727 | 2.410726421 | low |
| ge10 | matched_input | m_clim | BIO10_med | 1,699 | 0.767562807 | 0.232437193 | 4.30223746 | low |
| ge10 | matched_input | m_clim | BIO6_med | 1,699 | 0.506219238 | 0.493780762 | 2.025190282 | low |
| ge10 | matched_input | m_clim | elev_med | 1,699 | 0.463243073 | 0.536756927 | 1.863040698 | low |
| ge10 | glm_full_input | m_clim_breadth | MCWD_med | 2,158 | 0.595883601 | 0.404116399 | 2.474534573 | low |
| ge10 | glm_full_input | m_clim_breadth | BIO10_med | 2,158 | 0.760121207 | 0.239878793 | 4.168772016 | low |
| ge10 | glm_full_input | m_clim_breadth | BIO6_med | 2,158 | 0.548136746 | 0.451863254 | 2.213058907 | low |
| ge10 | glm_full_input | m_clim_breadth | elev_med | 2,158 | 0.592839359 | 0.407160641 | 2.456033071 | low |
| ge10 | glm_full_input | m_clim_breadth | MCWD_mad | 2,158 | 0.396914248 | 0.603085752 | 1.658138989 | low |
| ge10 | glm_full_input | m_clim_breadth | BIO10_mad | 2,158 | 0.655808204 | 0.344191796 | 2.905356872 | low |
| ge10 | glm_full_input | m_clim_breadth | BIO6_mad | 2,158 | 0.470441928 | 0.529558072 | 1.888367025 | low |
| ge10 | glm_full_input | m_clim_breadth | elev_mad | 2,158 | 0.58507029 | 0.41492971 | 2.410046753 | low |
| ge10 | matched_input | m_clim_breadth | MCWD_med | 1,699 | 0.618080474 | 0.381919526 | 2.618352643 | low |
| ge10 | matched_input | m_clim_breadth | BIO10_med | 1,699 | 0.77238819 | 0.22761181 | 4.393445143 | low |
| ge10 | matched_input | m_clim_breadth | BIO6_med | 1,699 | 0.57266433 | 0.42733567 | 2.340080809 | low |
| ge10 | matched_input | m_clim_breadth | elev_med | 1,699 | 0.581400228 | 0.418599772 | 2.388916733 | low |
| ge10 | matched_input | m_clim_breadth | MCWD_mad | 1,699 | 0.377365538 | 0.622634462 | 1.606078784 | low |
| ge10 | matched_input | m_clim_breadth | BIO10_mad | 1,699 | 0.63507612 | 0.36492388 | 2.740297508 | low |
| ge10 | matched_input | m_clim_breadth | BIO6_mad | 1,699 | 0.485375423 | 0.514624577 | 1.943164095 | low |
| ge10 | matched_input | m_clim_breadth | elev_mad | 1,699 | 0.572280355 | 0.427719645 | 2.337980057 | low |
| ge10 | glm_full_input | m_clim_island | MCWD_med | 2,158 | 0.554206677 | 0.445793323 | 2.243191962 | low |
| ge10 | glm_full_input | m_clim_island | BIO10_med | 2,158 | 0.75645465 | 0.24354535 | 4.106011461 | low |
| ge10 | glm_full_input | m_clim_island | BIO6_med | 2,158 | 0.493127171 | 0.506872829 | 1.972881447 | low |
| ge10 | glm_full_input | m_clim_island | elev_med | 2,158 | 0.475331316 | 0.524668684 | 1.905964717 | low |
| ge10 | glm_full_input | m_clim_island | is_island | 2,158 | 0.037546047 | 0.962453953 | 1.039010747 | low |
| ge10 | matched_input | m_clim_island | MCWD_med | 1,699 | 0.585202129 | 0.414797871 | 2.410812758 | low |
| ge10 | matched_input | m_clim_island | BIO10_med | 1,699 | 0.770187257 | 0.229812743 | 4.351368801 | low |
| ge10 | matched_input | m_clim_island | BIO6_med | 1,699 | 0.525238623 | 0.474761377 | 2.106321297 | low |
| ge10 | matched_input | m_clim_island | elev_med | 1,699 | 0.466285236 | 0.533714764 | 1.873659992 | low |
| ge10 | matched_input | m_clim_island | is_island | 1,699 | 0.044125121 | 0.955874879 | 1.046162026 | low |
| ge10 | glm_full_input | m_clim_mainland | MCWD_med | 2,101 | 0.554494728 | 0.445505272 | 2.244642347 | low |
| ge10 | glm_full_input | m_clim_mainland | BIO10_med | 2,101 | 0.758649097 | 0.241350903 | 4.143344764 | low |
| ge10 | glm_full_input | m_clim_mainland | BIO6_med | 2,101 | 0.482952699 | 0.517047301 | 1.934059027 | low |
| ge10 | glm_full_input | m_clim_mainland | elev_med | 2,101 | 0.476941379 | 0.523058621 | 1.911831598 | low |
| ge10 | matched_input | m_clim_mainland | MCWD_med | 1,646 | 0.586402491 | 0.413597509 | 2.417809533 | low |
| ge10 | matched_input | m_clim_mainland | BIO10_med | 1,646 | 0.772921805 | 0.227078195 | 4.40376938 | low |
| ge10 | matched_input | m_clim_mainland | BIO6_med | 1,646 | 0.514524193 | 0.485475807 | 2.05983488 | low |
| ge10 | matched_input | m_clim_mainland | elev_med | 1,646 | 0.468006985 | 0.531993015 | 1.879723928 | low |

**Table S17. Comparison of global and supertribe-specific woodiness–climate models**

Comparison of non-phylogenetic logistic regression models testing whether the relationship between growth form (woody vs. herbaceous) and climatic predictors differs among supertribes in Brassicaceae. Analyses are restricted to the ge10 dataset (species with ≥10 retained occurrence grid cells) and include only supertribes with a minimum sample size threshold.

The base model includes additive effects of drought (MCWD), frost (BIO6), heat (BIO10), and elevation, whereas the interaction model additionally allows slopes for each predictor to vary among supertribes. Model fit is compared using Akaike Information Criterion (AIC), likelihood ratio tests (LRT), and McFadden's pseudo-$R^2$.

Reported values include the number of species (n), number of supertribes retained (considering tribe Aethionemeae and unplaced species as two additional supertribes), minimum per-supertribe sample size, AIC for base and interaction models, ΔAIC (interaction – base), LRT statistics (degrees of freedom, likelihood ratio, and p-value), and pseudo-$R^2$ values with their difference ($\Delta R^2$). Negative ΔAIC and positive $\Delta R^2$ indicate improved model fit when allowing supertribe-specific climate responses.

| tier | n | n_supertribe | min_n_supertribe | glm_AIC_base | glm_AIC_int | glm_deltaAIC | glm_LRT_df | glm_LRT_LR | glm_LRT_p | glm_R2_base | glm_R2_int | glm_deltaR2 |
|---|---|---|---|---|---|---|---|---|---|---|---|---|
| ge10 | 2157 | 7 | 15 | 1,173.90 | 1,104.52 | -69.3786 | 30 | 129.38 | 2.63E-14 | 0.1078 | 0.2070 | 0.0992 |

#### Table S18. Coefficients of the ge10 supertribe interaction model

Regression coefficients from the ge10 interaction model (GLM) including supertribe-specific climate effects. The table reports main effects and interaction terms for drought (MCWD), frost (BIO6), heat (BIO10), and elevation, along with standard errors, confidence intervals, and P-values. Coefficients reflect deviations from the overall effect under sum contrasts.

| term | estimate | se | z | p | conf.low | conf.high | OR | OR_lo | OR_hi | logLik | AIC | tier | fit |
|---|---|---|---|---|---|---|---|---|---|---|---|---|---|
| (Intercept) | -2.679 | 0.10 | -26.491 | 0.00000 | -2.88 | -2.48 | 6.86E-02 | 5.63E-02 | 8.37E-02 | -581.95 | 1,173.90 | ge10 | base |
| MCWD_med | -0.438 | 0.11 | -3.903 | 0.00009 | -0.66 | -0.22 | 6.45E-01 | 5.18E-01 | 8.04E-01 | -581.95 | 1,173.90 | ge10 | base |
| BIO6_med | 0.902 | 0.14 | 6.621 | 0.00000 | 0.63 | 1.17 | 2.46E+00 | 1.89E+00 | 3.22E+00 | -581.95 | 1,173.90 | ge10 | base |
| BIO10_med | -0.266 | 0.17 | -1.572 | 0.11597 | -0.60 | 0.07 | 7.66E-01 | 5.50E-01 | 1.07E+00 | -581.95 | 1,173.90 | ge10 | base |
| elev_med | 0.018 | 0.13 | 0.139 | 0.88906 | -0.23 | 0.27 | 1.02E+00 | 7.94E-01 | 1.31E+00 | -581.95 | 1,173.90 | ge10 | base |
| (Intercept) | -4.590 | 162.14 | -0.028 | 0.97742 | -322.38 | 313.20 | 1.02E-02 | 9.85E-141 | 1.05E+136 | -517.26 | 1,104.52 | ge10 | interaction |
| MCWD_med | 0.228 | 254.22 | 0.001 | 0.99928 | -498.05 | 498.51 | 1.26E+00 | 5.01E-217 | 3.15E+216 | -517.26 | 1,104.52 | ge10 | interaction |
| BIO6_med | -0.179 | 155.23 | -0.001 | 0.99908 | -304.43 | 304.08 | 8.36E-01 | 6.11E-133 | 1.15E+132 | -517.26 | 1,104.52 | ge10 | interaction |
| BIO10_med | 0.339 | 231.86 | 0.001 | 0.99884 | -454.12 | 454.79 | 1.40E+00 | 6.03E-198 | 3.27E+197 | -517.26 | 1,104.52 | ge10 | interaction |
| elev_med | -0.196 | 153.06 | -0.001 | 0.99898 | -300.19 | 299.80 | 8.22E-01 | 4.25E-131 | 1.59E+130 | -517.26 | 1,104.52 | ge10 | interaction |
| SUPERTRIBE1 | 3.651 | 162.14 | 0.023 | 0.98204 | -314.15 | 321.45 | 3.85E+01 | 3.69E-137 | 4.02E+139 | -517.26 | 1,104.52 | ge10 | interaction |
| SUPERTRIBE2 | 2.459 | 162.14 | 0.015 | 0.98790 | -315.33 | 320.25 | 1.17E+01 | 1.13E-137 | 1.21E+139 | -517.26 | 1,104.52 | ge10 | interaction |
| SUPERTRIBE3 | 1.725 | 162.14 | 0.011 | 0.99151 | -316.06 | 319.51 | 5.61E+00 | 5.44E-138 | 5.79E+138 | -517.26 | 1,104.52 | ge10 | interaction |
| SUPERTRIBE4 | 0.897 | 162.14 | 0.006 | 0.99558 | -316.89 | 318.68 | 2.45E+00 | 2.38E-138 | 2.53E+138 | -517.26 | 1,104.52 | ge10 | interaction |
| SUPERTRIBE5 | 1.230 | 162.14 | 0.008 | 0.99395 | -316.56 | 319.02 | 3.42E+00 | 3.32E-138 | 3.53E+138 | -517.26 | 1,104.52 | ge10 | interaction |
| SUPERTRIBE6 | 2.014 | 162.14 | 0.012 | 0.99009 | -315.77 | 319.80 | 7.49E+00 | 7.26E-138 | 7.73E+138 | -517.26 | 1,104.52 | ge10 | interaction |
| MCWD_med:SUPERTRIBE1 | 3.196 | 254.26 | 0.013 | 0.98997 | -495.14 | 501.54 | 2.44E+01 | 9.15E-216 | 6.53E+217 | -517.26 | 1,104.52 | ge10 | interaction |
| MCWD_med:SUPERTRIBE2 | -0.051 | 254.22 | 0.000 | 0.99984 | -498.33 | 498.23 | 9.50E-01 | 3.78E-217 | 2.39E+216 | -517.26 | 1,104.52 | ge10 | interaction |
| MCWD_med:SUPERTRIBE3 | -0.960 | 254.22 | -0.004 | 0.99699 | -499.24 | 497.32 | 3.83E-01 | 1.53E-217 | 9.62E+215 | -517.26 | 1,104.52 | ge10 | interaction |
| MCWD_med:SUPERTRIBE4 | -0.899 | 254.22 | -0.004 | 0.99718 | -499.18 | 497.38 | 4.07E-01 | 1.62E-217 | 1.02E+216 | -517.26 | 1,104.52 | ge10 | interaction |
| MCWD_med:SUPERTRIBE5 | -0.337 | 254.22 | -0.001 | 0.99894 | -498.62 | 497.94 | 7.14E-01 | 2.84E-217 | 1.79E+216 | -517.26 | 1,104.52 | ge10 | interaction |
| MCWD_med:SUPERTRIBE6 | -0.722 | 254.23 | -0.003 | 0.99773 | -499.00 | 497.56 | 4.86E-01 | 1.93E-217 | 1.22E+216 | -517.26 | 1,104.52 | ge10 | interaction |
| BIO6_med:SUPERTRIBE1 | -7.684 | 155.29 | -0.049 | 0.96053 | -312.05 | 296.68 | 4.60E-04 | 3.02E-136 | 7.02E+128 | -517.26 | 1,104.52 | ge10 | interaction |
| BIO6_med:SUPERTRIBE2 | 0.157 | 155.23 | 0.001 | 0.99919 | -304.10 | 304.41 | 1.17E+00 | 8.54E-133 | 1.60E+132 | -517.26 | 1,104.52 | ge10 | interaction |
| BIO6_med:SUPERTRIBE3 | 1.666 | 155.23 | 0.011 | 0.99144 | -302.59 | 305.92 | 5.29E+00 | 3.86E-132 | 7.25E+132 | -517.26 | 1,104.52 | ge10 | interaction |
| BIO6_med:SUPERTRIBE4 | 1.649 | 155.23 | 0.011 | 0.99152 | -302.61 | 305.91 | 5.20E+00 | 3.80E-132 | 7.13E+132 | -517.26 | 1,104.52 | ge10 | interaction |
| BIO6_med:SUPERTRIBE5 | 2.468 | 155.23 | 0.016 | 0.98732 | -301.79 | 306.72 | 1.18E+01 | 8.60E-132 | 1.62E+133 | -517.26 | 1,104.52 | ge10 | interaction |
| BIO6_med:SUPERTRIBE6 | 1.565 | 155.23 | 0.010 | 0.99195 | -302.69 | 305.82 | 4.78E+00 | 3.49E-132 | 6.57E+132 | -517.26 | 1,104.52 | ge10 | interaction |
| BIO10_med:SUPERTRIBE1 | 4.036 | 231.90 | 0.017 | 0.98611 | -450.49 | 458.57 | 5.66E+01 | 2.25E-196 | 1.42E+199 | -517.26 | 1,104.52 | ge10 | interaction |
| BIO10_med:SUPERTRIBE2 | 0.514 | 231.86 | 0.002 | 0.99823 | -453.94 | 454.97 | 1.67E+00 | 7.18E-198 | 3.89E+197 | -517.26 | 1,104.52 | ge10 | interaction |
| BIO10_med:SUPERTRIBE3 | -1.462 | 231.86 | -0.006 | 0.99497 | -455.92 | 452.99 | 2.32E-01 | 9.96E-199 | 5.40E+196 | -517.26 | 1,104.52 | ge10 | interaction |
| BIO10_med:SUPERTRIBE4 | -0.827 | 231.86 | -0.004 | 0.99716 | -455.28 | 453.63 | 4.38E-01 | 1.88E-198 | 1.02E+197 | -517.26 | 1,104.52 | ge10 | interaction |
| BIO10_med:SUPERTRIBE5 | -0.492 | 231.86 | -0.002 | 0.99831 | -454.95 | 453.96 | 6.11E-01 | 2.62E-198 | 1.42E+197 | -517.26 | 1,104.52 | ge10 | interaction |
| BIO10_med:SUPERTRIBE6 | -1.431 | 231.87 | -0.006 | 0.99508 | -455.89 | 453.03 | 2.39E-01 | 1.02E-198 | 5.59E+196 | -517.26 | 1,104.52 | ge10 | interaction |
| elev_med:SUPERTRIBE1 | -1.434 | 153.10 | -0.009 | 0.99253 | -301.51 | 298.64 | 2.38E-01 | 1.14E-131 | 5.00E+129 | -517.26 | 1,104.52 | ge10 | interaction |
| elev_med:SUPERTRIBE2 | 0.072 | 153.06 | 0.000 | 0.99963 | -299.92 | 300.07 | 1.07E+00 | 5.55E-131 | 2.08E+130 | -517.26 | 1,104.52 | ge10 | interaction |
| elev_med:SUPERTRIBE3 | -0.025 | 153.06 | 0.000 | 0.99987 | -300.02 | 299.97 | 9.75E-01 | 5.04E-131 | 1.89E+130 | -517.26 | 1,104.52 | ge10 | interaction |
| elev_med:SUPERTRIBE4 | 0.063 | 153.06 | 0.000 | 0.99967 | -299.93 | 300.06 | 1.06E+00 | 5.50E-131 | 2.06E+130 | -517.26 | 1,104.52 | ge10 | interaction |
| elev_med:SUPERTRIBE5 | 0.957 | 153.06 | 0.006 | 0.99501 | -299.04 | 300.95 | 2.60E+00 | 1.35E-130 | 5.04E+130 | -517.26 | 1,104.52 | ge10 | interaction |
| elev_med:SUPERTRIBE6 | 0.173 | 153.06 | 0.001 | 0.99910 | -299.82 | 300.17 | 1.19E+00 | 6.13E-131 | 2.30E+130 | -517.26 | 1,104.52 | ge10 | interaction |

**Table S19. Supertribe-specific deviations from overall climate–woodiness slopes in ge10**
Derived deviations of supertribe-specific slopes from the overall climatic niche–woodiness relationship in the ge10 dataset. For each climatic predictor, values represent the difference between the slope estimated for a given supertribe and the corresponding overall slope across all supertribes. Positive values indicate stronger-than-average effects, whereas negative values indicate weaker-than-average effects. Slopes were derived from the interaction GLM using standardized predictors.

| SUPERTRIBE | term | slope_supertribe | overall | deviation | term_label |
|---|---|---|---|---|---|
| Aethionemeae | MCWD_med | 3.425 | -0.43812 | 3.863 | Drought (MCWD) |
| Arabodae | MCWD_med | 0.178 | -0.43812 | 0.616 | Drought (MCWD) |
| Brassicodae | MCWD_med | -0.731 | -0.43812 | -0.293 | Drought (MCWD) |
| Camelinodae | MCWD_med | -0.670 | -0.43812 | -0.232 | Drought (MCWD) |
| Heliophilodae | MCWD_med | -0.108 | -0.43812 | 0.330 | Drought (MCWD) |
| Hesperodae | MCWD_med | -0.493 | -0.43812 | -0.055 | Drought (MCWD) |
| unplaced | MCWD_med | 0.000 | -0.43812 | 0.438 | Drought (MCWD) |
| Aethionemeae | BIO6_med | -7.863 | 0.90155 | -8.764 | Frost (BIO6) |
| Arabodae | BIO6_med | -0.022 | 0.90155 | -0.923 | Frost (BIO6) |
| Brassicodae | BIO6_med | 1.487 | 0.90155 | 0.586 | Frost (BIO6) |
| Camelinodae | BIO6_med | 1.471 | 0.90155 | 0.569 | Frost (BIO6) |
| Heliophilodae | BIO6_med | 2.289 | 0.90155 | 1.387 | Frost (BIO6) |
| Hesperodae | BIO6_med | 1.387 | 0.90155 | 0.485 | Frost (BIO6) |
| unplaced | BIO6_med | 0.000 | 0.90155 | -0.902 | Frost (BIO6) |
| Aethionemeae | BIO10_med | 4.375 | -0.26592 | 4.641 | Heat (BIO10) |
| Arabodae | BIO10_med | 0.852 | -0.26592 | 1.118 | Heat (BIO10) |
| Brassicodae | BIO10_med | -1.123 | -0.26592 | -0.857 | Heat (BIO10) |
| Camelinodae | BIO10_med | -0.488 | -0.26592 | -0.222 | Heat (BIO10) |
| Heliophilodae | BIO10_med | -0.154 | -0.26592 | 0.112 | Heat (BIO10) |
| Hesperodae | BIO10_med | -1.092 | -0.26592 | -0.826 | Heat (BIO10) |
| unplaced | BIO10_med | 0.000 | -0.26592 | 0.266 | Heat (BIO10) |
| Aethionemeae | elev_med | -1.630 | 0.01771 | -1.648 | Elevation |
| Arabodae | elev_med | -0.124 | 0.01771 | -0.142 | Elevation |
| Brassicodae | elev_med | -0.221 | 0.01771 | -0.238 | Elevation |
| Camelinodae | elev_med | -0.133 | 0.01771 | -0.151 | Elevation |
| Heliophilodae | elev_med | 0.761 | 0.01771 | 0.743 | Elevation |
| Hesperodae | elev_med | -0.023 | 0.01771 | -0.041 | Elevation |
| unplaced | elev_med | 0.000 | 0.01771 | -0.018 | Elevation |

**Table S20. Input data coverage and environmental state balance across drought and frost thresholds used in BayesTraits analyses**

This table summarises data availability and binary state composition for all drought and frost thresholds used in the BayesTraits correlated-evolution analyses. For each environmental variable and threshold, we report the total number of species in the Brassicaceae Tree of Life (n_tips_tree), the number successfully mapped to the niche dataset (n_mapped_species), and the number with woodiness data (n_wood_present). Environmental data availability is given as the number of species with non-missing values (n_env_present). Thresholds for drought (Maximum Cumulative Water Deficit, MCWD) and frost (minimum temperature of the coldest month, BIO6) were defined using percentile cut-offs (p40, p50, p60) and, for frost, an additional mechanistic threshold at 0 °C. For each threshold, species were assigned to binary environmental states (env_1 vs env_0), and the counts of species in each state are reported (env_1_n, env_0_n). Across all thresholds, environmental data were available for 1,687 species, and sample sizes remained balanced between environmental states; by default at the median (p50) thresholds. This ensured adequate statistical power and comparability across threshold definitions in the correlated-evolution analyses.

| analysis | thr_label | thr_percentile | threshold_value | n_tips_tree | n_mapped_species | n_wood_present | n_env_value_present | n_env_present | env_1_n | env_0_n |
|---|---|---|---|---|---|---|---|---|---|---|
| drought | thr_p40 | 40 | -1067.1 | 3,055 | 3,055 | 3,055 | 1,687 | 1,687 | 675 | 1,012 |
| drought | thr_p50 | 50 | -885.0 | 3,055 | 3,055 | 3,055 | 1,687 | 1,687 | 844 | 843 |
| drought | thr_p60 | 60 | -700.5 | 3,055 | 3,055 | 3,055 | 1,687 | 1,687 | 1,013 | 674 |
| frost | thr_0C | NA | 0 | 3,055 | 3,055 | 3,055 | 1,687 | 1,687 | 773 | 914 |
| frost | thr_p40 | 40 | -0.1300 | 3,055 | 3,055 | 3,055 | 1,687 | 1,687 | 676 | 1,011 |
| frost | thr_p50 | 50 | 0.1100 | 3,055 | 3,055 | 3,055 | 1,687 | 1,687 | 845 | 842 |
| frost | thr_p60 | 60 | 0.3630 | 3,055 | 3,055 | 3,055 | 1,687 | 1,687 | 1,012 | 675 |

**Table S21. Bayes factor comparisons between dependent and independent BayesTraits models**

Log Bayes factor (logBF) values comparing dependent and independent models of correlated evolution between woodiness and environmental regime, computed from stepping-stone marginal likelihood estimates for each analysis (drought, frost) and threshold definition. Each row corresponds to a paired comparison of marginal likelihoods from the same chain. Positive values indicate support for the dependent model.

| analysis | thr_label | chain | dependent | independent | logBF | modelA | modelB |
|---|---|---|---|---|---|---|---|
| drought | thr_p40 | 1 | -1,739.84 | -1,746.95 | 14.212382 | dependent | independent |
| drought | thr_p40 | 2 | -1,739.30 | -1,748.93 | 19.264318 | dependent | independent |
| drought | thr_p40 | 3 | -1,742.03 | -1,747.32 | 10.595048 | dependent | independent |
| drought | thr_p40 | 4 | -1,741.92 | -1,746.63 | 9.435162 | dependent | independent |
| drought | thr_p40 | 5 | -1,739.84 | -1,747.57 | 15.448866 | dependent | independent |
| drought | thr_p40 | 6 | -1,741.41 | -1,747.00 | 11.184136 | dependent | independent |
| drought | thr_p40 | 7 | -1,743.42 | -1,748.18 | 9.506696 | dependent | independent |
| drought | thr_p40 | 8 | -1,740.25 | -1,747.54 | 14.585742 | dependent | independent |
| drought | thr_p40 | 9 | -1,742.37 | -1,747.50 | 10.273242 | dependent | independent |
| drought | thr_p40 | 10 | -1,743.68 | -1,747.80 | 8.23542 | dependent | independent |
| drought | thr_p40 | 11 | -1,742.87 | -1,747.68 | 9.623596 | dependent | independent |
| drought | thr_p40 | 12 | -1,742.49 | -1,748.32 | 11.652216 | dependent | independent |
| drought | thr_p40 | 13 | -1,740.97 | -1,747.58 | 13.231684 | dependent | independent |
| drought | thr_p40 | 14 | -1,741.10 | -1,747.29 | 12.395458 | dependent | independent |
| drought | thr_p40 | 15 | -1,740.79 | -1,747.61 | 13.654752 | dependent | independent |
| drought | thr_p40 | 16 | -1,740.66 | -1,748.24 | 15.169296 | dependent | independent |
| drought | thr_p40 | 17 | -1,742.61 | -1,746.74 | 8.263046 | dependent | independent |
| drought | thr_p40 | 18 | -1,741.73 | -1,746.69 | 9.92624 | dependent | independent |
| drought | thr_p40 | 19 | -1,743.41 | -1,747.39 | 7.941148 | dependent | independent |
| drought | thr_p40 | 20 | -1,742.64 | -1,747.34 | 9.409748 | dependent | independent |
| drought | thr_p50 | 1 | -1,733.01 | -1,741.87 | 17.718984 | dependent | independent |
| drought | thr_p50 | 2 | -1,734.01 | -1,742.91 | 17.801904 | dependent | independent |
| drought | thr_p50 | 3 | -1,735.53 | -1,742.03 | 12.992178 | dependent | independent |
| drought | thr_p50 | 4 | -1,731.84 | -1,743.33 | 22.990502 | dependent | independent |
| drought | thr_p50 | 5 | -1,733.22 | -1,741.56 | 16.677238 | dependent | independent |
| drought | thr_p50 | 6 | -1,734.37 | -1,741.87 | 14.998624 | dependent | independent |
| drought | thr_p50 | 7 | -1,734.13 | -1,742.11 | 15.949586 | dependent | independent |
| drought | thr_p50 | 8 | -1,733.69 | -1,742.31 | 17.244338 | dependent | independent |
| drought | thr_p50 | 9 | -1,733.36 | -1,742.99 | 19.270336 | dependent | independent |
| drought | thr_p50 | 10 | -1,733.00 | -1,740.53 | 15.067906 | dependent | independent |
| drought | thr_p50 | 11 | -1,734.36 | -1,741.42 | 14.116094 | dependent | independent |
| drought | thr_p50 | 12 | -1,735.35 | -1,742.18 | 13.666018 | dependent | independent |
| drought | thr_p50 | 13 | -1,732.82 | -1,741.81 | 17.981472 | dependent | independent |
| drought | thr_p50 | 14 | -1,732.12 | -1,741.61 | 18.996858 | dependent | independent |
| drought | thr_p50 | 15 | -1,734.46 | -1,742.36 | 15.800018 | dependent | independent |
| drought | thr_p50 | 16 | -1,733.94 | -1,741.87 | 15.853884 | dependent | independent |
| drought | thr_p50 | 17 | -1,734.45 | -1,742.50 | 16.098252 | dependent | independent |
| drought | thr_p50 | 18 | -1,734.46 | -1,741.48 | 14.038198 | dependent | independent |
| drought | thr_p50 | 19 | -1,733.89 | -1,742.88 | 17.961568 | dependent | independent |
| drought | thr_p50 | 20 | -1,734.78 | -1,742.34 | 15.108794 | dependent | independent |
| drought | thr_p60 | 1 | -1,738.92 | -1,755.31 | 32.768208 | dependent | independent |
| drought | thr_p60 | 2 | -1,739.48 | -1,754.92 | 30.887642 | dependent | independent |
| drought | thr_p60 | 3 | -1,738.27 | -1,753.84 | 31.130554 | dependent | independent |
| drought | thr_p60 | 4 | -1,738.36 | -1,753.97 | 31.234082 | dependent | independent |
| drought | thr_p60 | 5 | -1,739.51 | -1,753.17 | 27.323314 | dependent | independent |
| drought | thr_p60 | 6 | -1,739.06 | -1,754.86 | 31.598386 | dependent | independent |
| drought | thr_p60 | 7 | -1,738.95 | -1,754.35 | 30.791992 | dependent | independent |
| drought | thr_p60 | 8 | -1,740.52 | -1,753.21 | 25.383356 | dependent | independent |
| drought | thr_p60 | 9 | -1,739.71 | -1,754.97 | 30.520366 | dependent | independent |
| drought | thr_p60 | 10 | -1,740.68 | -1,753.13 | 24.905122 | dependent | independent |
| drought | thr_p60 | 11 | -1,740.07 | -1,753.70 | 27.246996 | dependent | independent |
| drought | thr_p60 | 12 | -1,740.40 | -1,753.48 | 26.168088 | dependent | independent |
| drought | thr_p60 | 13 | -1,738.85 | -1,753.36 | 29.018154 | dependent | independent |
| drought | thr_p60 | 14 | -1,739.11 | -1,754.33 | 30.437996 | dependent | independent |
| drought | thr_p60 | 15 | -1,739.04 | -1,754.36 | 30.647034 | dependent | independent |
| drought | thr_p60 | 16 | -1,739.74 | -1,754.60 | 29.727932 | dependent | independent |

#### Table S21 continued

| analysis | thr_label | chain | dependent | independent | logBF | modelA | modelB |
|---|---|---|---|---|---|---|---|
| drought | thr_p60 | 17 | -1,739.64 | -1,753.90 | 28.500768 | dependent | independent |
| drought | thr_p60 | 18 | -1,741.50 | -1,753.70 | 24.411408 | dependent | independent |
| drought | thr_p60 | 19 | -1,738.29 | -1,753.89 | 31.200054 | dependent | independent |
| drought | thr_p60 | 20 | -1,740.79 | -1,754.21 | 26.8447 | dependent | independent |
| frost | thr_p40 | 1 | -1,739.27 | -1,746.23 | 13.92776 | dependent | independent |
| frost | thr_p40 | 2 | -1,734.88 | -1,745.97 | 22.175886 | dependent | independent |
| frost | thr_p40 | 3 | -1,736.47 | -1,746.67 | 20.395602 | dependent | independent |
| frost | thr_p40 | 4 | -1,736.54 | -1,745.67 | 18.266114 | dependent | independent |
| frost | thr_p40 | 5 | -1,737.24 | -1,746.94 | 19.405142 | dependent | independent |
| frost | thr_p40 | 6 | -1,735.90 | -1,747.09 | 22.380116 | dependent | independent |
| frost | thr_p40 | 7 | -1,736.60 | -1,747.38 | 21.566324 | dependent | independent |
| frost | thr_p40 | 8 | -1,737.03 | -1,746.18 | 18.283632 | dependent | independent |
| frost | thr_p40 | 9 | -1,736.42 | -1,747.79 | 22.742572 | dependent | independent |
| frost | thr_p40 | 10 | -1,735.97 | -1,747.37 | 22.798226 | dependent | independent |
| frost | thr_p40 | 11 | -1,737.01 | -1,747.24 | 20.44758 | dependent | independent |
| frost | thr_p40 | 12 | -1,735.43 | -1,746.60 | 22.340754 | dependent | independent |
| frost | thr_p40 | 13 | -1,737.63 | -1,746.29 | 17.318242 | dependent | independent |
| frost | thr_p40 | 14 | -1,738.39 | -1,746.90 | 17.025108 | dependent | independent |
| frost | thr_p40 | 15 | -1,737.56 | -1,747.09 | 19.070382 | dependent | independent |
| frost | thr_p40 | 16 | -1,734.75 | -1,747.06 | 24.621676 | dependent | independent |
| frost | thr_p40 | 17 | -1,736.59 | -1,746.23 | 19.28162 | dependent | independent |
| frost | thr_p40 | 18 | -1,737.86 | -1,746.75 | 17.770174 | dependent | independent |
| frost | thr_p40 | 19 | -1,735.81 | -1,746.91 | 22.1895 | dependent | independent |
| frost | thr_p40 | 20 | -1,736.29 | -1,748.03 | 23.462232 | dependent | independent |
| frost | thr_p50 | 1 | -1,717.65 | -1,727.97 | 20.63968 | dependent | independent |
| frost | thr_p50 | 2 | -1,718.76 | -1,727.65 | 17.781154 | dependent | independent |
| frost | thr_p50 | 3 | -1,717.52 | -1,726.92 | 18.794982 | dependent | independent |
| frost | thr_p50 | 4 | -1,720.60 | -1,726.55 | 11.887424 | dependent | independent |
| frost | thr_p50 | 5 | -1,719.42 | -1,728.21 | 17.572302 | dependent | independent |
| frost | thr_p50 | 6 | -1,719.03 | -1,727.11 | 16.16202 | dependent | independent |
| frost | thr_p50 | 7 | -1,718.25 | -1,727.32 | 18.142942 | dependent | independent |
| frost | thr_p50 | 8 | -1,718.80 | -1,726.85 | 16.11034 | dependent | independent |
| frost | thr_p50 | 9 | -1,718.65 | -1,726.28 | 15.251538 | dependent | independent |
| frost | thr_p50 | 10 | -1,720.45 | -1,728.12 | 15.33921 | dependent | independent |
| frost | thr_p50 | 11 | -1,721.12 | -1,727.21 | 12.182038 | dependent | independent |
| frost | thr_p50 | 12 | -1,721.30 | -1,728.00 | 13.407054 | dependent | independent |
| frost | thr_p50 | 13 | -1,718.00 | -1,727.12 | 18.236312 | dependent | independent |
| frost | thr_p50 | 14 | -1,718.89 | -1,728.90 | 20.029958 | dependent | independent |
| frost | thr_p50 | 15 | -1,717.43 | -1,726.62 | 18.37285 | dependent | independent |
| frost | thr_p50 | 16 | -1,717.08 | -1,726.81 | 19.464548 | dependent | independent |
| frost | thr_p50 | 17 | -1,719.91 | -1,727.78 | 15.734224 | dependent | independent |
| frost | thr_p50 | 18 | -1,719.56 | -1,727.45 | 15.790164 | dependent | independent |
| frost | thr_p50 | 19 | -1,718.38 | -1,727.85 | 18.940026 | dependent | independent |
| frost | thr_p50 | 20 | -1,717.64 | -1,727.01 | 18.741902 | dependent | independent |
| frost | thr_p60 | 1 | -1,664.07 | -1,673.49 | 18.83821 | dependent | independent |
| frost | thr_p60 | 2 | -1,663.48 | -1,672.73 | 18.49435 | dependent | independent |
| frost | thr_p60 | 3 | -1,664.85 | -1,671.66 | 13.611554 | dependent | independent |
| frost | thr_p60 | 4 | -1,663.95 | -1,673.76 | 19.626278 | dependent | independent |
| frost | thr_p60 | 5 | -1,664.75 | -1,672.13 | 14.757062 | dependent | independent |
| frost | thr_p60 | 6 | -1,663.06 | -1,673.06 | 19.991084 | dependent | independent |
| frost | thr_p60 | 7 | -1,663.08 | -1,671.67 | 17.176128 | dependent | independent |
| frost | thr_p60 | 8 | -1,664.09 | -1,674.20 | 20.218942 | dependent | independent |
| frost | thr_p60 | 9 | -1,664.33 | -1,674.05 | 19.449906 | dependent | independent |
| frost | thr_p60 | 10 | -1,663.39 | -1,673.33 | 19.87612 | dependent | independent |
| frost | thr_p60 | 11 | -1,666.30 | -1,673.59 | 14.57443 | dependent | independent |
| frost | thr_p60 | 12 | -1,662.88 | -1,672.36 | 18.9501 | dependent | independent |

#### Table S21 continued

| analysis | thr_label | chain | dependent | independent | logBF | modelA | modelB |
|---|---|---|---|---|---|---|---|
| frost | thr_p60 | 13 | -1,662.99 | -1,674.25 | 22.526204 | dependent | independent |
| frost | thr_p60 | 14 | -1,661.97 | -1,672.13 | 20.32012 | dependent | independent |
| frost | thr_p60 | 15 | -1,663.75 | -1,672.59 | 17.695256 | dependent | independent |
| frost | thr_p60 | 16 | -1,664.01 | -1,673.49 | 18.975308 | dependent | independent |
| frost | thr_p60 | 17 | -1,665.01 | -1,673.36 | 16.706108 | dependent | independent |
| frost | thr_p60 | 18 | -1,662.64 | -1,672.23 | 19.171952 | dependent | independent |
| frost | thr_p60 | 19 | -1,663.59 | -1,672.51 | 17.85239 | dependent | independent |
| frost | thr_p60 | 20 | -1,665.55 | -1,672.42 | 13.737326 | dependent | independent |
| frost | thr_0C | 1 | -1,736.28 | -1,749.60 | 26.629988 | dependent | independent |
| frost | thr_0C | 2 | -1,736.95 | -1,749.60 | 25.30684 | dependent | independent |
| frost | thr_0C | 3 | -1,737.51 | -1,749.36 | 23.689594 | dependent | independent |
| frost | thr_0C | 4 | -1,734.70 | -1,751.05 | 32.69573 | dependent | independent |
| frost | thr_0C | 5 | -1,737.59 | -1,749.57 | 23.949822 | dependent | independent |
| frost | thr_0C | 6 | -1,737.18 | -1,750.65 | 26.944468 | dependent | independent |
| frost | thr_0C | 7 | -1,736.71 | -1,749.95 | 26.472058 | dependent | independent |
| frost | thr_0C | 8 | -1,738.38 | -1,749.34 | 21.926038 | dependent | independent |
| frost | thr_0C | 9 | -1,736.95 | -1,750.14 | 26.39733 | dependent | independent |
| frost | thr_0C | 10 | -1,737.95 | -1,749.43 | 22.966916 | dependent | independent |
| frost | thr_0C | 11 | -1,738.31 | -1,749.93 | 23.233912 | dependent | independent |
| frost | thr_0C | 12 | -1,737.46 | -1,749.61 | 24.311108 | dependent | independent |
| frost | thr_0C | 13 | -1,739.60 | -1,749.92 | 20.63315 | dependent | independent |
| frost | thr_0C | 14 | -1,736.01 | -1,750.51 | 29.000226 | dependent | independent |
| frost | thr_0C | 15 | -1,739.10 | -1,748.73 | 19.26857 | dependent | independent |
| frost | thr_0C | 16 | -1,737.44 | -1,751.20 | 27.529308 | dependent | independent |
| frost | thr_0C | 17 | -1,737.59 | -1,751.19 | 27.183304 | dependent | independent |
| frost | thr_0C | 18 | -1,738.19 | -1,750.59 | 24.814792 | dependent | independent |
| frost | thr_0C | 19 | -1,737.20 | -1,750.10 | 25.813784 | dependent | independent |
| frost | thr_0C | 20 | -1,736.75 | -1,749.63 | 25.75726 | dependent | independent |

**Table S22. Reversible-jump inclusion frequencies of transition rates across BayesTraits analyses**

Posterior inclusion frequencies of transition rate parameters (q) under the reversible-jump (RJ) dependent model, pooled across all chains for each analysis and threshold. Inclusion frequency represents the proportion of posterior samples in which a given rate parameter is non-zero, providing a measure of the necessity of that parameter under a parsimonious model. Values close to 1 indicate parameters consistently required to explain the data, whereas values near 0 indicate parameters frequently excluded. These frequencies reflect model structure support rather than effect size. Inclusion frequencies should be interpreted as evidence for parameter necessity rather than magnitude, and are therefore complementary to posterior summaries of rate values.

| analysis | thr_label | chain | n_iter | incl_q12 | incl_q13 | incl_q21 | incl_q24 | incl_q31 | incl_q34 | incl_q42 | incl_q43 |
|---|---|---|---|---|---|---|---|---|---|---|---|
| drought | thr_p40 | 1 | 1,801 | 1 | 1 | 1 | 1 | 0 | 1 | 1 | 1 |
| drought | thr_p40 | 2 | 1,801 | 1 | 0.529705719 | 1 | 0.529705719 | 0.470294281 | 1 | 1 | 0.828428651 |
| drought | thr_p40 | 3 | 1,801 | 1 | 0.971127152 | 1 | 0.971127152 | 0.028872848 | 1 | 1 | 1 |
| drought | thr_p40 | 4 | 1,801 | 1 | 1 | 1 | 1 | 0 | 1 | 1 | 1 |
| drought | thr_p40 | 5 | 1,801 | 1 | 1 | 1 | 1 | 0 | 1 | 1 | 1 |
| drought | thr_p40 | 6 | 1,801 | 1 | 0.134369795 | 1 | 0.134369795 | 0.865630205 | 1 | 1 | 1 |
| drought | thr_p40 | 7 | 1,801 | 1 | 1 | 1 | 1 | 0 | 1 | 1 | 1 |
| drought | thr_p40 | 8 | 1,801 | 1 | 1 | 1 | 1 | 0 | 1 | 1 | 1 |
| drought | thr_p40 | 9 | 1,801 | 1 | 0 | 1 | 0 | 1 | 1 | 1 | 0.761799001 |
| drought | thr_p40 | 10 | 1,801 | 1 | 1 | 1 | 1 | 0 | 1 | 1 | 1 |
| drought | thr_p40 | 11 | 1,801 | 1 | 0.951693504 | 1 | 0.951693504 | 0.048306496 | 1 | 0.951693504 | 1 |
| drought | thr_p40 | 12 | 1,801 | 1 | 1 | 1 | 1 | 0 | 1 | 1 | 1 |
| drought | thr_p40 | 13 | 1,801 | 1 | 0.652415325 | 1 | 0.652415325 | 0.347584675 | 1 | 1 | 1 |
| drought | thr_p40 | 14 | 1,801 | 1 | 0.737368129 | 1 | 0.737368129 | 0.262631871 | 0.997779012 | 1 | 1 |
| drought | thr_p40 | 15 | 1,801 | 1 | 0.841754581 | 1 | 0.841754581 | 0.158245419 | 1 | 1 | 1 |
| drought | thr_p40 | 16 | 1,801 | 1 | 1 | 1 | 1 | 0 | 1 | 1 | 1 |
| drought | thr_p40 | 17 | 1,801 | 1 | 1 | 1 | 1 | 0 | 1 | 1 | 1 |
| drought | thr_p40 | 18 | 1,801 | 1 | 1 | 1 | 1 | 0 | 1 | 1 | 1 |
| drought | thr_p40 | 19 | 1,801 | 1 | 0.320932815 | 1 | 0.320932815 | 0.679067185 | 1 | 1 | 1 |
| drought | thr_p40 | 20 | 1,801 | 1 | 1 | 1 | 1 | 0 | 1 | 1 | 1 |
| drought | thr_p50 | 1 | 1,801 | 0.474181011 | 1 | 0.474181011 | 1 | 0.525818989 | 0.529705719 | 0.474181011 | 0.529705719 |
| drought | thr_p50 | 2 | 1,801 | 1 | 1 | 1 | 1 | 0 | 1 | 1 | 1 |
| drought | thr_p50 | 3 | 1,801 | 1 | 0.744586341 | 1 | 0.744586341 | 0.255413659 | 1 | 1 | 0.744586341 |
| drought | thr_p50 | 4 | 1,801 | 1 | 1 | 1 | 1 | 0 | 1 | 1 | 1 |
| drought | thr_p50 | 5 | 1,801 | 0.931149361 | 0.803442532 | 0.931149361 | 0.803442532 | 0.265408107 | 0.930594114 | 0.931149361 | 0.803442532 |
| drought | thr_p50 | 6 | 1,801 | 0.8856191 | 0.1143809 | 0.8856191 | 0.1143809 | 1 | 0.8856191 | 0.8856191 | 0.1143809 |
| drought | thr_p50 | 7 | 1,801 | 1 | 0 | 1 | 0 | 1 | 0.943920044 | 1 | 0.98278734 |
| drought | thr_p50 | 8 | 1,801 | 0.767906718 | 1 | 0.767906718 | 1 | 0.232093282 | 0.789561355 | 0.767906718 | 1 |
| drought | thr_p50 | 9 | 1,801 | 1 | 0 | 1 | 0 | 1 | 0.879511383 | 1 | 0.635202665 |
| drought | thr_p50 | 10 | 1,801 | 1 | 1 | 1 | 1 | 0 | 1 | 1 | 1 |
| drought | thr_p50 | 11 | 1,801 | 1 | 1 | 1 | 1 | 0 | 1 | 1 | 1 |
| drought | thr_p50 | 12 | 1,801 | 1 | 0 | 1 | 0 | 1 | 1 | 1 | 1 |
| drought | thr_p50 | 13 | 1,801 | 1 | 0 | 1 | 0 | 1 | 0.891171571 | 1 | 0.233203776 |
| drought | thr_p50 | 14 | 1,801 | 0.720155469 | 0.955024986 | 0.720155469 | 0.955024986 | 0.324819545 | 0.720155469 | 0.720155469 | 0.955024986 |
| drought | thr_p50 | 15 | 1,801 | 1 | 0 | 1 | 0 | 1 | 1 | 1 | 0.464186563 |
| drought | thr_p50 | 16 | 1,801 | 0.922265408 | 0.901721266 | 0.922265408 | 0.901721266 | 0.176013326 | 0.922265408 | 0.922265408 | 0.922265408 |
| drought | thr_p50 | 17 | 1,801 | 0.941143809 | 0.058856191 | 0.941143809 | 0.058856191 | 1 | 0.941143809 | 0.941143809 | 0.253747918 |
| drought | thr_p50 | 18 | 1,801 | 1 | 0 | 1 | 0 | 0 | 1 | 1 | 0 |
| drought | thr_p50 | 19 | 1,801 | 1 | 0.782343143 | 1 | 0.782343143 | 0.217656857 | 1 | 1 | 0.782343143 |
| drought | thr_p50 | 20 | 1,801 | 1 | 0.397556913 | 1 | 0.397556913 | 0.602443087 | 1 | 1 | 0.397556913 |
| drought | thr_p60 | 1 | 1,801 | 0.994447529 | 1 | 1 | 1 | 0.005552471 | 0.994444444 | 0.994447529 | 1 |
| drought | thr_p60 | 2 | 1,801 | 0.926707385 | 1 | 1 | 1 | 0.073292615 | 0.970571904 | 0.926707385 | 1 |
| drought | thr_p60 | 3 | 1,801 | 1 | 0.837867851 | 0.445308162 | 0.83620211 | 0.718489728 | 0.823986674 | 0.445308162 | 0.445308162 |
| drought | thr_p60 | 4 | 1,801 | 1 | 0.838978345 | 0.295946696 | 0.838978345 | 0.865074958 | 0.870627429 | 0.295946696 | 0.179344808 |
| drought | thr_p60 | 5 | 1,801 | 1 | 0.813436979 | 0.968350916 | 0.837312604 | 0.19433648 | 0.973903387 | 0.968350916 | 0.968350916 |
| drought | thr_p60 | 6 | 1,801 | 0.013881177 | 0.013881177 | 0.013881177 | 1 | 0.013002364 | 0.986118823 | 0.986118823 | 1 |
| drought | thr_p60 | 7 | 1,801 | 0.925596891 | 0.303720155 | 0.997779012 | 0.303720155 | 0.770682954 | 0.898945031 | 0.925596891 | 0.997779012 |
| drought | thr_p60 | 8 | 1,801 | 0.663520267 | 0.835091616 | 1 | 0.835091616 | 0.501388118 | 0.865630205 | 0.851193781 | 1 |
| drought | thr_p60 | 9 | 1,801 | 1 | 1 | 1 | 1 | 0 | 1 | 1 | 1 |
| drought | thr_p60 | 10 | 1,801 | 0 | 1 | 1 | 1 | 1 | 0.445308162 | 0.611882288 | 1 |
| drought | thr_p60 | 11 | 1,801 | 1 | 1 | 1 | 1 | 0 | 1 | 1 | 1 |
| drought | thr_p60 | 12 | 1,801 | 0.902276513 | 1 | 1 | 1 | 0.097723487 | 0.978333333 | 1 | 1 |
| drought | thr_p60 | 13 | 1,801 | 1 | 0.128262077 | 1 | 0.128262077 | 0.871737923 | 0.952248751 | 1 | 1 |
| drought | thr_p60 | 14 | 1,801 | 0.921710161 | 1 | 1 | 1 | 0.078289839 | 0.982768205 | 1 | 1 |
| drought | thr_p60 | 15 | 1,801 | 0.909494725 | 0.339811216 | 0.968350916 | 0.339811216 | 0.782343143 | 0.919489173 | 0.968350916 | 0.968350916 |
| drought | thr_p60 | 16 | 1,801 | 1 | 1 | 0.867851194 | 1 | 0.132148806 | 1 | 0.867851194 | 0.867851194 |
| drought | thr_p60 | 17 | 1,801 | 1 | 1 | 1 | 1 | 0 | 1 | 1 | 1 |
| drought | thr_p60 | 18 | 1,801 | 1 | 0.476957246 | 0.915047196 | 0.476957246 | 0.607995558 | 0.921710161 | 0.915047196 | 0.915047196 |
| drought | thr_p60 | 19 | 1,801 | 1 | 1 | 0.141588007 | 1 | 0.858411993 | 0.735555556 | 0.141588007 | 0.141588007 |
| drought | thr_p60 | 20 | 1,801 | 0.919489173 | 1 | 0.820655192 | 1 | 0.259855636 | 0.919444444 | 0.820655192 | 1 |
| frost | thr_p40 | 1 | 1,801 | 0.85896724 | 0.85896724 | 0.564686285 | 0.85896724 | 0.629650194 | 0.85896724 | 0.576346474 | 0.618888889 |
| frost | thr_p40 | 2 | 1,801 | 1 | 1 | 0.913381455 | 1 | 0.913381455 | 0.913381455 | 0.913888889 | 0.086618545 |
| frost | thr_p40 | 3 | 1,801 | 1 | 1 | 1 | 1 | 1 | 1 | 0 | 1 |
| frost | thr_p40 | 4 | 1,801 | 0.891171571 | 0.891171571 | 0.826207662 | 0.780122154 | 0.826207662 | 0.780122154 | 0.296501943 | 0.861398176 |
| frost | thr_p40 | 5 | 1,801 | 0.011104942 | 0.011104942 | 1 | 0.368167203 | 1 | 0.127151582 | 0.988895058 | 0.924898902 |
| frost | thr_p40 | 6 | 1,801 | 0.992781788 | 0.992781788 | 0.753470294 | 0.992781788 | 0.753470294 | 0.992781788 | 0.253747918 | 0.766111111 |
| frost | thr_p40 | 7 | 1,801 | 0.915602443 | 0.915602443 | 0.901166019 | 0.915602443 | 0.901166019 | 0.915602443 | 0.152137701 | 0.913434514 |
| frost | thr_p40 | 8 | 1,801 | 0.979455858 | 0.979455858 | 0.887284842 | 0.992781788 | 0.887284842 | 0.992781788 | 0.12770683 | 0.902575588 |
| frost | thr_p40 | 9 | 1,801 | 0.970571904 | 0.970571904 | 0.858411993 | 0.970571904 | 0.858411993 | 0.970571904 | 0.171016102 | 0.863964464 |
| frost | thr_p40 | 10 | 1,801 | 1 | 1 | 1 | 1 | 1 | 1 | 0 | 1 |
| frost | thr_p40 | 11 | 1,801 | 0.935036091 | 0.935036091 | 0.707384786 | 0.935036091 | 0.707384786 | 0.935036091 | 0.357579123 | 0.727373681 |
| frost | thr_p40 | 12 | 1,801 | 0.634092171 | 0.634092171 | 0.744586341 | 0.634092171 | 0.744586341 | 0.634092171 | 0.58467518 | 0.775084555 |
| frost | thr_p40 | 13 | 1,801 | 0.980011105 | 0.980011105 | 0.8667407 | 0.938922821 | 0.8667407 | 0.932259856 | 0.145474736 | 0.897185526 |
| frost | thr_p40 | 14 | 1,801 | 0.934480844 | 0.934480844 | 0.95613548 | 0.621876735 | 0.95613548 | 0.934480844 | 0.421987785 | 0.952672606 |
| frost | thr_p40 | 15 | 1,801 | 0.96946141 | 0.96946141 | 0.741810105 | 0.96946141 | 0.741810105 | 0.96946141 | 0.288728484 | 0.759022765 |
| frost | thr_p40 | 16 | 1,801 | 0.868406441 | 0.868406441 | 0.309827873 | 0.868406441 | 0.444752915 | 0.868406441 | 0.821765686 | 0.377777778 |
| frost | thr_p40 | 17 | 1,801 | 0.576901721 | 0.576901721 | 1 | 0.991116047 | 1 | 0.991116047 | 1 | 0.181935484 |
| frost | thr_p40 | 18 | 1,801 | 0.673514714 | 0.673514714 | 0.85896724 | 0.673514714 | 0.85896724 | 0.673514714 | 0.467518046 | 0.843888889 |
| frost | thr_p40 | 19 | 1,801 | 0.936146585 | 0.936146585 | 0.780677401 | 0.936146585 | 0.780677401 | 0.936146585 | 0.283176013 | 0.790116602 |
| frost | thr_p40 | 20 | 1,801 | 0.887840089 | 0.887840089 | 1 | NA | 1 | 1 | 1 | 0.112159911 |
| frost | thr_p50 | 1 | 1,801 | 1 | 1 | 1 | 1 | 1 | 1 | 0 | 1 |
| frost | thr_p50 | 2 | 1,801 | 1 | 1 | 0.96946141 | 1 | 0.96946141 | 1 | 0.03053859 | 0.97832129 |
| frost | thr_p50 | 3 | 1,801 | 1 | 1 | 0.894503054 | 1 | 0.894503054 | 1 | 0.105496946 | 0.916111111 |
| frost | thr_p50 | 4 | 1,801 | 1 | 1 | 1 | 1 | 1 | 1 | 0 | 1 |
| frost | thr_p50 | 5 | 1,801 | 1 | 1 | 1 | 1 | 1 | 1 | 0 | 1 |
| frost | thr_p50 | 6 | 1,801 | 0.980566352 | 0.980566352 | 0.678511938 | 0.980566352 | 0.678511938 | 0.980566352 | 0.34092171 | 0.7 |
| frost | thr_p50 | 7 | 1,801 | 1 | 1 | 0.81898945 | 1 | 0.81898945 | 1 | 0.18101055 | 0.848804892 |
| frost | thr_p50 | 8 | 1,801 | 1 | 1 | 0.330927263 | 1 | 0.330927263 | 1 | 0.669072737 | 0.388117712 |
| frost | thr_p50 | 9 | 1,801 | 0.961687951 | 0.961687951 | 0.575791227 | 0.961687951 | 0.575791227 | 0.961687951 | 0.462520822 | 0.646436526 |
| frost | thr_p50 | 10 | 1,801 | 0.973903387 | 0.973903387 | 0.479733481 | 0.973903387 | 0.479733481 | 0.973903387 | 0.546363132 | 0.546970539 |
| frost | thr_p50 | 11 | 1,801 | 0.97723487 | 0.97723487 | 0.621876735 | 0.97723487 | 0.621876735 | 0.97723487 | 0.400888395 | 0.65 |
| frost | thr_p50 | 12 | 1,801 | 0.9905608 | 0.9905608 | 0.88950583 | 0.9905608 | 0.88950583 | 0.9905608 | 0.11993337 | 0.889444444 |
| frost | thr_p50 | 13 | 1,801 | 1 | 1 | 0.961132704 | 1 | 0.961132704 | 1 | 0.038867296 | 0.961089494 |
| frost | thr_p50 | 14 | 1,801 | 1 | 1 | 0.397001666 | 1 | 0.397001666 | 1 | 0.602998334 | 0.46 |
| frost | thr_p50 | 15 | 1,801 | 1 | 1 | 1 | 1 | 1 | 1 | 0 | 1 |
| frost | thr_p50 | 16 | 1,801 | 0.996668517 | 0.996668517 | 0.953359245 | 0.996668517 | 0.953359245 | 0.996668517 | 0.049972238 | 0.956111111 |
| frost | thr_p50 | 17 | 1,801 | 1 | 1 | 1 | 1 | 1 | 1 | 0 | 0.999444753 |
| frost | thr_p50 | 18 | 1,801 | 1 | 1 | 0 | 1 | 0 | 0 | 1 | 0.122846026 |
| frost | thr_p50 | 19 | 1,801 | 0.999444753 | 0.999444753 | 0.316490838 | 0.999444753 | 0.316490838 | 0.999444753 | 0.684064409 | 0.396331295 |
| frost | thr_p50 | 20 | 1,801 | 1 | 1 | 1 | 1 | 1 | 1 | 0 | 1 |
| frost | thr_p60 | 1 | 1,801 | 1 | 1 | 1 | 1 | 1 | 1 | 0 | 1 |
| frost | thr_p60 | 2 | 1,801 | 0.770682954 | 0.999444753 | 0.770682954 | 1 | 0.770682954 | 0.997740113 | 0.229317046 | 0.770682954 |
| frost | thr_p60 | 3 | 1,801 | 0.611882288 | 0.998889506 | 0.611882288 | 0.999444753 | 0.611882288 | 0.997747748 | 0.388672959 | 0.611327041 |
| frost | thr_p60 | 4 | 1,801 | 1 | 1 | 1 | 1 | 1 | 1 | 0 | 1 |
| frost | thr_p60 | 5 | 1,801 | 1 | 1 | 1 | 1 | 1 | 1 | 0 | 1 |
| frost | thr_p60 | 6 | 1,801 | 1 | 1 | 1 | 1 | 1 | 1 | 0 | 1 |
| frost | thr_p60 | 7 | 1,801 | 1 | 1 | 1 | 1 | 1 | 1 | 0 | 1 |
| frost | thr_p60 | 8 | 1,801 | 1 | 1 | 1 | 1 | 1 | 1 | 0 | 1 |
| frost | thr_p60 | 9 | 1,801 | 1 | 1 | 1 | 1 | 1 | 1 | 0 | 1 |
| frost | thr_p60 | 10 | 1,801 | 1 | 1 | 1 | 1 | 1 | 1 | 0 | 1 |
| frost | thr_p60 | 11 | 1,801 | 1 | 1 | 1 | 1 | 1 | 1 | 0 | 1 |
| frost | thr_p60 | 12 | 1,801 | 1 | 1 | 1 | 1 | 1 | 1 | 0 | 1 |
| frost | thr_p60 | 13 | 1,801 | 1 | 1 | 1 | 1 | 1 | 1 | 0 | 1 |
| frost | thr_p60 | 14 | 1,801 | 0.92948362 | 1 | 0.92948362 | 1 | 0.92948362 | 1 | 0.07051638 | 1 |
| frost | thr_p60 | 15 | 1,801 | 1 | 1 | 1 | 1 | 1 | 1 | 0 | 1 |
| frost | thr_p60 | 16 | 1,801 | 0.742365353 | 0.987784564 | 0.742365353 | 0.987784564 | 0.742365353 | 0.987688864 | 0.269850083 | 0.858411993 |
| frost | thr_p60 | 17 | 1,801 | 1 | 1 | 1 | 1 | 1 | 1 | 0 | 1 |
| frost | thr_p60 | 18 | 1,801 | 1 | 1 | 1 | 1 | 1 | 1 | 0 | 1 |
| frost | thr_p60 | 19 | 1,801 | 1 | 1 | 1 | 1 | 1 | 1 | 0 | 1 |
| frost | thr_p60 | 20 | 1,801 | 1 | 1 | 1 | 1 | 1 | 1 | 0 | 1 |
| frost | thr_0C | 1 | 1,801 | 0.984453082 | 0.984453082 | 0.753470294 | 0.984453082 | 0.753470294 | 0.984453082 | 0.262631871 | 0.765555556 |
| frost | thr_0C | 2 | 1,801 | 1 | 1 | 1 | 1 | 1 | 1 | 0 | 1 |
| frost | thr_0C | 3 | 1,801 | 0.975569128 | 0.975569128 | 1 | 0.975569128 | 1 | 0.975569128 | 0.024430872 | 0.983888889 |
| frost | thr_0C | 4 | 1,801 | 1 | 1 | 1 | 1 | 1 | 1 | 0 | 1 |
| frost | thr_0C | 5 | 1,801 | 1 | 1 | 1 | 1 | 1 | 1 | 0 | 1 |
| frost | thr_0C | 6 | 1,801 | 1 | 1 | 0.905052748 | 1 | 0.905052748 | 1 | 0.094947252 | 0.907222222 |
| frost | thr_0C | 7 | 1,801 | 1 | 1 | 0.870627429 | 1 | 0.870627429 | 1 | 0.129372571 | 0.900610772 |
| frost | thr_0C | 8 | 1,801 | 1 | 1 | 1 | 1 | 1 | 1 | 0 | 1 |
| frost | thr_0C | 9 | 1,801 | 1 | 1 | 1 | 1 | 1 | 1 | 0 | 1 |
| frost | thr_0C | 10 | 1,801 | 1 | 1 | 1 | 1 | 1 | 1 | 0 | 1 |
| frost | thr_0C | 11 | 1,801 | 1 | 1 | 0.519156024 | 1 | 1 | 1 | 0.480843976 | 0.606111111 |
| frost | thr_0C | 12 | 1,801 | 1 | 1 | 0.543031649 | 1 | 0.543031649 | 1 | 0.456968351 | 0.584119933 |
| frost | thr_0C | 13 | 1,801 | 1 | 1 | 1 | 1 | 1 | 1 | 0 | 1 |
| frost | thr_0C | 14 | 1,801 | 1 | 1 | 1 | 1 | 1 | 1 | 0 | 1 |
| frost | thr_0C | 15 | 1,801 | 1 | 1 | 0.916712937 | 1 | 0.916712937 | 1 | 0.083287063 | 0.936146585 |
| frost | thr_0C | 16 | 1,801 | 1 | 1 | 0.675735702 | 1 | 0.68184342 | 1 | 0.324264298 | 0.700166574 |
| frost | thr_0C | 17 | 1,801 | 0.978345364 | 0.978345364 | 1 | 0.978345364 | 1 | 0.978345364 | 0.021654636 | 0.991662034 |
| frost | thr_0C | 18 | 1,801 | 0.937257079 | 0.936146585 | 0.664075514 | 0.998717126 | 0.664075514 | 1 | 0.471404775 | 0.627222222 |
| frost | thr_0C | 19 | 1,801 | 0.872848418 | 0.872848418 | 1 | 0.872848418 | 1 | 0.872848418 | 0.127151582 | 0.986666667 |
| frost | thr_0C | 20 | 1,801 | 0.993892282 | 0.993892282 | 0.921710161 | 0.993892282 | 0.921710161 | 0.993892282 | 0.074403109 | 0.927222222 |

**Table S23. Posterior summaries of transition rate parameters (q) across BayesTraits analyses, thresholds, and models**

Posterior summaries of transition rate parameters (q) across BayesTraits analyses, thresholds, and models. For each environmental variable (drought and frost), threshold, model (dependent and reversible-jump dependent), and transition-rate parameter (q12–q43), the table reports posterior inclusion frequency, posterior median, and the bounds of the 95% credible interval (lo and hi).

| analysis | thr_label | model | param | inclusion | median | lo | hi |
|---|---|---|---|---|---|---|---|
| drought | thr_p40 | dependent | q12 | 1 | 0.056579 | 0.04441695 | 0.0711332 |
| drought | thr_p40 | dependent | q13 | 1 | 0.016877 | 0.011263 | 0.0242291 |
| drought | thr_p40 | dependent | q21 | 1 | 0.085761 | 0.071725 | 0.10263605 |
| drought | thr_p40 | dependent | q24 | 1 | 0.021818 | 0.015905 | 0.028993525 |
| drought | thr_p40 | dependent | q31 | 1 | 0.290702 | 0.207804425 | 0.402422825 |
| drought | thr_p40 | dependent | q34 | 1 | 0.0532435 | 0.017164025 | 0.110484925 |
| drought | thr_p40 | dependent | q42 | 1 | 0.052352 | 0.031195 | 0.077991525 |
| drought | thr_p40 | dependent | q43 | 1 | 0.0542315 | 0.03053785 | 0.080910575 |
| drought | thr_p40 | dependent_RJ | q12 | 1 | 0.063939 | 0.054708325 | 0.077900725 |
| drought | thr_p40 | dependent_RJ | q13 | 1 | 0.020306 | 0.016547 | 0.024728 |
| drought | thr_p40 | dependent_RJ | q21 | 1 | 0.064068 | 0.055214475 | 0.079192 |
| drought | thr_p40 | dependent_RJ | q24 | 1 | 0.020306 | 0.016547 | 0.024728 |
| drought | thr_p40 | dependent_RJ | q31 | 1 | 0.3149775 | 0.2249945 | 0.43607925 |
| drought | thr_p40 | dependent_RJ | q34 | 1 | 0.0633 | 0.0197229 | 0.0742622 |
| drought | thr_p40 | dependent_RJ | q42 | 1 | 0.062993 | 0.019198475 | 0.0731051 |
| drought | thr_p40 | dependent_RJ | q43 | 1 | 0.063778 | 0.0506777 | 0.078004 |
| drought | thr_p50 | dependent | q12 | 1 | 0.0710185 | 0.057038075 | 0.0874531 |
| drought | thr_p50 | dependent | q13 | 1 | 0.018479 | 0.011717 | 0.02740005 |
| drought | thr_p50 | dependent | q21 | 1 | 0.05591 | 0.04487995 | 0.06872005 |
| drought | thr_p50 | dependent | q24 | 1 | 0.020221 | 0.015101 | 0.0262901 |
| drought | thr_p50 | dependent | q31 | 1 | 0.3689715 | 0.2609662 | 0.515050025 |
| drought | thr_p50 | dependent | q34 | 1 | 0.1071855 | 0.048376975 | 0.200784325 |
| drought | thr_p50 | dependent | q42 | 1 | 0.062368 | 0.039945275 | 0.08750945 |
| drought | thr_p50 | dependent | q43 | 1 | 0.037528 | 0.016606425 | 0.06272605 |
| drought | thr_p50 | dependent_RJ | q12 | 1 | 0.0616825 | 0.05134795 | 0.073368625 |
| drought | thr_p50 | dependent_RJ | q13 | 1 | 0.02 | 0.015174 | 0.024514525 |
| drought | thr_p50 | dependent_RJ | q21 | 1 | 0.0616825 | 0.05134795 | 0.073368625 |
| drought | thr_p50 | dependent_RJ | q24 | 1 | 0.02 | 0.015174 | 0.024514525 |
| drought | thr_p50 | dependent_RJ | q31 | 1 | 0.3746935 | 0.015174 | 0.5427956 |
| drought | thr_p50 | dependent_RJ | q34 | 1 | 0.0614835 | 0.0260609 | 0.073282675 |
| drought | thr_p50 | dependent_RJ | q42 | 1 | 0.0616815 | 0.051340475 | 0.073368625 |
| drought | thr_p50 | dependent_RJ | q43 | 1 | 0.021684 | 0.015174 | 0.063917525 |
| drought | thr_p60 | dependent | q12 | 1 | 0.110303 | 0.090294425 | 0.134118575 |
| drought | thr_p60 | dependent | q13 | 1 | 0.016051 | 0.008161475 | 0.028136525 |
| drought | thr_p60 | dependent | q21 | 1 | 0.049397 | 0.039143325 | 0.062009525 |
| drought | thr_p60 | dependent | q24 | 1 | 0.021726 | 0.017073425 | 0.02712315 |
| drought | thr_p60 | dependent | q31 | 1 | 0.5860845 | 0.3675601 | 0.94704195 |
| drought | thr_p60 | dependent | q34 | 1 | 0.140242 | 0.033723025 | 0.31936845 |
| drought | thr_p60 | dependent | q42 | 1 | 0.0643625 | 0.042012475 | 0.089915525 |
| drought | thr_p60 | dependent | q43 | 1 | 0.0414365 | 0.021782425 | 0.067588525 |
| drought | thr_p60 | dependent_RJ | q12 | 1 | 0.10761 | 0.04923665 | 0.13349015 |
| drought | thr_p60 | dependent_RJ | q13 | 1 | 0.022003 | 0.017260475 | 0.049403025 |
| drought | thr_p60 | dependent_RJ | q21 | 1 | 0.04847 | 0.02642185 | 0.05977305 |
| drought | thr_p60 | dependent_RJ | q24 | 1 | 0.021961 | 0.01724295 | 6.18747605 |
| drought | thr_p60 | dependent_RJ | q31 | 1 | 0.706831 | 0 | 1.091542175 |
| drought | thr_p60 | dependent_RJ | q34 | 1 | 0.089984 | 0.021290475 | 28.99380515 |
| drought | thr_p60 | dependent_RJ | q42 | 1 | 0.052946 | 0.042498175 | 28.99380515 |
| drought | thr_p60 | dependent_RJ | q43 | 1 | 0.0479875 | 0.020371475 | 6.186312 |
| frost | thr_p40 | dependent | q12 | 1 | 0.0357335 | 0.025783475 | 0.048387775 |
| frost | thr_p40 | dependent | q13 | 1 | 0.024749 | 0.019751425 | 0.030810525 |
| frost | thr_p40 | dependent | q21 | 1 | 0.111226 | 0.093195375 | 0.131858525 |
| frost | thr_p40 | dependent | q24 | 1 | 0.009439 | 0.00445 | 0.01724705 |
| frost | thr_p40 | dependent | q31 | 1 | 0.070672 | 0.04723995 | 0.099660775 |
| frost | thr_p40 | dependent | q34 | 1 | 0.0333695 | 0.01736495 | 0.05538105 |
| frost | thr_p40 | dependent | q42 | 1 | 0.483165 | 0.28957485 | 0.72700615 |
| frost | thr_p40 | dependent | q43 | 1 | 0.1400515 | 0.037552375 | 0.3405756 |
| frost | thr_p40 | dependent_RJ | q12 | 1 | 0.025258 | 0.02051595 | 0.038713575 |
| frost | thr_p40 | dependent_RJ | q13 | 1 | 0.025131 | 0.020238475 | 0.037295525 |
| frost | thr_p40 | dependent_RJ | q21 | 1 | 0.097029 | 0.083908475 | 0.120574525 |
| frost | thr_p40 | dependent_RJ | q24 | 1 | 0.0234875 | 0 | 0.028685 |
| frost | thr_p40 | dependent_RJ | q31 | 1 | 0.096628 | 0.079842025 | 0.11885825 |
| frost | thr_p40 | dependent_RJ | q34 | 1 | 0.024899 | 0.006866475 | 0.1157531 |
| frost | thr_p40 | dependent_RJ | q42 | 1 | 0.584563 | 0.0979709 | 0.886885675 |
| frost | thr_p40 | dependent_RJ | q43 | 1 | 0.0954985 | 0.006836275 | 0.783678925 |
| frost | thr_p50 | dependent | q12 | 1 | 0.036184 | 0.025424425 | 0.048614 |
| frost | thr_p50 | dependent | q13 | 1 | 0.0249415 | 0.01949695 | 0.031671725 |
| frost | thr_p50 | dependent | q21 | 1 | 0.0673615 | 0.056180325 | 0.08051405 |
| frost | thr_p50 | dependent | q24 | 1 | 0.01167 | 0.00736995 | 0.017785525 |
| frost | thr_p50 | dependent | q31 | 1 | 0.064868 | 0.04100495 | 0.092362575 |
| frost | thr_p50 | dependent | q34 | 1 | 0.0268285 | 0.008667475 | 0.052790575 |
| frost | thr_p50 | dependent | q42 | 1 | 0.286266 | 0.1952667 | 0.499111725 |
| frost | thr_p50 | dependent | q43 | 1 | 0.115334 | 0.04785365 | 0.23977835 |
| frost | thr_p50 | dependent_RJ | q12 | 1 | 0.023129 | 0.018641475 | 0.03035 |
| frost | thr_p50 | dependent_RJ | q13 | 1 | 0.023121 | 0.018628475 | 0.030269525 |
| frost | thr_p50 | dependent_RJ | q21 | 1 | 0.066141 | 0.056699475 | 0.07633405 |
| frost | thr_p50 | dependent_RJ | q24 | 1 | 0.023111 | 0.018592475 | 0.030260525 |
| frost | thr_p50 | dependent_RJ | q31 | 1 | 0.066139 | 0.056673475 | 0.076331575 |
| frost | thr_p50 | dependent_RJ | q34 | 1 | 0.023497 | 0.018634 | 0.0673013 |
| frost | thr_p50 | dependent_RJ | q42 | 1 | 0.393066 | 0.245622925 | 0.78951735 |
| frost | thr_p50 | dependent_RJ | q43 | 1 | 0.0652515 | 0.0247917 | 0.075910575 |
| frost | thr_p60 | dependent | q12 | 1 | 0.0444865 | 0.032947475 | 0.058137 |
| frost | thr_p60 | dependent | q13 | 1 | 0.026637 | 0.020316 | 0.034585575 |
| frost | thr_p60 | dependent | q21 | 1 | 0.049413 | 0.040760425 | 0.0594291 |
| frost | thr_p60 | dependent | q24 | 1 | 0.012367 | 0.007933 | 0.017954525 |
| frost | thr_p60 | dependent | q31 | 1 | 0.0624795 | 0.0389708 | 0.092005725 |
| frost | thr_p60 | dependent | q34 | 1 | 0.014853 | 0.004010475 | 0.035241625 |
| frost | thr_p60 | dependent | q42 | 1 | 0.216864 | 0.160099075 | 0.2866906 |
| frost | thr_p60 | dependent | q43 | 1 | 0.04056 | 0.01525795 | 0.085164825 |
| frost | thr_p60 | dependent_RJ | q12 | 1 | 0.0482415 | 0.040803425 | 0.056716525 |
| frost | thr_p60 | dependent_RJ | q13 | 1 | 0.019191 | 0.015775 | 0.02326405 |
| frost | thr_p60 | dependent_RJ | q21 | 1 | 0.048244 | 0.04083295 | 0.056716525 |
| frost | thr_p60 | dependent_RJ | q24 | 1 | 0.019164 | 0.015621475 | 0.023047525 |
| frost | thr_p60 | dependent_RJ | q31 | 1 | 0.048207 | 0.0400539 | 0.05671005 |
| frost | thr_p60 | dependent_RJ | q34 | 1 | 0.019158 | 0.015246475 | 0.023387 |
| frost | thr_p60 | dependent_RJ | q42 | 1 | 0.248676 | 0.1904629 | 0.318993775 |
| frost | thr_p60 | dependent_RJ | q43 | 1 | 0.043061 | 0.016831 | 0.055084525 |
| frost | thr_0C | dependent | q12 | 1 | 0.039614 | 0.028551 | 0.05288805 |
| frost | thr_0C | dependent | q13 | 1 | 0.025475 | 0.019946 | 0.03230805 |
| frost | thr_0C | dependent | q21 | 1 | 0.0856275 | 0.07109095 | 0.103018525 |
| frost | thr_0C | dependent | q24 | 1 | 0.012081 | 0.006505 | 0.02026915 |
| frost | thr_0C | dependent | q31 | 1 | 0.0666905 | 0.04218555 | 0.093818675 |
| frost | thr_0C | dependent | q34 | 1 | 0.0385935 | 0.022132 | 0.060972675 |
| frost | thr_0C | dependent | q42 | 1 | 0.47381 | 0.2941399 | 0.715333675 |
| frost | thr_0C | dependent | q43 | 1 | 0.143762 | 0.04263285 | 0.329610625 |
| frost | thr_0C | dependent_RJ | q12 | 1 | 0.026336 | 0.021865475 | 0.0423323 |
| frost | thr_0C | dependent_RJ | q13 | 1 | 0.026105 | 0.02110295 | 0.031472625 |
| frost | thr_0C | dependent_RJ | q21 | 1 | 0.0771875 | 0.0669239 | 0.091766775 |
| frost | thr_0C | dependent_RJ | q24 | 1 | 0.026056 | 0.020608425 | 0.03107405 |
| frost | thr_0C | dependent_RJ | q31 | 1 | 0.0764565 | 0.03970485 | 0.088396575 |
| frost | thr_0C | dependent_RJ | q34 | 1 | 0.026658 | 0.022187475 | 0.07510305 |
| frost | thr_0C | dependent_RJ | q42 | 1 | 0.643269 | 0.4218604 | 0.905391525 |
| frost | thr_0C | dependent_RJ | q43 | 1 | 0.076124 | 0.025687225 | 0.09135035 |

#### *Supplementary References*

1. L. A. Nikolov, P. Shushkov, B. Nevado, X. Gan, I. A. Al-Shehbaz, D. Filatov, C. D. Bailey, M. Tsiantis, Resolving the backbone of the Brassicaceae phylogeny for investigating trait diversity. *New Phytol.* **222**, 1638–1651 (2019).

2. K. P. Hendriks, C. Kiefer, I. A. Al-Shehbaz, C. D. Bailey, A. Hooft van Huysduynen, L. A. Nikolov, L. Nauheimer, A. R. Zuntini, D. A. German, A. Franzke, M. A. Koch, M. A. Lysak, Ó. Toro-Núñez, B. Özüdoğru, V. R. Invernón, N. Walden, O. Maurin, N. M. Hay, P. Shushkov, T. Mandáková, M. E. Schranz, M. Thulin, M. D. Windham, I. Rešetnik, S. Španiel, E. Ly, J. C. Pires, A. Harkess, B. Neuffer, R. Vogt, C. Bräuchler, H. Rainer, S. B. Janssens, M. Schmull, A. Forrest, A. Guggisberg, S. Zmarzty, B. J. Lepschi, N. Scarlett, F. W. Stauffer, I. Schönberger, P. Heenan, W. J. Baker, F. Forest, K. Mummenhoff, F. Lens, Global Brassicaceae phylogeny based on filtering of 1,000-gene dataset. *Curr. Biol.* **33**, 4052-4068.e6 (2023).
